# A tRNA Modification-based strategy for Identifying amiNo acid Overproducers (AMINO)

**DOI:** 10.1101/2022.11.21.517450

**Authors:** Hao Guo, Xiaoyan Ma, Ning Wang, Tingting Ding, Bo Zheng, Liwei Guo, Chaoyong Huang, Wuyuan Zhang, Lichao Sun, Yi-Xin Huo

**Author notes:** Correspondence and requests for materials should be addressed to Y.-X.H. These authors contributed equally (Hao Guo, Xiaoyan Ma).

## Abstract

Amino acids have a multi-billion-dollar market with rising demand, prompting the development of high-performance microbial factories. However, a general screening strategy applicable to all proteinogenic and non-proteinogenic amino acids is still lacking. Modification of the critical structure of tRNA could decrease the aminoacylation level of tRNA catalyzed by aminoacyl-tRNA synthetases. Involved in a two-substrate sequential reaction, amino acids with increased concentration could elevate the reduced aminoacylation rate caused by specific tRNA modification. Here, we developed a selection system for overproducers of specific amino acids using corresponding engineered tRNAs and reporter genes. As a proof-of-concept, overproducers of five amino acids such as L-tryptophan were screened out by growth-based and/or fluorescence-activated cell sorting (FACS)-based screening from random mutation libraries of *Escherichia coli* and *Corynebacterium glutamicum*, respectively. This study provided a universal strategy that could be applied to screen overproducers of proteinogenic and non-proteinogenic amino acids in amber-stop-codon-recoded or non-recoded hosts.

## Introduction

The amino acids have been widely applied in the food, animal feed, pharmaceutical, and cosmetic industries ^1^. The worldwide market value of amino acids is expected to reach $ 29.6 billion in 2026 with a compound annual growth rate (CAGR) of 6∼8% ^2^. Among these amino acids, L-tryptophan has the highest CAGR of 13.8%, projecting to reach $ 2.37 billion by the end of 2026 ^3^. Nevertheless, the current production of amino acid fermentation cannot satisfy the rapidly expanding demand for most Lamino acids (Fig. 1a) ^4-6^. Although rational metabolic engineering strategies have contributed to the high titers, a limited understanding of host strains and the lack of beneficial mutations have hindered the further improvement of amino acid production. Traditionally, amino acid overproducers can be screened out from random mutant libraries by using amino acid analogs. A toxic amino acid analog, which has a similar size, structure, and charge properties as the proteinogenic amino acids, would compete with its corresponding amino acid for the finite tRNAs in the process of protein biosynthesis ^7,8^. Once inserted into any polypeptide, the analog could impede the synthesis or function of that polypeptide, resulting in growth retardation or even cell death ^9,10^. Cells overproducing an amino acid could produce functional proteins to survive the stresses from the analog of the amino acid ^11^, forming the basis of the analog-dependent strategy for screening amino acid overproducers. For instance, L-leucine overproduction strains were selected using a high concentration of 4azaleucine ^12,13^. However, an analog not only disrupts protein synthesis but also interferes with cellular activities ^11,14,15^. Mutants with enhanced amino acid production may not survive these adverse effects. Furthermore, only a limited number of appropriate analogs have been discovered for screening amino acid overproducers.

**Fig. 1.**
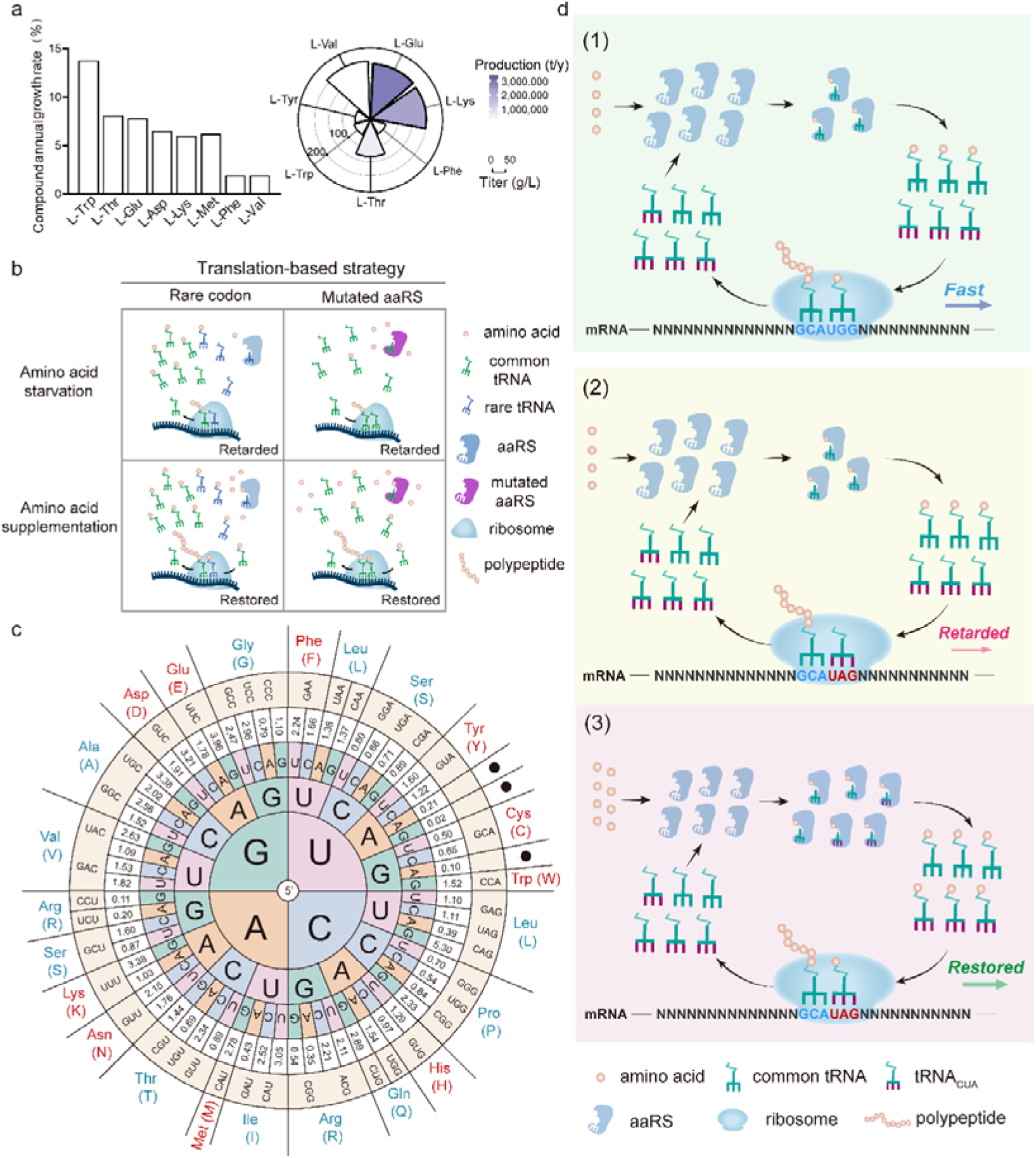
Amino acid productions and schematic diagram of developing the AMINO strategy. **a**. CAGR of amino acids market size (left), the annual productions (right, represented by color intensity), and the fermentation titers (right, represented by bar height) for seven selected amino acids. **b**. Schematic diagram of the rare codon-based selection system and the mutant aaRS-based selection system for screening amino acid overproducers. A rare codon, that often corresponds to a rare tRNA isoacceptor, is translated to its corresponding amino acid when the synonymous common isoacceptors are preferentially charged. Increased intracellular concentrations of the corresponding amino acid could rescue the retarded protein translation caused by a rare codon-rich gene. A mutant aminoacyl-tRNA synthetase (aaRS) with a low affinity to its cognate amino acid led to the retarded translation of the native protein, which could be recovered by enhanced intracellular synthesis of the corresponding amino acid. **c**. Codon usage and the anticodon of tRNAs in *E. coli*. **d**. Illustration of the AMINO strategy. The UAG codon introduced into a reporter gene was only decoded by the tRNA_CUA_. (1) Compared with the common tRNA, the tRNA_CUA_ has a poor charging capacity to its corresponding aaRS, (2) thus causing the retarded protein translation of the reporter gene, (3) The retarded translation could be restored when the intracellular corresponding amino acid is sufficient.

A novel screening method that conquers the intrinsic disadvantage of analogs has been developed based on protein translation (Fig. 1b, left column). A rare codon could be translated by the rare tRNA to encode its corresponding amino acid when the synonymous common tRNAs are preferentially charged by intracellular free amino acid ^10^. As the first instance of the protein translation-based strategy, *Escherichia coli* and *Corynebacterium glutamicum* overproducers of L-leucine, L-serine, or L-arginine have been obtained by coupling the concentrations of intracellular corresponding amino acid with cell growth or color formation using rare codon-substituted protein markers. A series of overproducers of L-proline have also been acquired using this rare codon-based strategy, leading to the high titer of the L-proline derivative (trans-4-hydroxy-L-proline) ^16^. Furthermore, efficient enzymes involved in amino acid overproduction could also be screened out ^17^. Nevertheless, this rare codon-based screening system could only be applied to the amino acids with multiple codons (Fig 1c) and tRNAs.

Another protein translation-based screening method has been recently established by using a mutant aminoacyl-tRNA synthetase (aaRS), which has a low affinity to its corresponding amino acid (Fig. 1b, right column). Strains harboring the mutant isoleucyl-tRNA synthetase showed retarded growth, thus L-isoleucine overproducers exhibiting fast-growth phenotype could be selected out ^18^. However, this strategy could only be applied to screen overproducers of an amino acid with a mutant aaRS of low amino acid affinity, which is currently unavailable for most amino acids, and it is time-consuming to mutate a specific aaRS.

The aminoacylation reaction belongs to the two-substrate sequential reaction ^19,20^. The affinity of the aaRS for amino acid and tRNA as well as the concentration of amino acid and tRNA determine the overall rate of aminoacylation reaction ^21-26^. In *E. coli*, the amino acid glutamate, glutamine, or arginine, binds the corresponding pre-assembled aaRS-tRNA complex. In contrast, each of the other 17 amino acids directly binds the corresponding aaRS to form an aaRS-aa complex as the basis for the following tRNA binding ^27-29^. When tRNA of glutamate, glutamine, or arginine binds to aaRS before amino acids, an impaired tRNA with lower affinity to aaRS could reduce the concentration of the aaRS-tRNA complex. Previous experimental data showed that the reaction rate of aa-tRNA formation between aaRS-tRNA and amino acid positively correlated with the concentration of amino acid until the aaRS-tRNA was saturated ^22^. In the presence of an impaired tRNA, a higher amino acid concentration could facilitate the reduced amount of aaRS-tRNA complex to generate aa-tRNA, which was supported by our theoretical calculation (Supplementary Note 1). In contrast, any tRNA of other 17 amino acids binds to preassembled aaRS-aa instead of free aaRS ^23,30^. The overall aminoacylation rate depends not only on the binding affinity between tRNA and aaRS-aa but also on the concentration of aaRS-aa until the concentration reaches saturation level ^23,30^. When an impaired tRNA with low binding affinity to aaRS-aa decreases the transfer of aa from aaRS-aa to tRNA, an increased concentration of amino acid could elevate the concentration of aaRS-aa and then facilitate the conversion of the impaired tRNA to generate aa-tRNA, in agreement with our theoretical calculation (Supplementary Note 1).

Taken together, if a low affinity of the tRNA partially represses the tRNA-dependent aminoacylation reaction, the increased supply of the amino acid might be able to partially rescue the overall aminoacylation rate (Fig. 1d), which is supported by previous experimental data ^31,32^ (Fig. 2a). Thus, an impaired tRNA could amplify the effect of amino acid concentration on the rate of aminoacylation reaction. Based on the above rationale, amino acid overproducers could be screened out by using a modified tRNA with impaired affinity to aaRS or aaRS-aa, depending on the targeted amino acid.

**Fig. 2.**
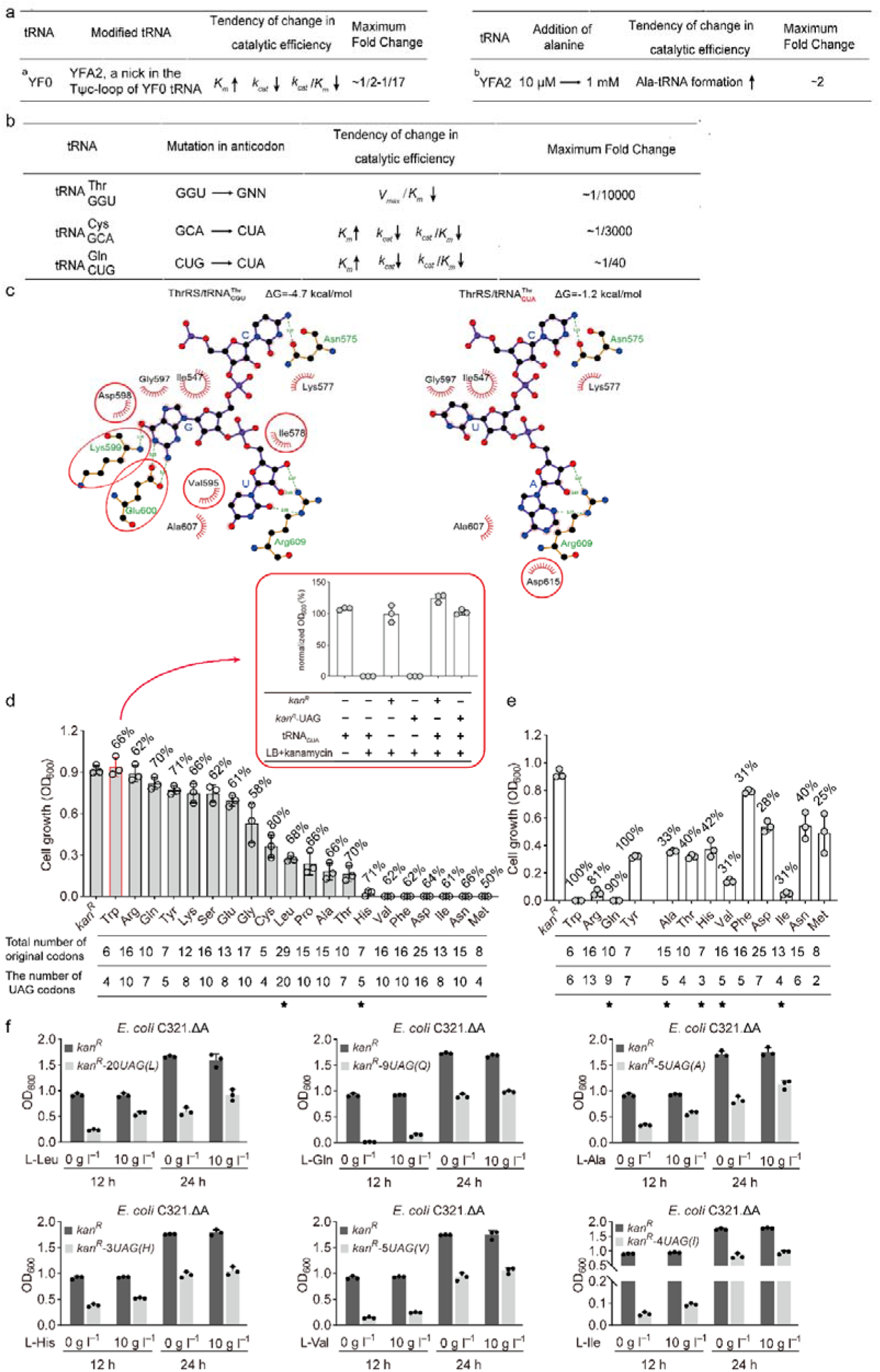
Design of the tRNA_CUA_-based system. **a**. Effect of the amino acid supplement on aminoacylation properties in the presence of a tRNA with a low affinity to its aaRS. ^a, b^ indicated data sources ^31,32^. **b**. Effect of the modifications in the anticodon of tRNA^Thr^, tRNA^Cys^, and tRNA^Gln^ on aminoacylation properties in *E. coli*. Fold change is normalized to a catalytic constant of 1.0 for the control substrates. **c**. The role of tRNA anticodon in the interaction between ThrRS-Thr and tRNA^Thr^. The detailed interaction of ThrRS-Thr residues with 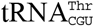(left) and 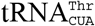 (right). **d**. Effects of the expression of UAG codon-rich *kan*^*R*^ variants on the growth of C321.ΔA strain in LB medium supplemented with kanamycin. Inset: the effect of the expression of tRNA_CUA_ on cell growth in the presence of kanamycin. The proportion and number of codon replacements by UAG were indicated on the top and bottom of the bar for each amino acid. **e**. Response of cell growth to increased UAG replacement for tryptophan, arginine, glutamine, and tyrosine codons, and decreased UAG replacement for alanine, threonine, phenylalanine, and other 6 amino acids in the presence of kanamycin. The OD_600_ was measured at 12 h in LB medium supplemented with kanamycin. The proportion and number of codon replacements by UAG were indicated on the top and bottom of the bar for each amino acid. The strains showing the positive response of amino acid feeding were marked as asterisks. **f**. Cell growth of *E. coli* C321.ΔA harboring pKan-20UAG(L), pKan-9UAG(Q), pKan-5UAG(A), pKan-3UAG(H), pKan-5UAG(V), and pKan-4UAG(I) in LB medium and LB medium supplemented with the corresponding amino acid in the presence of kanamycin.

There are multiple tRNA engineering strategies to decrease the binding affinity between tRNA and aaRS or aaRS-aa by modifying the critical structure of a tRNA, such as the anticodon or the rest part of the anticodon loop ^33^, the receptor stem ^34^, and the D-loop ^35^. For all proteinogenic amino acids except serine, leucine, and alanine, the corresponding aaRS directly interacts with the anticodon of the corresponding tRNA, and therefore the anticodon serves as the identity element of these tRNAs. Modifying the anticodon of the tRNA resulted in a lower *k*_*cat*_ and/or higher *K*_*m*_ compared with that of its original tRNA ^33,35-42^ (Fig. 2b, Supplementary Table 1). For serine, leucine, and alanine, the anticodon of tRNA might also be indirectly involved in the aminoacylation of tRNA since the modification of anticodon also resulted in the lower aminoacylation efficiency compared with that of its original tRNA ^43-45^.

Among the tRNA engineering strategies, modification of the anticodon might be the most convenient approach to lower the binding affinity of tRNA to aaRS. If the anticodon of the tRNA were modified, changing the original cognate codon in the reporter gene to one that pairs with the modified anticodon could easily restore the amino acid incorporation during translation. More importantly, the engineered codon-anticodon pair could couple the amino acid concentration with the translation efficiency of reporter genes. As a proof of concept, we employed the tRNA_CUA_ and the amber stop codon UAG to construct the screening and selection system. First, the UAG-decoding tRNA_CUA_ could be obtained from the engineered tRNAs of native proteinogenic amino acids (Fig. 1c) or the orthogonal aaRS/tRNA pairs of non-proteinogenic amino acids in various microbes such as *E. coli* ^46^, *Bacillus subtilis* ^47^, and *Salmonella typhimurium* ^48^. Furthermore, the amber stop codon UAG is a natural non-sense codon that has the lowest usage frequency among all 3 stop codons in *E*.*coli* (0.08 for UAG, 0.59 for UAA, and 0.33 for UGA) and other microbial species ^49^. Therefore, there is a minimal impact on the translation termination of genomic genes once the stop codon in structural genes was recognized by tRNA_CUA_. The presence of both the release factor 1 (RF1) and the tRNA_CUA_ might maintain the translation of the endogenous and exogenous genes harboring UAG, respectively. In this regard, it might be convenient to generate the growth-based selection system and the optical signal-based screening system, which are independent of toxic compounds, in amber-stop-codon-recoded or non-recoded microbes.

Here we report a strategy designated A tRN<u>A m</u>odification-based strategy for identifying ami<u>n</u>o acid <u>o</u>verproducers named AMINO. The tRNA_CUA_-based selection system according to the AMINO strategy was constructed by replacing a defined number of the codons of an individual amino acid with UAG in antibiotic resistance gene *kan*^*R*^. Strains harboring the UAG-equipped *kan*^*R*^ gene variants exhibited distinct growth phenotypes in the presence of kanamycin. Taking L-phenylalanine, L-aspartic acid, and L-tryptophan as instances, we demonstrated that the inhibitory effect of UAG substitution was correlated with the frequency of the UAG codon. As a proof-of-concept, overproducers of L-tryptophan, L-phenylalanine, and L-aspartic acid were screened out using the tRNA_CUA_-based selection system from mutation libraries of *E. coli* C321.ΔA strain, in which all 321 UAG amber stop codons in structural genes were replaced with the ochre stop codon UAA and the RF1 was knocked out. The underlying mechanism for the enhanced production of L-tryptophan in selected mutant strains from mutation libraries of *E. coli* C321.ΔA strain was further investigated by transcriptomic analysis and genomic sequencing. Finally, the tRNA_CUA_-based selection system was proved to be effective for other proteinogenic amino acids such as L-cysteine, L-lysine, L-glutamic acid, and L-glycine, as well as non-proteinogenic amino acids such as 3-methylhistidine and 5-hydroxytryptophan, in non-recoded *E. coli* strains (e.g., MG1655 derivative and XL10-gold) or *C. glutamicum*. As a proof-of-concept, a strain producing 20.3 g l^−1^ L-tryptophan in the bioreactor was constructed based on genotype analysis of mutants generated by genomic large-fragment deletions. The AMINO strategy based on this mechanism is universal and convenient that can be applied to screen overproducers of proteinogenic amino acids and non-proteinogenic amino acids.

## Results

### Design of the tRNA_CUA_-based system

To further verify the critical role of tRNA anticodon in the interaction between aaRS and tRNA, the available structures of class I and class II aaRSs such as threonyl-tRNA synthetase (ThrRS) ^50^ (PDB: 1QF6), cysteinyl-tRNA synthetase (CysRS) ^51^ (PDB: 1U0B), and glutaminyl-tRNA synthetase (GlnRS) ^52^ (PDB: 1O0B) were selected to generate the structures of ThrRS-tRNA_CUA_, CysRS-tRNA_CUA_, and GlnRS-tRNA_CUA_ by modifying the tRNA anticodon to CUA. In the case of ThrRS, the model with purple and bold bonds represents the tRNA anticodon base. The tRNA anticodon was labeled in blue uppercase letters. The model with yellow and thin bonds depicts the amino acid residues of ThrRS that interact with the tRNA anticodon. The interaction between the tRNA anticodon and the hydrophilic residues was indicated in green lines. The amino acid residues with red spokes pointing toward tRNA anticodon represent the residues involved in hydrophobic interactions. The hydrophilic and hydrophobic residues were labeled in green and black, respectively. The carbon atoms, oxygen atoms, and nitrogen atoms were indicated in black, red, and blue, respectively. The binding affinity of ThrRS and tRNA^Thr^ was calculated using the PDBePISA tool ^53^. If the calculated solvation-free energy (ΔG) increases, it represents that the binding affinity decreases. Compared with the wild-type 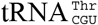, 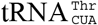 exhibited a weakened interaction with ThrRS-Thr at the anticodon region where the hydrogen bonding between 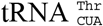 and Lys599, Glu600, respectively, as well as the hydrophobic interaction between 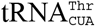 and Ile578, Val595, Asp598, respectively, were all broken (Fig. 2c). Correspondingly, the ΔG value between ThrRS-Thr and the anticodon region of 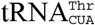 was changed from the original –4.7 kcal mol^−1^ to –1.2 kcal mol^−1^ after anticodon modification (Fig. 2c). The weakened interaction between the tRNA and the aaRS or aaRS-aa was also observed for Gln or Cys (Supplementary Fig. 1). These results suggested that replacing the tRNA anticodon of amino acids with CUA might reduce the interaction between the tRNA and the aaRS-aa or the aaRS when anticodon is involved in the recognition.

Accordingly, when the corresponding tRNA_CUA_ was expressed to decode the UAG codon, replacing the codon of 17 amino acids as well as Leu, Ala, and Ser with the UAG codon might result in a reduced translation rate of reporter genes. Furthermore, the frequency of the UAG codon substitution for different amino acids might influence the inhibitory effect since the translation rate of the UAG codon depends on the catalytic properties of each aaRS as well as the concentrations of amino acid and tRNA. To this end, a reporter gene *kan*^*R*^ encoding the aminoglycoside aminotransferase (3′) type 1a was subjected to UAG codon substitution. The tRNA_CUA_ of specific amino acid was then generated by substituting the native anticodon of the corresponding tRNA by CUA, which was overexpressed in the C321.ΔA strain under the control of a constitutive lpp promoter ^54^ (Supplementary Fig. 2). It is convenient to demonstrate the inhibitory effect of tRNA_CUA_ on UAG-rich gene expression in the C321.ΔA strain due to the lack of competition from RF1. To test the inhibitory effect of UAG substitution on the translation of *kan*^*R*^, approximately 70% of the codons of each amino acid individually of the *kan*^*R*^ gene were substituted by UAG. As shown in Fig. 2d, a strain containing *kan*^*R*^*-UAG* gene variants exhibited variable growth phenotypes in the presence of kanamycin. The growth of strains was not detectable when the UAG codon encodes Val, Phe, Asp, Ile, Asn, and Met, individually. Compared with the wild-type *kan*^*R*^, the growth of strain containing *kan*^*R*^*-UAG* gene variants was not significantly impacted when the UAG codon encodes Trp, Arg, Gln, and Tyr. These results suggested that the expression of the *kan*^*R*^*-UAG* genes might be related to the number of the UAG codon and the codon of the specific amino acid.

We then investigated the effects of tuning the UAG codon frequency on the expression of the reporter gene. For those strains carrying the *kan*^*R*^-variants containing UAG codons for Ala, Thr, His, Val, Phe, Asp, Asn, or Met, the cell growth was inhibited which could be rescued by decreasing the number of UAG codons in *kan*^*R*^ to below 45% of the total codon number of the corresponding amino acid in the presence of kanamycin (Fig. 2e). For those such as Trp, Arg, Gln, and Tyr that did not induce growth inhibition, replacing 80∼100% of their codons in *kan*^*R*^ with UAG induced growth retardation in the presence of kanamycin. The preliminary evidence suggested that the expression of the UAG-rich marker gene could be fine-tuned by changing the frequency of the UAG codon.

The translational dependency of *kan*^*R*^*-UAG* gene variants on the tRNA_CUA_ was also investigated using pKan-4UAG(W). As shown in the inset of Fig. 2d, the strain harboring pKan-4UAG(W) failed to grow in the absence of the tRNA_CUA_ but grew normally in the presence of the tRNA_CUA_. In addition, the overexpression of the tRNA_CUA_ had no significant impact on cell growth in the absence of the *kan*^*R*^*-4UAG(W)* gene. These results suggested overexpression of the tRNA_CUA_ is essential for translating the UAG codons in *kan*^*R*^.

We preliminarily investigated whether the supply of amino acid could partially restore the growth retardation of the strain harboring UAG-rich *kan*^*R*^ genes in Fig. 2d, e. As shown in Fig. 2f, the OD_600_ values of strains harboring pKan-20UAG(L), pKan-9UAG(Q), pKan-5UAG(A), pKan-3UAG(H), pKan-5UAG(V), and pKan-4UAG(I) were increased when 10 g l^−1^ the corresponding amino acid was supplemented in LB medium in the presence of kanamycin. These results demonstrated that the addition of amino acids could elevate the reduced rate of aminoacylation caused by tRNA_CUA_.

### Effects of UAG codon frequency on the selection system

To further explore the inhibitory effect of UAG codon frequency on the translation rate of the reporter gene, the strains exhibiting the arrested-growth phenotype were first investigated using strain containing pKan-10UAG(F) and pKan-16UAG(D) (Fig. 3a). To alleviate the growth-inhibition effect, we gradually reduced the number of UAG codons in *kan*^*R*^. After 12 h incubation in LB medium supplemented with kanamycin, the OD_600_ of the strain carrying pKan-8UAG(F) and pKan-7UAG(F) recovered to 10.7% and 32.2% of that of the strain harboring pKan, respectively. Such frequency-dependent inhibition manner had been observed in a series of diluted LB media (Fig. 3b, left) which might help to magnify the growth difference between the strain harboring the UAG-rich *kan*^*R*^ and the one harboring wild-type *kan*^*R*^. Similarly, the growth of strain harboring pKan-7UAG(D) was recovered to 45.1% of that of the strain harboring pKan while strain harboring pKan-9UAG(D), pKan-13UAG(D), pKan-15UAG(D), and pKan-16UAG(D) did not show significant growth (Fig. 3b, middle). These results suggested that cell growth could be improved by decreasing the number of UAG codons in *kan*^*R*^.

**Fig. 3.**
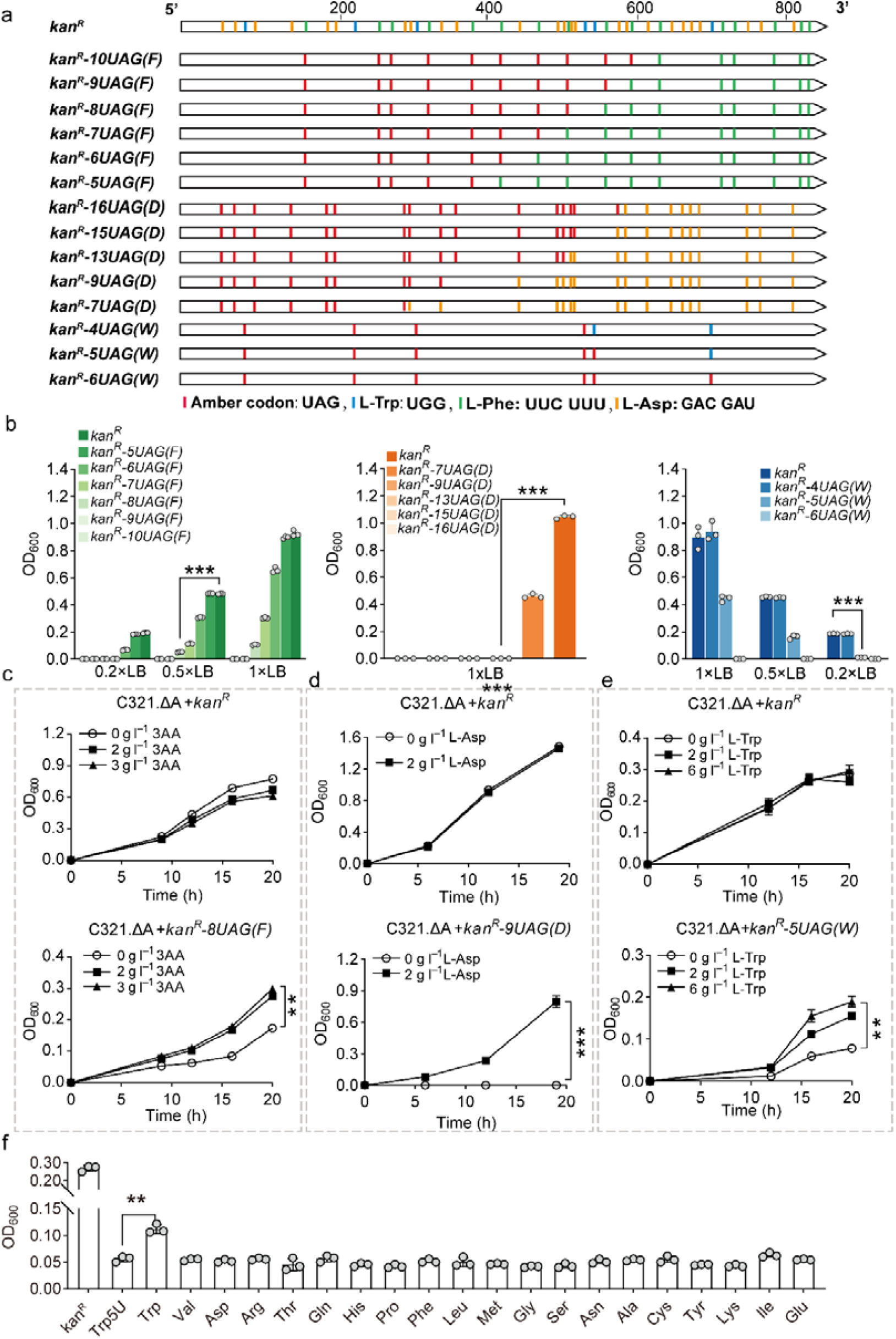
Effects of the frequency of UAG codon and amino acid feeding on *kan*^*R*^ expressions. **a**. Different numbers of the original codons of L-phenylalanine, L-aspartic acid, and L-tryptophan on the wild-type *kan*^*R*^ were replaced by UAG codon, generating *kan*^*R*^*-10UAG(F), kan*^*R*^*-9UAG(F), kan*^*R*^*-8UAG(F), kan*^*R*^*-7UAG(F), kan*^*R*^*-6UAG(F), kan*^*R*^*-5UAG(F), kan*^*R*^*-16UAG(D), kan*^*R*^*-15UAG(D), kan*^*R*^*-13UAG(D), kan*^*R*^*-9UAG(D), kan*^*R*^*-7UAG(D), kan*^*R*^*-4UAG(W), kan*^*R*^*-5UAG(W)*, and *kan*^*R*^*-6UAG(W)*. **b**. Influences of different numbers of the L-phenylalanine, L-aspartic acid, and L-tryptophan codons replaced by UAG codon on OD_600_ of C321.ΔA strain. **c**. Effects of feeding a mixture of three amino acids (3AA: L-phenylalanine, L-tryptophan, and L-tyrosine) in 0.5×LB supplemented with kanamycin on the growth of C321.ΔA strain harboring pKan-8UAG(F), in which eight L-phenylalanine codons on *kan*^*R*^ were substituted by UAG codon. **d**. Effect of 2.0 g l^−1^ L-aspartic acid and 2.0 g l^−1^ MOPS in LB supplemented with kanamycin on the growth of C321.ΔA strain harboring pKan-9UAG(D). **e**. Effects of feeding 2.0 g l^−1^ and 6.0 g l^−1^ L-tryptophan in 0.2×LB supplemented with kanamycin on cell growth of C321.ΔA strain harboring pKan-5UAG(W), in which five tryptophan codons on *kan*^*R*^ were replaced by UAG codon. **f**. Changes in ODs of C321.ΔA strain harboring pKan-5UAG(W) after cultivation in 0.2×LB in the presence of kanamycin supplemented with 2.0 g l^−1^ 20 proteinogenic amino acids at 30°C for 16 h. Values and error bars represent the mean and the s.d. (n = 3, \*\**P* < 0.01, \*\*\**P* < 0.001 as determined by two-tailed t-test).

We then hypothesized that the inhibitory effect of UAG substitution on cell growth could be enhanced by increasing the number of UAG codons (Fig. 3a). To test this, the strain carrying pKan-4UAG(W) that exhibited no significant response (Fig. 2d) was utilized for further modification. As a result, the OD_600_ of the strain carrying pKan-5UAG(W) was decreased to 49.6%, 35.4%, and 6.7% of that of the strain harboring pKan in LB, 0.5×LB, and 0.2×LB medium supplemented with kanamycin, respectively (Fig. 3b, right). The growth of strain harboring pKan-6UAG(W) was not detectable in these media. Similarly, when all the L-tryptophan codons of fluorescence reporter gene *rfp* were replaced with UAG codons, the expression of RFP was not detectable (Supplementary Fig. 3). These results further confirmed that the inhibitory effect of UAG substitution was positively correlated with the frequency of the UAG codon.

### Feeding amino acids restored cell growth

To further investigate whether the supply of amino acids could alleviate the induced growth inhibition in the presence of the appropriate amount of UAG codons, the growth of the strain harboring pKan-8UAG(F), pKan-9UAG(D), and pKan-5UAG(W) was investigated in the medium supplemented with different concentrations of the corresponding amino acids in the presence of kanamycin. As a result, feeding amino acids significantly restored the growth of the strain harboring the UAG-rich *kan*^*R*^ genes while it had a negligible effect on the growth of the strain harboring the wild-type *kan*^*R*^. Specifically, the OD_600_ of the strain harboring pKan-8UAG(F) in a 0.5×LB medium was increased 71.8% by feeding aromatic amino acids mixtures (3AA) consisting of 3.0 g l^−1^ L-phenylalanine, 3.0 g l^−1^ L-tryptophan, and 0.3 g l^−1^ L-tyrosine (Fig. 3c), as the sole supply of 3.0 g l^−1^ L-phenylalanine would inhibit the growth of the strain harboring pKan. Similarly, the addition of 2.0 g l^−1^ of L-aspartic acid significantly restored the growth of the strain harboring pKan-9UAG(D) (Fig. 3d). Adding 2.0 g l^−1^ of L-tryptophan and 6.0 g l^−1^ of L-tryptophan to the 0.2×LB medium increased the OD_600_ of the strain harboring pKan-5UAG(W) by 42.8% and 71.0%, respectively (Fig. 3e).

To modify the stringency of the selection system, the expression level of tRNA_CUA_ was regulated using promoters with different strengths. When tRNA_CUA_ was driven by a weak promoter *P*_*J23106*_ (iGEM Part: BBa_J23106) ^55,56^, the cell growth in the 0.2×LB medium supplemented with kanamycin was 59% of that driven by the *lpp* promoter (Supplementary Fig. 4). These results suggested that the stringency of the selection system could be regulated by adjusting the expression strength of tRNA_CUA_ which correlated positively with cell growth ^57^.

**Fig. 4.**
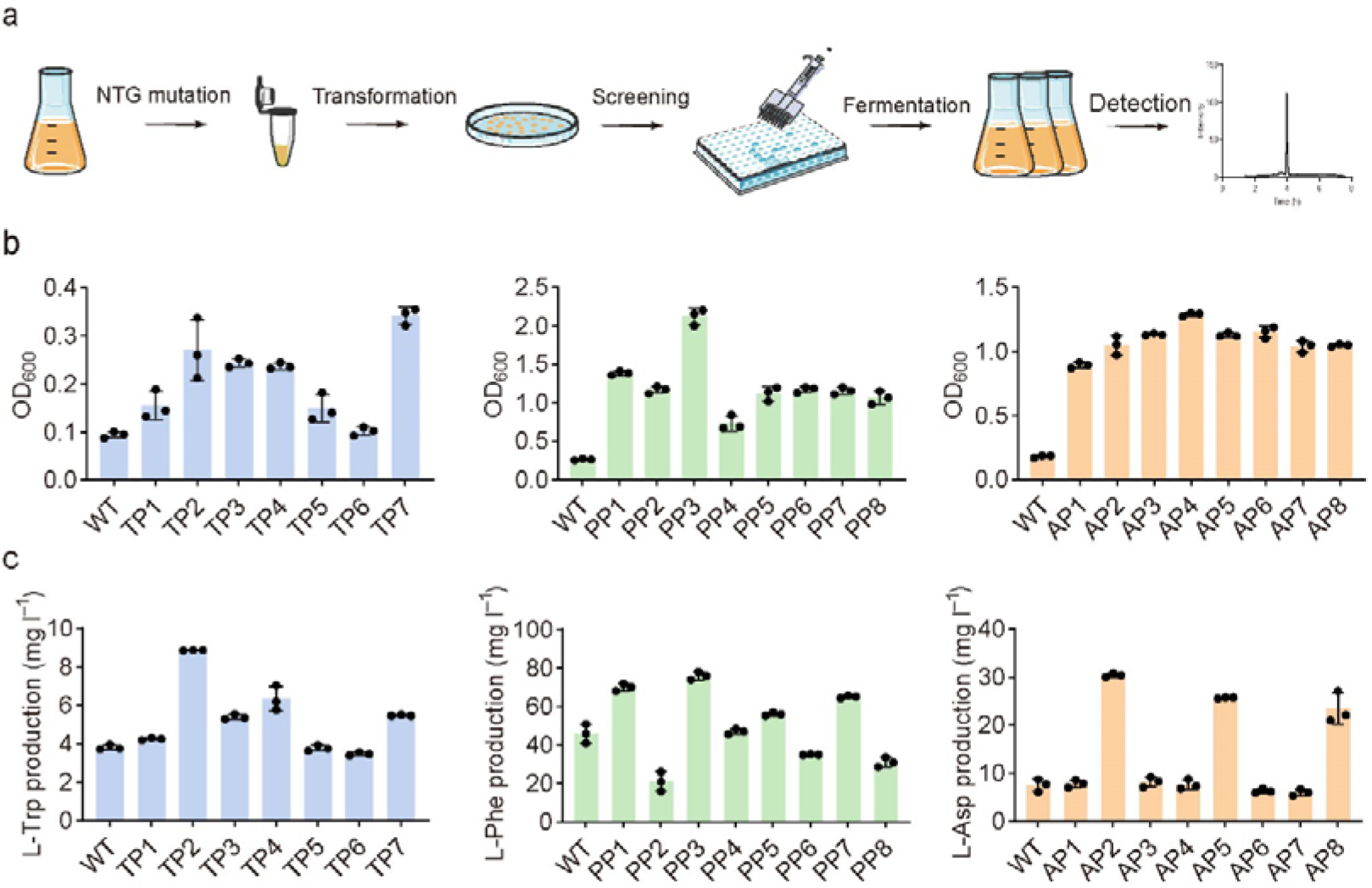
Obtaining amino acid overproducers by the tRNA_CUA_-based system. **a**. Workflow of the tRNA_CUA_-based strain selection. C321.ΔA strain cultured to the exponential phase in LB medium was resuspended in 0.1 M sodium citrate solution (pH 5.5) containing 50 μg ml^−1^ NTG for 45, 60, and 75 minutes to induce mutations. The mutagenesis library of the C321.ΔA strain was made into competent cells and transformed with pKan-5UAG(W) for the selection of L-tryptophan overproducers. The cells exhibiting superior growth in 0.2×LB medium supplemented with kanamycin were selected and inoculated into the fermentation medium. The production of L-tryptophan, L-phenylalanine, and L-aspartic acid in these cultures was measured by HPLC. **b**. Cell growth of various mutants selected by the tRNA_CUA_-based system after cultivation in 0.2×LB and LB media supplemented with kanamycin at 30°C. **c**. Titers of L-tryptophan, L-phenylalanine, and L-aspartic acid produced by C321.ΔA mutants after cultivation in M9-modified medium at 30°C for 60 h, respectively. Values and error bars represent the mean and the s.d. (n = 3).

To verify the fidelity of the selection system, we investigated the growth of the strain carrying pKan-5UAG(W) or pKan-9UAG(D) by feeding other 19 amino acids besides the one of interest in the presence of kanamycin. For the L-tryptophan, a 42% increase in cell growth was observed when feeding 2.0 g l^−1^ L-tryptophan, while the other 19 amino acids showed no significant improvement in cell growth (Fig. 3f). High fidelity was also observed for L-aspartic acid as the cell growth could only be restored by feeding L-aspartic acid (Supplementary Fig. 5).

**Fig. 5.**
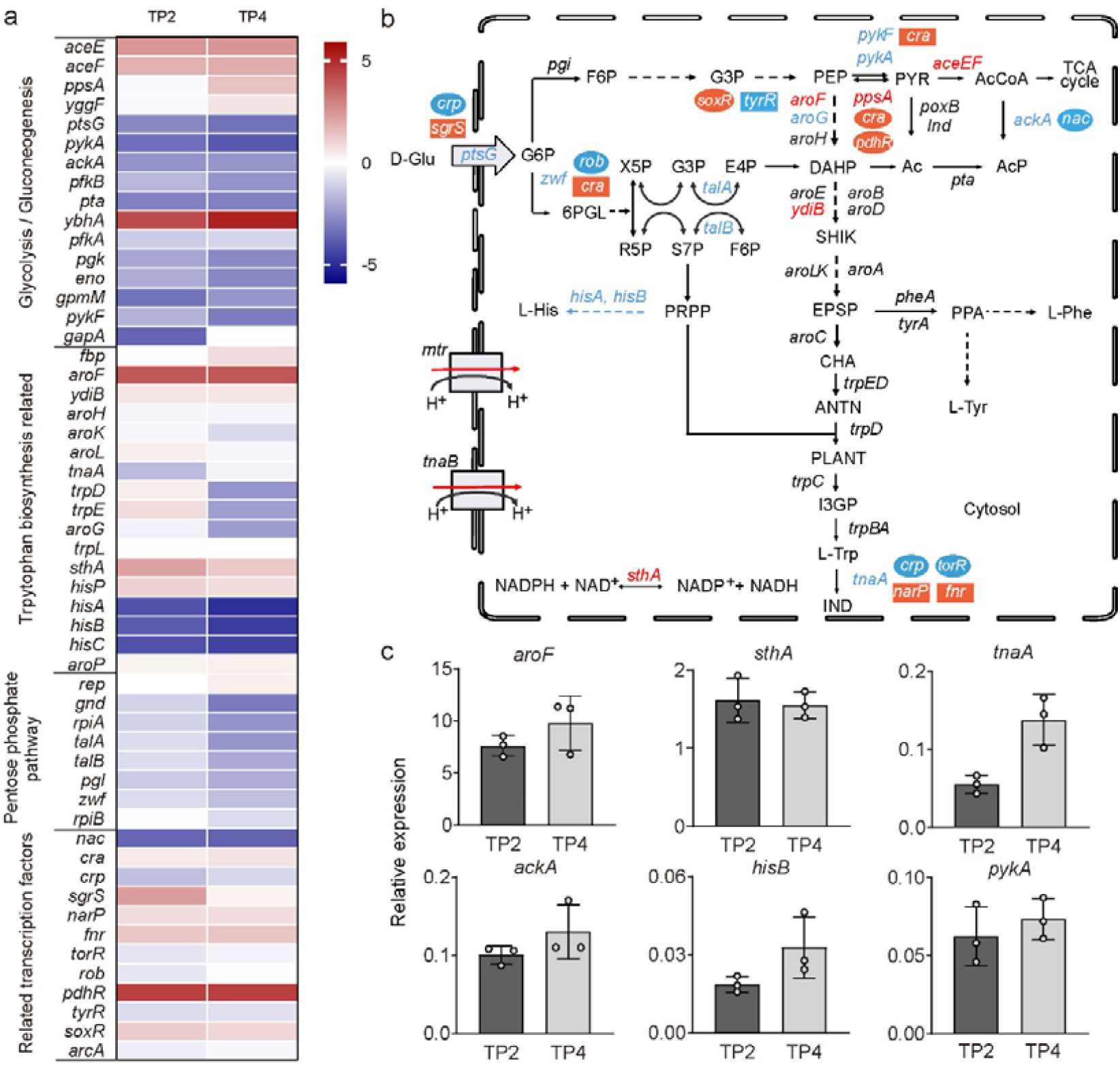
Transcription profiles of L-tryptophan biosynthetic pathway in TP2 and TP4. **a**. Transcriptome analysis of TP2, TP4, and the C321.ΔA strains. Red indicates up-regulated genes whereas blue represents down-regulated genes, according to the RPKM of TP2 and TP4 divided by that of the wild-type strain (in log2). Genes involved in glycolysis, L-tryptophan biosynthesis, pentose phosphate (PP) pathway, and related regulation are presented in the heatmap. **b**. The L-tryptophan biosynthetic pathway in *E. coli*. Genes colored in red and blue indicate the up- and down-regulated genes, respectively. The oval and rectangle represent transcription activators and repressors for the corresponding gene, respectively, with up-regulated factors in orange and down-regulated ones in blue. Abbreviations: Glu, glucose; G6P, glucose-6-phosphate; G3P, glyceraldehyde-3-phosphate; 3-PYR, 3-phosphonooxypruvate; PEP, phosphoenolpyruvate; AcCoA, acetyl coenzyme A; Ac, acetate; AcP, acetyl phosphate; IND, indole; X5P, xylulose-5-phosphate; E4P, erythrose-4-phosphate; R5P, ribose-5-phosphate; S7P, sedoheptulose-7-phosphate; F6P, fructose-6-phosphate; PRPP, phosphoribosyl pyrophosphate; PYR, pyruvate; DAHP, 3-deoxy-D-arabinoheptulosonate 7-phosphate; SHIK, shikimate; EPSP, 5-enolpyruvyl shikimate 3-phosphate; CHA, chorismate; ANTN, anthranilate; PLANT, N-(5′-phosphoribosyl)-anthranilate; I3GP, (1S,2R)-1-C-(indol-3-yl) glycerol 3-phosphate); PPA, prephenate; Trp, tryptophan. Phe, phenylalanine; Tyr, tyrosine. **c**. qRT-PCR verification of the genes related to L-tryptophan biosynthesis. Values and error bars represent the mean and the s.d. (n = 3).

### Obtaining amino acid overproducers by the selection system

To assess the validity of the selection system, the system was employed to screen amino acid overproducers using the plasmid pKan-5UAG(W) (Fig. 4a). The mutagenesis library of the C321.ΔA strain was generated by NTG treatment for 0, 45, 60, and 75 mins with fatality rates of 0, 57, 88, and 92%, respectively (Supplementary Fig. 6). After the treatment, mutants were cultivated and made into competent cells. The pKan-5UAG(W) was transformed into the mutation libraries for the selection of L-tryptophan overproducers. The top seven mutants with the highest OD_600_ values (0.10∼0.34) were selected in 0.2×LB medium supplemented with kanamycin (Fig. 4b). Among these strains, the strain TP2 produced L-tryptophan at a titer of 8.9 mg l^−1^, which was 2.3-fold that of the reference strain. Other mutant strains such as TP3, TP4, and TP7 achieved titers of 5.5, 6.4, and 6.5 mg l^−1^ (Fig. 4c). These results indicated that the tRNA_CUA_-based system could be applied to the C321.ΔA strain for obtaining mutant strains with improved L-tryptophan production. Based on the same strategy, three and four mutant strains with high production of L-aspartic acid and L-phenylalanine were identified from a UV mutation library using pKan-9UAG(D) and pKan-8UAG(F), respectively. Compared with the reference strain, L-aspartic acid production was increased by 2∼4 folds, achieving the highest titer in a shake-flask fermentation. For L-phenylalanine, the mutant strains showed up to a 1.6-fold increase in the production of L-phenylalanine compared with that of the reference strain (Fig. 4c).

**Fig. 6.**
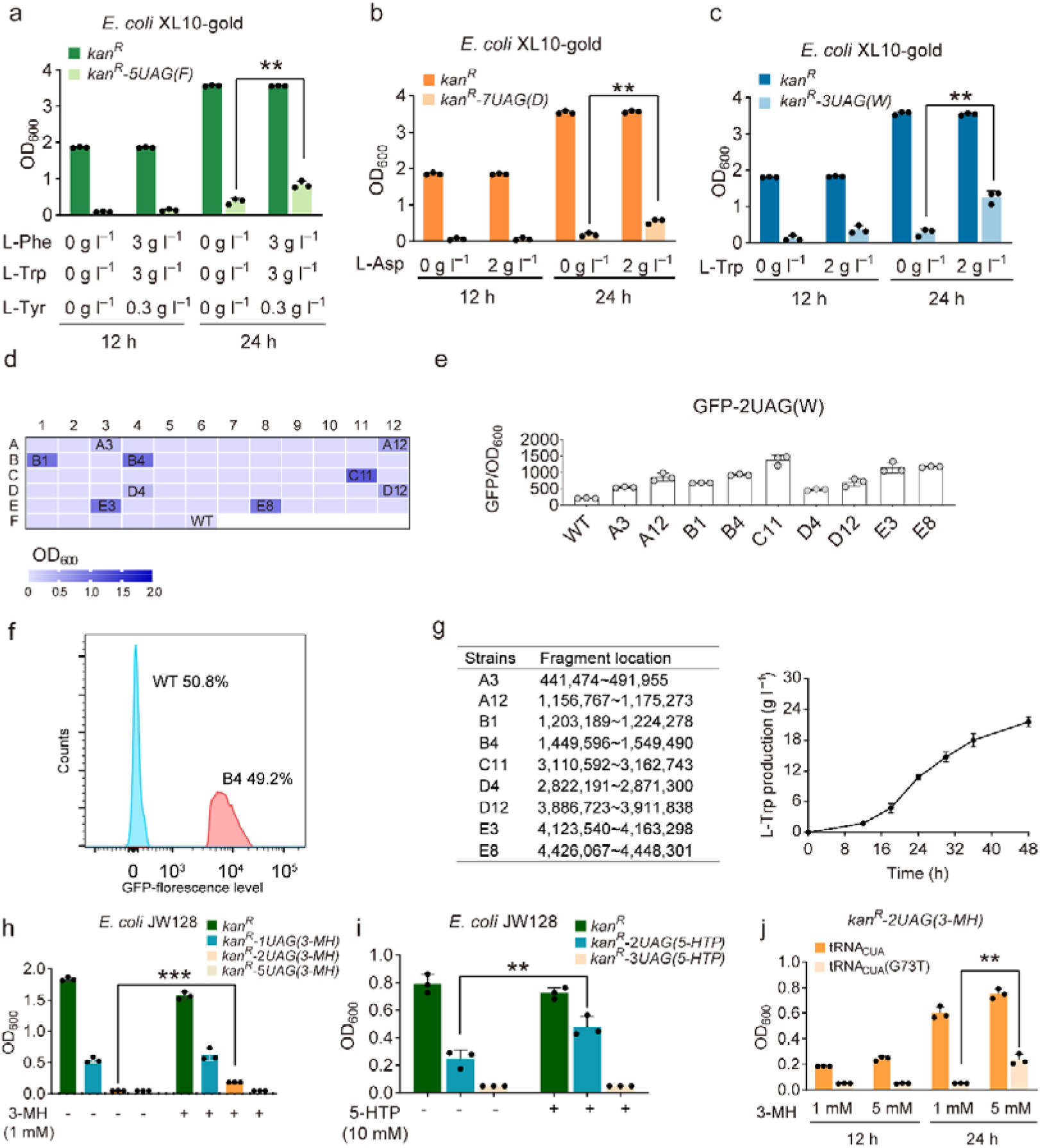
Application of the tRNA_CUA_-based system on the proteinogenic and non-proteinogenic amino acids in *E. coli* strains harboring the native UAG codons in the genome. Cell growth of *E. coli* XL10-gold harboring **a**. pKan-5UAG(F), **b**. pKan-7UAG(D), and **c**. pKan-3UAG(W) in TB medium and TB medium supplemented with kanamycin and the corresponding amino acids. Selection of candidate L-tryptophan overproducers from a library of MG1655 derivative strains by using **d** pKan-3UAG(W) and **e** pGFP-2UAG(W), based on cell growth and fluorescence, respectively. **f**. Application of fluorescence-activated cell sorting to screen mutant strains according to the fluorescence level of the mutant strain B4 and the wild-type strain. **g**. Selection using the genomic large-fragment-deletion library, and the fed-batch fermentation titer of a reverse-engineered L-tryptophan overproducer. Locations of the deleted fragments were labeled according to the MG1655 genome with accession No. U00096. **h**. Effects of the frequency of UAG codon and feeding of 3-methylhistidine (3-MH) on *kan*^*R*^ expressions in LB medium supplemented with kanamycin. **i**. Effects of the frequency of UAG codon and feeding of 5-hydroxytryptophan (5-HTP) on *kan*^*R*^ expressions in the presence of kanamycin. The right bars in the figure showed the cell growth when 10 mM 5-hydroxytryptophan was supplemented in a 0.5×LB medium in 24 h cultivation. **j**. Effects of the modification of tRNA accept stem and feeding 1 mM and 5 mM of 3-methylhistidine (3-MH) on the cell growth for strain JW128 harboring pKan-2UAG(3-MH) in LB medium supplemented with kanamycin.

### Mechanisms of increased amino acid production

To investigate the underlying mechanism for the enhanced production of L-tryptophan, RNA sequencing was performed on the C321.ΔA strain as well as the mutant strains TP2 and TP4. Forty-one genes involved in aromatic amino acids biosynthesis, the glycolysis pathway (EMP), and the pentose phosphate (PP) pathway were identified by differential gene expression analysis (Fig. 5a). Specifically, *pykA* encoding pyruvate kinase II was significantly downregulated 10.5- and 15.1-fold in TP2 and TP4 compared with that of the C321.ΔA strain. The *ackA* gene encoding acetate kinase, which catalyzes the formation of acetate from acetyl-CoA, was down-regulated 5.8- and 5.6-fold in TP2 and TP4, respectively. The expressions of *hisA, hisB*, and *hisC*, which are involved in L-histidine biosynthesis, were also downregulated in TP2 and TP4. In contrast, several genes involved in the biosynthesis of L-tryptophan showed upregulated expression in mutant strains. For example, the expression of *aroF* that catalyzes the biosynthesis of 3-deoxy-D-arabinoheptulosonate 7-phosphate, the key precursor of L-tryptophan, was increased up to 14.5- and 15.1-fold, in TP2 and TP4, respectively. In addition, *tnaA* encoding the tryptophanase, which is involved in the degradation of L-tryptophan, decreased 3.1- and 1.2-fold in TP2 and TP4. Moreover, cofactor metabolism was also optimized in the mutant strains, in which *sthA* encoding soluble pyridine nucleotide transhydrogenase for the interconversion of NADH and NADPH was upregulated approximately 1.5-fold in TP2 and TP4 (Fig. 5b). These results suggested that cofactor regulation and flux redirection by enhancing the L-tryptophan biosynthetic pathway along with depressing the competing and degradation pathways are beneficial for the production of L-tryptophan (Fig. 5b). The qRT-PCR analysis using *aroF, sthA, tnaA, ackA, hisB*, and *pykA* as instances showed trends similar to the transcriptomic results (Fig. 5c). Changes in gene transcript levels are positively correlated with their corresponding regulators (Supplementary Table 2).

To identify key mutations in the mutant strains with improved L-tryptophan production, genomic sequencing was performed on the mutants and the reference strain. Several mutations that are related to amino acid accumulation were identified (Supplementary Table 3), including the S326F for L-aromatic amino acid transporter AroP and the H389Y for L-phenylalanine transporter PheP, both involved in L-tryptophan accumulation.

### Applicability of the selection system to hosts without amber stop codon replacement

The above screening system was established in the C321. ΔA strain. To explore the applicability of the AMINO strategy, two *E. coli* strains retaining the native UAG codons in the genome, JW128 and XL10-gold were used as the initial hosts. JW128 ^58^ is a genetically manipulable *E. coli* strain derived from the MG1655, whereas XL10-gold is usually applied in gene cloning. Compared with the JW128 and XL10-gold strains harboring pKan, the growth of JW128 and XL10-gold strains harboring pKan-5UAG(F), pKan-7UAG(D), and pKan-3UAG(W) significantly decreased in LB or TB media supplemented with kanamycin, respectively. After feeding the corresponding amino acids, the growth of XL10-gold harboring pKan-5UAG(F) (Fig. 6a), pKan-7UAG(D) (Fig. 6b), and pKan-3UAG(W) (Fig. 6c) was increased approximate 2-fold, 3.1-fold, and 3.6-fold in TB medium supplemented with kanamycin, respectively. For the JW128 harboring pKan-5UAG(F), pKan-7UAG(D), and pKan-3UAG(W), increased cell growth was also observed in LB or TB media supplemented with kanamycin, respectively, when the corresponding amino acid was supplemented (Supplementary Fig. 7). These results suggested that the AMINO strategy is independent of the amber-stop-codon-recoded strain.

To demonstrate the applicability of the tRNA_CUA_-based selection system in selecting amino acid overproducers from non-recoded strains, the plasmid pKan-3UAG(W) was introduced into an unpublished library of 65 MG1655 derivative strains with different genomic large-fragment deletions ranging from 14 to 143 kb. A total of nine strains exhibiting growth advantages were selected in the LB medium supplemented with kanamycin, respectively (Fig. 6d and 6e). Similarly, the higher GFP/OD_600_ of the selected mutant strains harboring pGFP-2UAG(W) was observed compared with that of the wild-type strain harboring pGFP-2UAG(W) after 12 h cultivation. To evaluate the efficiency of the tRNA_CUA_-based selection, the mixed culture containing mutants harboring pGFP-2UAG(W) and the wild-type strain harboring pGFP-2UAG(W) in a 1:1 ratio was analyzed using fluorescence-activated cell sorting (FACS). The sorting ratio, the top 1%, was set to screen candidates with high fluorescence intensity. PCR analysis was employed to verify the genotype of selected strains. The difference in the fluorescent signal was observed in Fig. 6f and Supplementary Fig. 8. Ten colonies of the selected candidates exhibiting higher fluorescence intensity were further proved to be the mutant strains by colony PCR. Based on the analysis of the deleted genomic fragments, multiple genes involved in aromatic amino acid metabolism (e.g., *tnaA*), carbon assimilation (e.g., *ptsG*), and nitrogen regulation (e.g., *glnK*), were discovered, the deletion of which is likely to favor L-tryptophan production. We further knocked out related genes (e.g., *trpR, tnaAB, mtr*) and overexpressed the L-tryptophan biosynthetic pathway (e.g., *trpABCDE*) in the JW128. The recombinant strain TP10 is capable of producing 20.3 g l^−1^ L-tryptophan in the bioreactor (Fig. 6g). The productivity is 0.42 g l^-1^ h^-1^ and the yield is 0.15 g g^-1^ glucose. These results suggested that the tRNA_CUA_-based selection system could screen variants with the better productive performance from the mutant library, which could be utilized to de novo engineer the amino acid overproducers.

In addition, the tRNA_CUA_-based selection system was applied to two non-proteinogenic amino acids, 3-methylhistidine and 5-hydroxytryptophan. For 3-methylhistidine, the codons encoding His3 and His183 on *kan*^*R*^ were replaced with UAG, and the aaRS-tRNA pair for incorporation of 3-methylhistidine into the protein was also employed to assemble the tRNA_CUA_-based system. Strain JW128 harboring such a system did not show significant growth in the presence of kanamycin unless 3-methylhistidine, not even for L-histidine was added. A positive response in OD_600_ of up to 0.24 was observed after 12 h cultivation when the concentration of supplemented 3-methylhistidine increased from 1 mM to 5 mM (Fig. 6h and 6j). For 5-hydroxytryptophan, the increased growth of strain JW128 harboring pKan-2UAG(5-HTP) was also observed when 10 mM 5-hydroxytryptophan was supplemented in the presence of kanamycin (Fig. 6i). Hence, the selection system could be applied to select overproducers of 3-methylhistidine and 5-hydroxytryptophan, as well as the potential enzymes that convert L-histidine and L-tryptophan to 3-methylhistidine and 5-hydroxytryptophan, respectively.

To verify whether modification of other parts of tRNA outside of the anticodon region could be used to construct the selection system, the G73 within the accept stem of tRNA was mutated to T for decreasing the tRNA binding affinity to aaRS of 3-methylhistidine ^34^. As shown in Fig. 6j, the strain harboring tRNA (G73T) could not grow until 24 h cultivation in the presence of kanamycin and 1 mM 3-methylhistidine. When 5 mM 3-methylhistidine was supplemented, the OD_600_ of the strain harboring tRNA (G73T) could be restored to approximately 30% of that of the strain harboring the original tRNA, indicating that the G73T tRNA could be utilized to screen 3-methylhistidine overproducers. These results suggested that the target for tRNA modification in the AMINO strategy is not limited to the anticodon region.

### An alternative tRNA_CUA_-based selection system

To facilitate the adoption of the selection system in selecting overproducers of any amino acid, a modified selection marker was designed by adding a UAG-rich leader tag at the 5’-end of the *cm*^*R*^. Translation of the tagged *cm*^*R*^ containing multiple UAG codons should be terminated in the absence of the corresponding tRNA_CUA_. Therefore, a tRNA_CUA_ carrying an amino acid is not only essential to read through the UAG-rich tagged *cm*^*R*^ but also determines the amino acid overproducer to be selected. The expression of the tagged *cm*^*R*^ should correlate positively with an increased concentration of amino acid of interest (Fig. 7a). The tagged *cm*^*R*^ offered an alternative marker for the tRNA_CUA_-based system. When this marker was applied to the different amino acids, only the tRNA_CUA_ need to be changed according to the amino acid of interest.

**Fig. 7.**
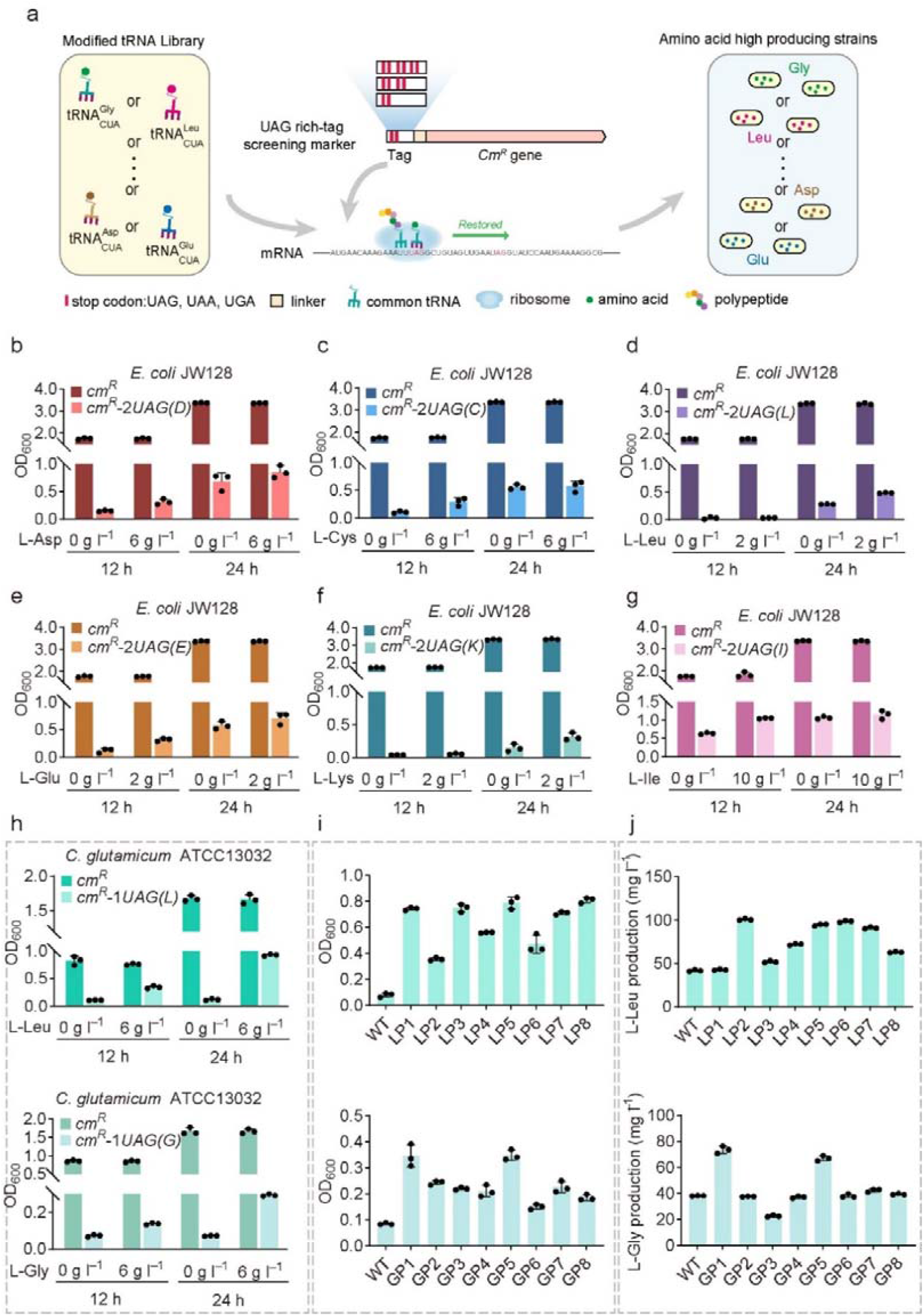
Application of the tRNA_CUA_-based system on the proteinogenic amino acid in *E. coli* and *C. glutamicum* strains. **a**. Illustration of an alternative selection marker consisting of a UAG-rich leader tag at the 5’-end of the *cm*^*R*^ gene with tRNA_CUA_ of the targeted amino acid. Cell growth of *E. coli* JW128 harboring **b** pCm-2UAG(D), **c** pCm-2UAG(C), **d** pCm-2UAG(L), **e** pCm-2UAG(E), **f** pCm-2UAG(K), and **g** pCm-2UAG(I) in LB medium and LB medium supplemented with the corresponding amino acids in the presence of chloramphenicol. **h**. Cell growth of *C. glutamicum* ATCC13032 harboring pCm-1UAG(L) and pCm-1UAG(G) in 2×LBHIS medium supplemented with the corresponding amino acids in the presence of chloramphenicol. **i**. Cell growth of various mutants selected by the tRNA_CUA_-based system after cultivation in 2×LBHIS medium in the presence of chloramphenicol at 30°C for 16 h. **j**. Titers of L-leucine and L-glycine produced by *C. glutamicum* ATCC13032 mutants after cultivation in CGXII medium at 30°C for 48 h. Values and error bars represent the mean and the s.d. (n =3).

To demonstrate its reliability, such a system was first applied in the JW128. Taking L-aspartic acid (Fig. 7b), L-cysteine (Fig. 7c), L-leucine (Fig. 7d), L-glutamate (Fig. 7e), L-lysine (Fig. 7f), and L-isoleucine (Fig. 7g) as instances, the cell densities of JW128 harboring pCm-2UAG(D), pCm-2UAG(C), pCm-2UAG(L), pCm-2UAG(E), pCm-2UAG(K) or pCm-2UAG(I) were lower than that of the strain harboring pCm in the LB medium supplemented with 50 mg l^−1^ chloramphenicol. When different concentrations of the corresponding amino acids were supplemented, cell densities of JW128 harboring pCm-2UAG(D), pCm-2UAG(C), pCm-2UAG(L), pCm-2UAG(E), pCm-2UAG(K) or pCm-2UAG(I) were increased in varying degrees. Furthermore, *C. glutamicum*, a common host for amino acid fermentation, was utilized to demonstrate the versatility of this system, exemplified here with L-leucine and L-glycine. To apply the tRNA_CUA_-based system in *C. glutamicum*, plasmids pCm-1UAG(L) and pCm-1UAG(G) were introduced into the strain ATCC 13032, respectively. As shown in Fig. 7h, the cell densities of the strain harboring pCm-1UAG(L) and pCm-1UAG(G) were significantly lower than that of the strain harboring pCm in a 2×LBHIS medium containing 15 mg l^−1^ chloramphenicol. Similarly, cell densities of the strain harboring pCm-1UAG(L) and pCm-1UAG(G) were partially restored when the 6 g l^−1^ corresponding amino acid was supplemented. Based on the above system, the eight mutants with the highest OD_600_ values for these two amino acids were selected from a UV mutation library, respectively (Fig. 7i). Among these strains, the strains LP2, LP5, LP6, and LP7 produced approximate 100 mg l^−1^ L-leucine, which was about 2-fold that of the wild-type strain. Other mutant strains such as GP1 and GP5 achieved an L-glycine titer of 73.5 and 67 mg l^−1^, which was approximately 2-fold that of the wild-type strain, respectively (Fig. 7j). These results suggested that the AMINO strategy could be employed to obtain overproducers of L-leucine and L-glycine in *C. glutamicum*.

## Discussion

In this study, we developed a selection strategy that could be applied to screen overproducers of proteinogenic and non-proteinogenic amino acids. This strategy is universal, convenient, and nontoxic, which takes advantage of the critical structural element of the tRNA in living organisms. Such a system based on this strategy only includes a UAG codon-rich reporter gene and an artificial anticodon-redesigned tRNA (tRNA_CUA_), thus having little impact on the cellular metabolism of host cells. The mRNA secondary structures of the reporter genes were not significantly influenced by the UAG substitution (Supplementary Fig. 9) ^59^. When the UAG codon encoded different amino acids of the Kan^R^, strains exhibited varied growth responses to kanamycin, individually (Fig. 2d and 2e). Herein, we proved that growth phenotypes could be fine-tuned in a UAG codon frequency-dependent manner. The reduced expression of UAG-rich reporter genes could be partially recovered through feeding or increased endogenous synthesis of the corresponding amino acid in amber-stop-codon-recoded or non-recoded microbial hosts. As a proof of concept, marker genes and derivatives were used to select amino acid overproducers, and overproduction strains of L-tryptophan, L-phenylalanine, L-aspartic acid, L-leucine, or L-glycine were screened out from mutation libraries of the corresponding hosts in a one-round screening. In addition, a strain producing over 20 g l^−1^ L-tryptophan in a fed-batch fermentation was de novo engineered based on the analysis of the genotype of selected mutants. Furthermore, the selection system could be potentially extended for screening high-efficient enzymes and bio-bricks for amino acid biosynthesis as well as amino acid analogs such as 4- and 5-fluorotryptophan ^60^ and L-β-(thieno[3,2-b]pyrrolyl)alanine ^61^ that are involved in the aminoacylation reaction.

Various transcription factor-based biosensors have been successfully established for screening amino acid overproducers. To date, only several amino acids such as L-arginine, L-histidine, L-lysine, L-methionine, L-leucine, L-isoleucine, L-valine, L-cysteine, and L-serine had their corresponding biosensor since this approach utilized the specific binding property of the biosensor toward the corresponding amino acid ^62,63^. Therefore, most of these biosensors are limited to a specific amino acid and depend on the RNA polymerase of a specific microbe. Recently, the protein translation-based selection systems based on the rare codon or mutant aaRS were sequentially developed to screen amino acid overproducers. For *E. coli*, ten amino acids have different competing tRNAs for the rare codon-based selection system ^64^ and only six amino acids have reported aaRS mutants with low activities ^65-70^, which limited the application of these methods.

A higher concentration of one substrate could raise the reduced rate of reaction caused by the modification of the other substrate in a two-substrate sequential reaction. Therefore, the mechanism could be used to construct a selection strategy for screening the overproducers of a specific substrate involved in a two-substrate sequential reaction. However, the applicability and fidelity of this selection strategy have not been proved in metabolic engineering. In this study, we demonstrated that an increased supply of amino acids could rescue the reduced rate of aminoacylation caused by modification of the critical structure of tRNA. This tRNA_CUA_-based amino acid screening system is a proof-of-concept for this selection strategy.

For the tRNA_CUA_-based system, if an aminoacylation reaction mainly depends on the specific anticodon recognition and any change in the anticodon might minimize or eliminate the recognition of its corresponding aaRS. For example, 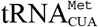 and 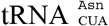 in *E. coli* were reported to be deprived of the original aminoacylation reaction ^71,72^. We also did not observe the growth rescue by adding 10 g l^-1^ L-Met and L-Asn in this study. However, a mutant metRS^W461AD456HQ211R^ has been successfully engineered to recognize the 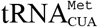 ^73^. Theoretically, coupling the mutant metRS with tRNA could be applied to screen the overproducers of L-methionine in the future. For 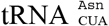, a modified tRNA with G73 mutation that led to lowered affinity to AsnRS could be used to construct a whole-cell-based selection system as reported ^18,74^. Furthermore, other codon/anticodon pairs such as UGA/tRNA_UCA_, UCG/tRNA_CGA_, and UCA/tRNA_UGA_ are alternatives to our selection system ^72,75,76^.CUA

When tRNA_CUA_ could be recognized by different aaRSs, for instance, tRNATrp could be recognized by TrpRS and GlnRS ^54,77^. Such phenomenon might affect the fidelity of the tRNA_CUA_-based system. However, we did not observe any fidelity issue in our cross-feeding experiments (Fig. 3f), probably because the incorporation of Gln in the corresponding Trp sites could destroy the function or folding of the expressed Kan^R^ protein. It was reported that positioning selenocysteine in the key functional^33^ position of the reporter protein could increase the fidelity of the selection system ^78^. Here, we added multiple UAG sites in the *kan*^*R*^ protein to minimize or even eliminate the possibility of producing functional Kan^R^ if misincorporation of Gln occurred at the Trp site. Our approach could eliminate the potential false-positive strains selected by the tRNA_CUA_-based system.

Increasing the efficiency of the tRNA_CUA_-based system could decrease the occurrence of potential false-positive strains by fine-tuning the number of UAG codons in the reporter genes and reducing the abundance of tRNA via controlling its promoter strength. Modification of tRNA anticodon decreased the aminoacylation rate in varying degrees, thereby resulting in different selection stringency (Supplementary Table 1) when the tRNA_CUA_-based system was applied to different amino acids. Other engineered tRNA_CUA_s, nonsense codons, and corresponding decoding tRNAs, as well as orthogonal aaRS-tRNA pairs, are alternatives for adjusting the efficiency of the selection system ^54,76,79^.

The transcriptome and genome analysis of L-tryptophan overproducers showed that the enhanced L-tryptophan biosynthesis was probably caused by the redirected flux and altered cofactor metabolism. The selection system could be utilized to obtain the amino acid overproducers ^80,81^. The industrial production strains, which have a high intracellular amino acid concentration that could saturate the selection system, are not suitable for direct screening via our strategy. However, the AMINO strategy could be applied to laboratory production strains to screen or select beneficial production mutations. By analyzing the omics information of the amino acid overproducers, useful clues for amino acid overproduction, such as beneficial mutations and enzymes, could be identified. The identified beneficial genes or mutations could then be used for de novo construction of an amino acid-overproducing strain or modification of hyper-producers of amino acid. Further modifications of the industrial overproduction strains, such as gene overexpression or gene deletion, are applicable through classic systematic metabolic engineering strategies. In the case of the down-regulated *ackA*, the repressed transcription was probably caused by a significant down-regulation of its activator Nac, which harbored a G to A mutation located right in the middle of the –12 element of its σ^54^-dependent promoter. Such mutation in the key promoter element could severely disrupt transcription initiation and thus lead to decreased expression of the downstream genes (Supplementary Tables 2 and 3). Hence, this method could break the bottleneck of rational design approaches, which are often restricted by the poor understanding of cell metabolisms and the lack of beneficial enzymes in targeted strains ^82^.

Taken together, we developed a tRNA modification-based strategy for identifying ami<u>n</u>o acid overproducers. Results showed that introducing UAG codons into the reporter genes along with overexpressing the artificial tRNA_CUA_ resulted in reduced translation efficiency, which could be restored through feeding or producing the corresponding amino acid. As a proof-of-concept work, *E. coli* strains overproducing three amino acids were successfully selected from random mutation libraries, and the selection system was also demonstrated to be suitable for other organisms such as *C. glutamicum*. This work provided a universal, convenient, and nontoxic strategy for high-throughput screening of amino acid overproducers. The mechanism behind this strategy might shed new light on the construction of a screening and selection system for overproducers of substrates involved in other two-substrate reactions.

## Material and methods

### Chemicals, strains, and culture conditions

All chemicals mentioned were purchased from Sinopharm Chemical Reagent Co. Ltd. (Beijing, China) if not otherwise specified. L-tryptophan, L-phenylalanine, L-aspartic acid, and antibiotics were purchased from Solarbio (Beijing, China). Oligonucleotides were synthesized by GENEWIZ Bio Inc. (Suzhou, China). The *E. coli* XL10-gold, MG1655 derivative (JW128), C321.ΔA, and the engineered strains used in this study were listed in Supplementary Table 4. LB medium consists of 5.0 g l^−1^ yeast extract, 10 g l^−1^ tryptone, and 10 g l^−1^ NaCl. Two-fold and Five-fold dilutions of LB medium were made to prepare the 0.5×LB and 0.2×LB medium. TB medium consists of 24 g l^−1^ yeast extract, 12 g l^−1^ tryptone, 5.0 g l^−1^ glycerol, 2.31 g l^−1^ KH_2_PO_4_, and 12.54 g l^−1^ K_2_HPO_4_. The LBHIS medium consists of 5.0 g l^−1^ yeast extract, 10 g l^−1^ tryptone, 10 g l^−1^ NaCl, and 10 g l^−1^ brain heart infusion. Modified M9 medium consists of 11.28 g l^− 1^ 5×M9 minimal salt, 40 g l^−1^ glucose, 1 mM MgSO_4_, 1 mM CaCl_2_, 10 μg ml^−1^ vitamin B_1_, and 1 g l^−1^ yeast extract. Amino acid fermentation was carried out using a modified M9 minimal growth medium (CGXII for *C. glutamicum*) and the fed-batch medium was used as described previously ^83^. For *E. coli*, fifty μg ml^−1^ of kanamycin, twenty-five μg ml^−1^ of chloramphenicol, or fifty μg ml^−1^ of spectinomycin were routinely supplemented in broth or solid media, respectively. Twenty-five μg ml^−1^ of kanamycin, fifteen μg ml^−1^ of chloramphenicol, or 80 μg ml^−1^ of spectinomycin were routinely supplemented in broth or solid media for *C. glutamicum*, respectively. The C321.ΔA and derivatives, as well as the *C. glutamicum* ATCC 13032 and derivatives, were generally cultivated at 30°C in a shaker at 200 rpm whereas other *E. coli* strains were cultivated at 37°C. PCR-based gene synthesis was applied to introduce the UAG codons into the *kan*^*R*^ gene.

### Plasmid construction

Gibson assembly technique was employed for the construction of all plasmids. The plasmids and primers used in this study are described in Supplementary Tables 4 and 5, respectively. The expression cassette of the wild-type *kan*^*R*^ including its promoter and terminator was amplified from pET-28a, which was cloned into the pKan backbone containing the *spec*^*R*^ gene. Detailed information on the *kan*^*R*^, *rfp*, and tagged *cm*^*R*^ gene harboring UAG codons are provided in Supplementary Tables 6, 7, and 8. The tRNA decoding of the corresponding amino acid was constructed by the replacement of the original tRNA anticodon to CUA (Supplementary Table 9), which was expressed under the control of a constitutive *lpp* promoter and the *rrnB* T1 terminator ^54^. To construct the selection system, these fragments of *kan*^*R*^-*UAG*s and *rfp-3UAG(W)* with the corresponding tRNA were synthesized and cloned into the pKan vector by the primers. To construct the pKan, pKan-4UAG(W)-delta, 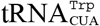, and pKan-4UAG(W), the fragments *kan*^*R*^-*4UAG(W)* and 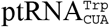were ligated into the pKan vector, respectively. Furthermore, the expression cassette of the wild-type *kan*^*R*^ was ligated with the expression cassette of 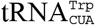, which was cloned into the pKan vector. The recombinant plasmids described above were transformed into the C321.ΔA strain. To construct the convenient system, the part of the UAG-rich NusA protein tag was ligated with the N-terminus of the *cm*^*R*^ gene by a flexible (GGGGS)_3_ linker. The genes were synthesized by ∼60 bp oligonucleotides with 20 bp overlaps ^84^.

### Construction of the tRNA_CUA_-based system

The pKan-UAG serial plasmids were transformed into C321.ΔA, XL10-gold, and MG1655 derivatives (JW128) for generating the corresponding strains, respectively. The correct transformants were checked by colony PCR. For each strain, three single colonies were picked from the agar plates and pre-cultured into a 5 ml LB medium supplemented with kanamycin (50 μg ml^−1^) overnight. The optical density at 600 nm was determined at a specified time point. One μl pre-culture was transferred into the 1 ml corresponding medium except 0.2×LB for obtaining OD_600_ values of 0.6×10^−3^ in a 2 ml 96-well deep well plate. Fifteen μl pre-culture was transferred into the 1 ml 0.2×LB medium for the growth test. C321.ΔA strain harboring pKan-5UAG(W), pKan-8UAG(F), or pKan-9UAG(D) was inoculated into an LB medium with different dilution factors (LB, 0.5×LB, or 0.2×LB), which was supplemented with L-tryptophan (2.0 g l^−1^ and 6.0 g l^−1^), 3AA (2.0 g l^−1^ L-tryptophan and L-phenylalanine, 0.3 g l^−1^ L-tyrosine, or 3 g l^−1^ L-tryptophan and L-phenylalanine, 0.3 g l^−1^ L-tyrosine), 2.0 g l^−1^ L-aspartic acid. Strains XL10-gold and JW128 harboring pKan-3UAG(W), pKan-5UAG(F), or pKan-7UAG(D) inoculated into TB or LB media, respectively, which were supplemented with 6.0 g l^−1^ L-tryptophan, 3AA (3 g l^−1^ L-tryptophan and L-phenylalanine, 0.3 g l^−1^ L-tyrosine), 6.0 g l^−1^ L-aspartic acid. JW128 strain harboring pKan-2UAG(3-MH), or pKan-2UAG(5-HTP) was inoculated into LB or 0.5×LB media which was supplemented with 3-methylhistidine (1 mM, 5 mM, and 10 mM), as well as 10 mM 5-hydroxytryptophan respectively. JW128 strain harboring pCm-2UAG(D), pCm-2UAG(C), pCm-2UAG(L), pCm-2UAG(E), pCm-2UAG(K), or pCm-2UAG(I) was inoculated into LB medium which was supplemented with 6.0 g l^−1^ L-aspartic acid, 6.0 g l^−1^ L-cysteine, 2.0 g l^−1^ L-leucine, 2.0 g l^−1^ L-glutamine, 2.0 g l^−1^ L-lysine, and 10.0 g l^−1^ L-isoleucine. *C. glutamicum* strains ATCC 13032 harboring pCm-1UAG(L) and pCm-1UAG(G) were inoculated into a 2×LBHIS medium, which was supplemented with 6.0 g l^−1^ L-leucine and 6.0 g l^−1^ L-glycine. Specifically, appropriate acids, bases, and MOPS were simultaneously used for pH adjustment since supplementation of L-aspartic acid, and L-lysine could influence the pH value of the medium. To monitor cell growth, a 200 μl culture was taken periodically into a 96-well microplate. The OD_600_ measurement was evaluated in triplicate at defined time points by a microplate reader.

### Mutant library construction

The mutant library of the C321.ΔA strain was created by NTG (N-methyl-N’-nitro-N-nitrosoguanidine) or UV treatment. A 50 μl of overnight C321.ΔA pre-culture was transferred into a 50 ml LB medium and cultured to the logarithmic phase (OD_600_ 0.4∼0.6). The cells were spun down and resuspended in 0.1 M (pH 5.5) sodium citrate solution containing 50 μg ml^−1^ NTG for 45, 60, and 75 minutes. After the treatment, mutants were washed thoroughly with 0.1 M (pH 7.1) nitrate solution and incubated in a shaker (200 rpm) at 30°C for 1 h. 100 μl of the culture was inoculated into a 5 ml LB medium and incubated at 200 rpm at 30°C until the OD_600_ reached 0.4∼0.6. For UV mutagenesis, 50 μl of overnight pre-cultured strains were transferred into a fresh medium and cultured to the logarithmic phase. 200 μl of cells were transferred to the center of the petri dish which was exposed to UV light for the appropriate treatment. A fatal of 97% was set as the threshold. The survival rate was calculated by spreading the mutant and control strains on the LB plate at each NTG and UV exposure time point for the quantification of survival colonies.

### Amino acid fermentation and measurement

To screen amino acid overproducers from mutation libraries, the mutant strains were made into competent cells and transformed with the corresponding plasmids for the selection of targeted amino acid overproducers. The strains with different genomic large-fragment deletions were made into competent cells and transformed with pKan-3UAG(W) and pGFP-2UAG(W) for the selection of L-tryptophan overproducers, respectively. The transformants were picked and cultured in the corresponding medium with the appropriate markers, respectively. Rapid-growing cells and high GFP/OD_600_ were screened out and then inoculated into the corresponding medium, respectively. The large fragment mutant library and the reference strain were transformed with the pKan-3UAG(W). For fed-batch cultivations, 100 ml seed cultures of TP10 strain were cultivated to a 3 L fermenter (Parallel-bioreactor, China) containing 1000 ml fed-batch medium at 32 C. Base (30% (v/v) NH_4_OH) was added to control pH at 6.7∼7.0. The initial airflow was 2 L min^−1^ and the stirring speed was set to 500∼1000 rpm min^−1^ for maintaining the dissolved oxygen level above 30%. To quantify the extracellular concentration of L-tryptophan, L-phenylalanine, L-aspartic acid, L-leucine, and L-glycine, the cell cultures were centrifuged at 12,000 rpm for 5 min, and the supernatant was analyzed using HPLC equipped with a UV detector and an Agilent UHPLC-C18 column (1.8 μm×250 mm, 2.1 μm) according to a previously described protocol ^85^. The concentrations of L-tryptophan, L-phenylalanine, L-aspartic acid, L-leucine, and L-glycine were calculated through a calibration curve obtained with the standard solution of L-tryptophan, L-phenylalanine, L-aspartic acid, L-leucine, and L-glycine, respectively.

### RNA extraction and transcriptome analysis

C321. ΔA strain and the selected L-tryptophan overproducers (TP2 and TP4) were cultivated and harvested at the logarithmic phase in a 0.2×LB medium. Total RNAs were isolated using Transzol reagent (Transgen) and redissolved in 10 mM sodium acetate (pH 4.5). The RNA sequencing was performed on cells from the wild-type C321.ΔA (the reference strain) and mutants (TP2 and TP4) which were cultured in a 0.2×LB medium and harvested in the exponential phase. Clean reads of 21.62, 20.21, and 20.45 million were obtained from TP2, TP4, and reference strains, respectively. Reverse transcription was performed by using PrimeScript™ RT Master Mix. Quantitative RT-PCR was performed with QuantStudio 3 or 5 Real-Time PCR Systems using TB Green^®^ Premix Ex Taq™ GC (Perfect Real Time), Takara Biomedical Technology (Beijing).

The housekeeping gene *gyrA* was used as a reference gene as it tends to maintain steady expression under various growth conditions ^86^. The cycling condition was as follows: 30s denaturation at 95°C, followed by 40 cycles of 5s at 94°C, 55°C for 30 s, and 72°C for 25 s. Melting curve analysis was performed by raising the temperature from 60 to 95°C at a rate of 0.1°C s^−1^, with five signal acquisitions per degree. Data were acquired from three biological replicates and each sample was measured in duplicate.

### Genome sequencing

The genomic DNA was extracted from cells harvested at the logarithmic growth phase in a 0.2×LB medium. The genomic DNA samples of the mutant strains TP2 and TP4 were sequenced by Illumina HiSeq/Nova 2×150 bp platform, producing approximately 20 million clean reads. Raw reads containing more than 20 low-quality bases were filtered out and the clean reads were generated using Cutadapt (v1.9.1). Comparative genomic analysis including the SNV and InDel annotations was analyzed using BWA (version 0.7.17).

## Acknowledgments

The work completed at the Beijing Institute of Technology was supported by the National Key R&D Program of China (Grant No. 2019YFA0904104), the National Natural Science Foundation of China (Grant No. 32000059), and the Fundamental Research Funds for the Central Universities. We also thank the Biological and Medical Engineering Core Facilities of the Beijing Institute of Technology for supporting several experimental equipments.

## Author contributions

Y.X.H. and X.M. generated the idea. Y.X.H., X.M., H.G., and N.W. designed the project. H.G., N.W., T.D., B.Z., L.G., C.H., W. Z., and L.S. carried out the experiments. H.G., X.M., and Y.X.H. analyzed the data. H.G., L.S., X.M., and Y.X.H, wrote the manuscript.

## Competing interests

The authors declare no competing interests.

## Data availability

Data associated with this project can be found at the NCBI under BioProject PRJNA751045. The transcriptomic data of the wild-type, TP2, and TP4 strains can be found under BioSample SRR15305620, SRR15305619, and SRR15305618, respectively. The genomic data of the TP2 and TP4 strains can be found under BioSample SRR15305617 and SRR15305616. The authors declare that the data are available from the authors upon request. The link to the reviewers is https://www.ncbi.nlm.nih.gov/bioproject/?term=PRJNA751045.

## Notes

### Competing Interest Statement

The authors have declared no competing interest.

