## Supplementary materials for "A tRNA Modification-based strategy for Identifying amiNo acid Overproducers (AMINO)"

Guo *et al*.

**This file includes:**

Supplementary Note 1

Supplementary Figs. 1 to 9

Supplementary Tables 1 to 9

References (1 to 39)

**Supplementary** **Note 1**

Aminoacylation reaction catalyzed by aminoacyl-tRNA synthetases except for GlnRS, GluRS, ArgRS, or LysRS I, can be simplified to describe as follows (1-4):

$E+S_{1}+\mathrm{ATP}\overset{K_{1}}{\leftrightharpoons}E\cdot S_{1}-AMP+PPi$	(1)

$E\cdot S_{1}-\mathrm{AMP}+S_{2}\overset{K_{2}}{\leftrightharpoons}E+P+\mathrm{AMP}$	(2)

*E* is the enzyme, *S*_1_ is the amino acid (aa), *S*_2_ is the tRNA, $E\cdot S_{1}-$AMP is the aaRS-aminoacyl-adenylate complex and *P* is the product. When tRNA and amino acid were considered the two major substrates, we assumed that the aminoacylation reaction could be further simplified as follow:

$E+S_{1}+S_{2}\overset{K}{\to}E+P$	(3)

According to the Michaelis-Menten equation, the aminoacylation rate is

$v=\frac{v_{max}\left[ S_{1} \right]\left[ S_{2} \right]}{K_{m}^{S_{1}}\left[ S_{2} \right]+K_{m}^{S_{2}}\left[ S_{1} \right]+\left[ S_{1} \right]\left[ S_{2} \right]}$	(4)

It follows that,

$\frac{1}{v}=\frac{K_{m}^{S_{1}}}{v_{max}}\frac{1}{\left[ S_{1} \right]}+\frac{K_{m}^{S_{2}}}{v_{max}}\frac{1}{\left[ S_{2} \right]}+\frac{1}{\left[ v_{max} \right]}$ (5)

We should mention that some aaRSs such as GlnRS, GluRS, ArgRS, or LysRS I first bind the tRNA instead of its cognate amino acid. However, the different binding sequences did not influence the total aminoacylation rate.

Based on our assumption that the parameters $K_{m}^{S_{1}}, v_{max}$, and$\left[ S_{2} \right]$ keep constant in the aminoacylation reaction, the following equation is used:

$y=\frac{A}{\left[ S_{1} \right]}+\frac{K_{m}^{S_{2}}}{B}+C$ (6)

where y represents the reciprocal of the aminoacylation rate, A is defined as $\frac{K_{m}^{S_{1}}}{v_{max}}$, B is defined as $\frac{1}{\left[ S_{2} \right]v_{max}}$, and C is defined as $\frac{1}{\left[ v_{max} \right]}$.

If $K_{m}^{S_{2}}$ increases by modifying the anticodon of tRNA to CUA, a decrease of $\frac{A}{\left[ S_{1} \right]}$, e.g., an increase of $\left[ S_{1} \right]$, is a way to keep the reciprocal of the aminoacylation rate $y$ constant.

**Supplementary Table 1** Effects of the modifications in the anticodon of tRNAs on aminoacylation properties in *E*. *coli*.

| tRNAs that cannot bind to aaRS directly but to aaRS-aa complex | | | | | |
| --- | --- | --- | --- | --- | --- |
| aaRS | tRNA | Mutation in anticodon | The tendency of change in catalytic efficiency | Maximum Fold Change | Ref |
| HisRS | tRNAHis GUG | GUG **→** CUA | relative fluorescence of the GFP | ~1/2 | 5 |
| PheRS | tRNAPhe GAA | GAA **→** CUU | kinetics of dipeptide bond | ~1/60 | 6 |
| AlaRS | tRNAAla GGC/tRNAAla UGC | GGC/UGC **→** CUA | *K_m_* acceptor activity | ~1/3 | 7,8 |
| CysRS | tRNACys GCA | GCA **→** CUA | *K_m_* *k_cat_* *k_cat_ /K_m_* | ~1/3100 | 9 |
| GlyRS | tRNAGly GCC | GCC **→** CUA | *V_max_*/*K_m_* | ~1/41 | 10,11 |
| AspRS | tRNAAsp GUC | GUC **→** CUA | *K_m_ k_cat_ /K_m_* | ~1/3.4×10^6^ | 12 |
| LysRS | tRNALys UUU | UUU **→** UUG | *K_m_ V_max_* *V_max_ /K_m_* | ~1/4.55 | 13 |
| LeuRS | tRNALeu CAG | CAG **→** CUA | *k_cat_* | ~1/10 | 14 |
| MetRS | tRNAMet CAU | CAU **→** CUA | acceptor activity | ~1/100 | 15 |
| AsnRS | tRNAAsn GUU | GUU **→** CUU | aminoacylation rate | N/A | 16 |
| SerRS | tRNASer CGA | CGA **→** CUA | *K_m_* *k_cat_* *k_cat_/K_m_*  acceptor activity | ~1/3 | 11,17 |
| TyrRS | tRNATyr GUA | GUA **→** CUA | relative fluorescence of the GFP | ~1/30 | 18 |
| ThrRS | tRNAThr GGU | GGU **→** GNN | *V_max_*/*K_m_* | ~1/10000 | 19 |
| IleRS | tRNAIle CAU | CAU **→** NAU | aminoacylation rate | ~1/10 | 20 |
| TrpRS | tRNATrp CCA | CCA **→** CUA | *K_m_* | ~1/60 | 21 |
| ProRS | tRNAPro UGG | UGG **→** CUA | *k_cat_/K_m_* | ~1/150 | 22 |
| ValRS | tRNAVal UAC | UAC **→** UAA | *K_m_* *k_cat_* *k_cat_/K_m_* | ~1/5800 | 23 |
| tRNAs that can bind to aaRS independent of amino acid | | | | | |
| ArgRS | tRNAArg ACG | ACG **→** CUA | *K_m_* *V_max_* *V_max_ /K_m_* | ~1/2 | 13,24 |
| GlnRS | tRNAGln CUG | CUG **→** CUA | *K_m_* *k_cat_* *k_cat_ /K_m_* | ~1/40 | 25 |
| GluRS | tRNAGlu UUC | UUC **→** UUG | *K_m_* *k_cat_* *k_cat_ /K_m_* | ~1/105 | 26 |

**Supplementary Table 2.** Related regulations and genetic mutations regarding up- and down-regulated genes.

**Supplementary Table 3.** The detailed information on mutation positions in the genome of the mutant strains TP2 and TP4.

**Supplementary Table 4.** Strains and plasmids used in this study.

**Supplementary Table 5.** Primers used in this study.

**Supplementary Table 6.** The detailed information of *kan^R^*-*UAG*s in the tRNA_CUA_-based selection system.

**Supplementary Table 7.** The detailed information of pKan-UAGs in the tRNA_CUA_-based selection system.

**Supplementary Table 8.** The sequences of the *kan^R^*, *rfp*, *gfp*, and tagged *cm^R^* gene harboring UAG codons

**Supplementary Table 9.** The sequences of the tRNA decoding the UAG codon

**Supplementary Figures**

**a**

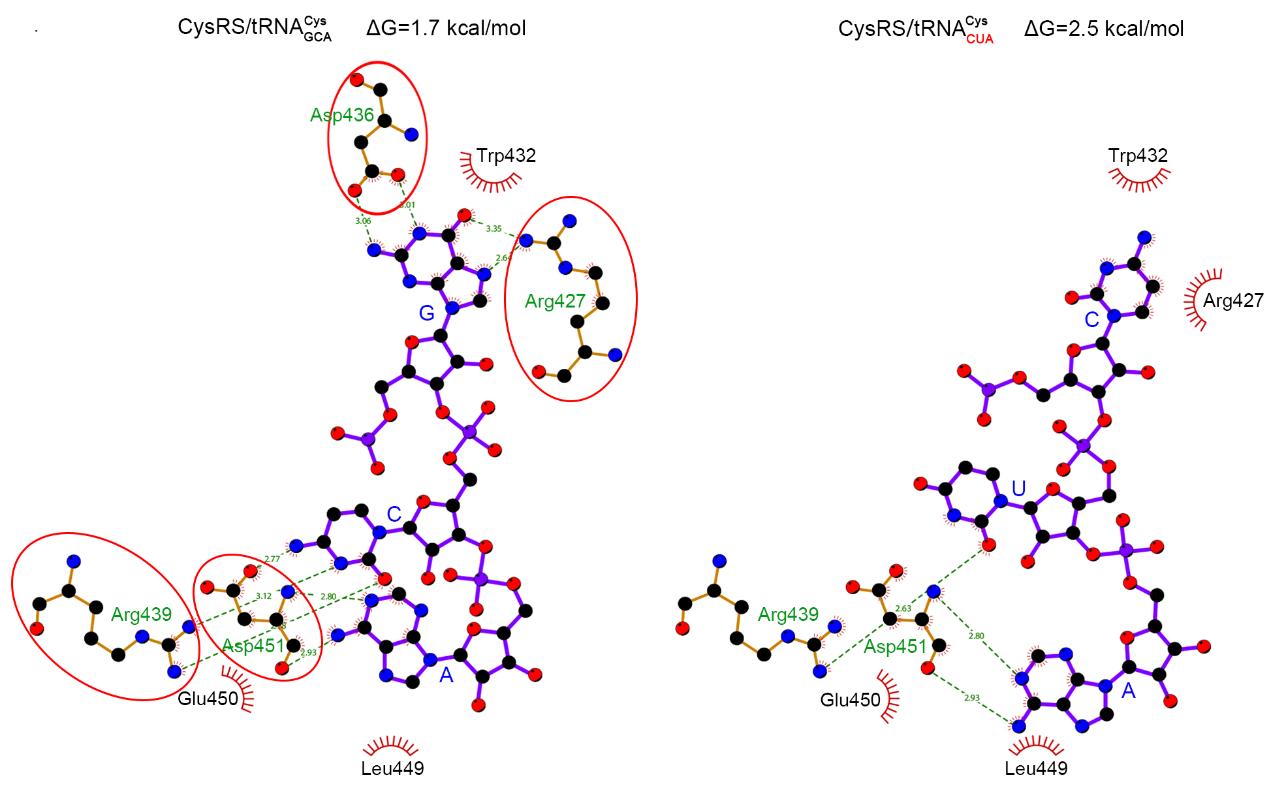

**b**

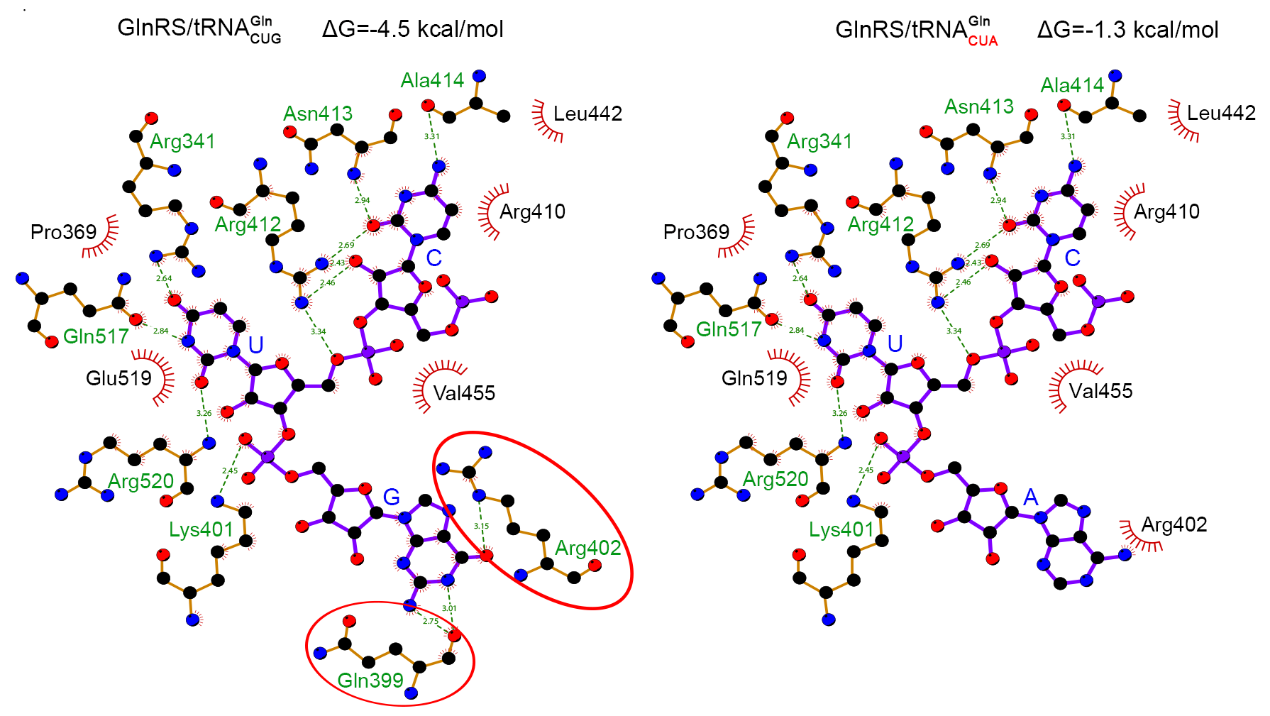

**Supplementary Figure 1.** The role of tRNA anticodon in the interaction between aaRS and tRNA in tRNA–aaRS complexes. **a**. The detailed interaction of CysRS residues with tRNACys GCA (left) and tRNACys CUA (right). The complex structure of CysRS-tRNA_GCA_ (PDB: 1U0B) was used to generate the structure of CysRS-tRNA_CUA_ by modifying tRNA anticodon nucleotides to CUA. **b**. The detailed interaction of GlnRS residues with tRNAGln CUG (left) and tRNAGln CUA (right). The complex structure of GlnRS-tRNA_CUG_ (PDB: 1O0B) was used to generate the structure of GlnRS-tRNA_CUA_ by modifying tRNA anticodon nucleotides to CUA. The interaction between aaRS and the anticodon region was investigated using Ligplot software. The model with purple and bold bonds represents the tRNA anticodon base. The tRNA anticodon nucleotides were labeled in blue uppercase letters. The model with yellow and thin bonds depicts the amino acid residues of aaRS that interact with the tRNA anticodon. The hydrophilic and hydrophobic residues were labeled in green and black, respectively. The interaction between the tRNA anticodon and the hydrophilic residues was indicated in green lines and the distance was labeled using numbers in green colors. The amino acid residues with red spokes pointing toward tRNA anticodon represent the residues involved in hydrophobic interactions. The carbon atoms, oxygen atoms, and nitrogen atoms were indicated in black, red, and blue, respectively. The binding affinity of aaRS and tRNA was calculated using the PDBePISA tool as reflected by the solvation-free energy (ΔG).

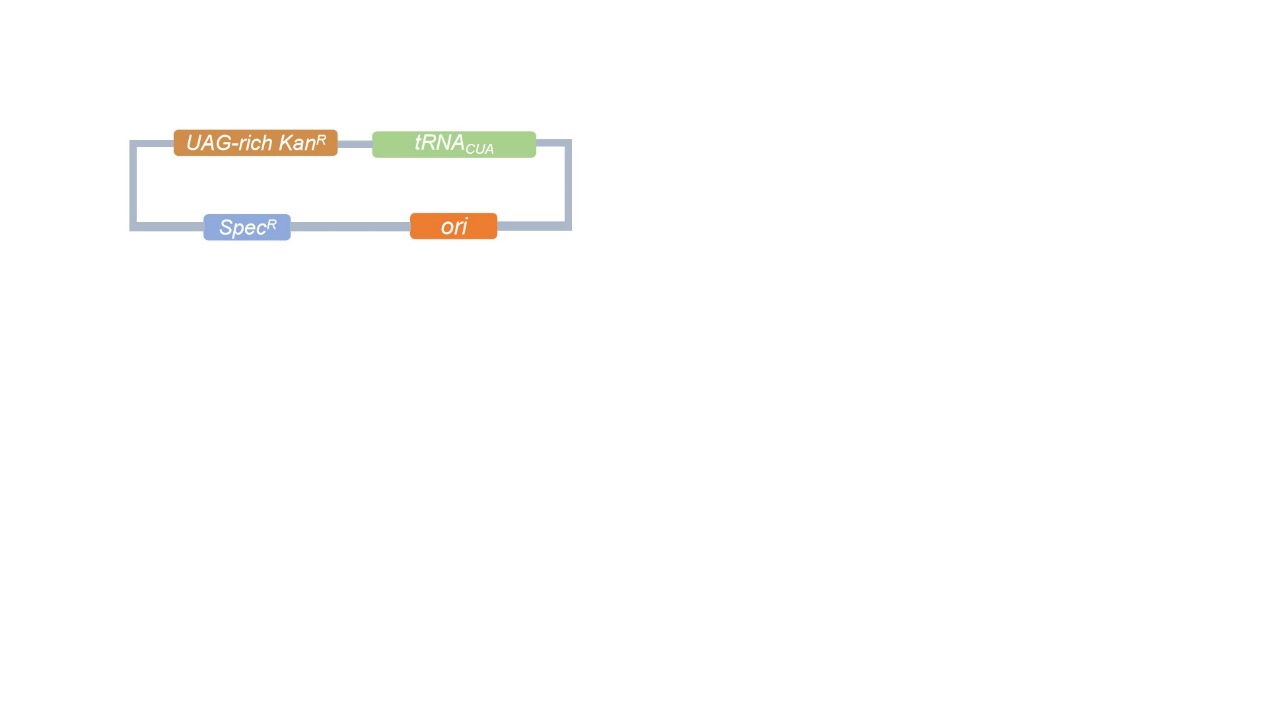

**Supplementary Figure 2.** The plasmid map for the construction of the *kan^R^*-based selection system

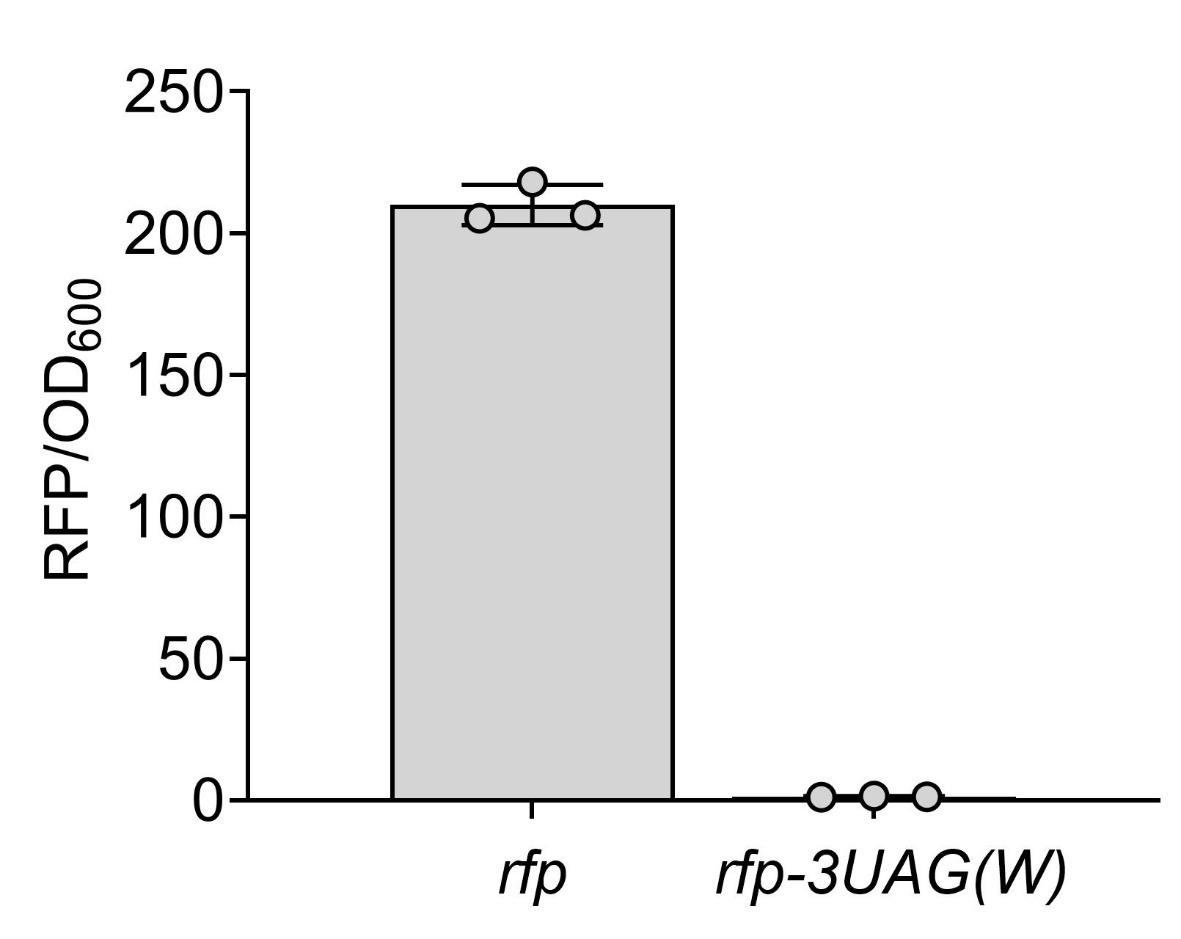

**Supplementary Figure 3**. Effects of the incorporation of UAG codons on the RFP expression in LB medium. Values and error bars represent the mean and the s.d. (n = 3).

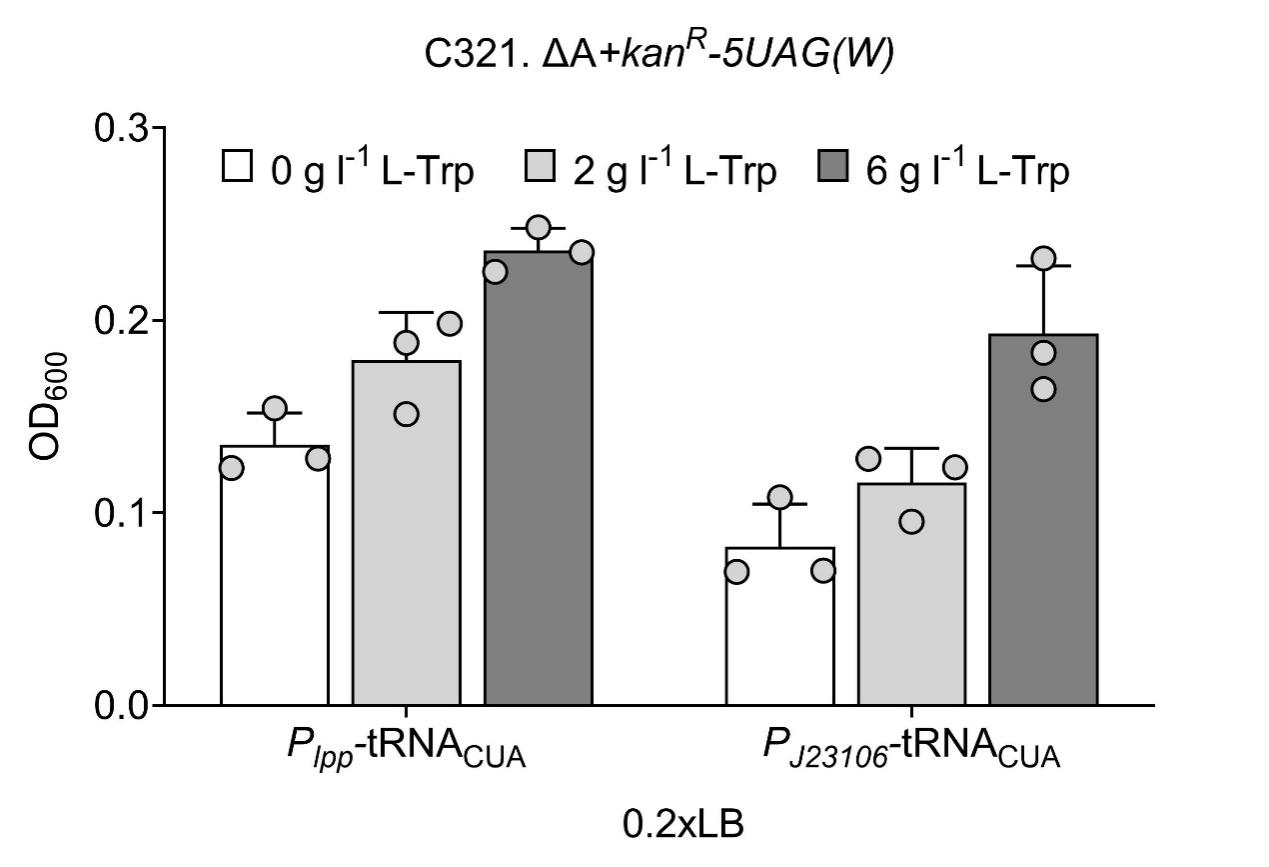

**Supplementary Figure 4**. Selection stringency mediated by promoter strength. Effects of the weak promoter *P_J23106_* of tRNATrp CUAon the cell OD_600_ for C321. ΔA strains harboring the pKan-5UAG(W) gene. Values and error bars represent the mean and the s.d. (n = 3).

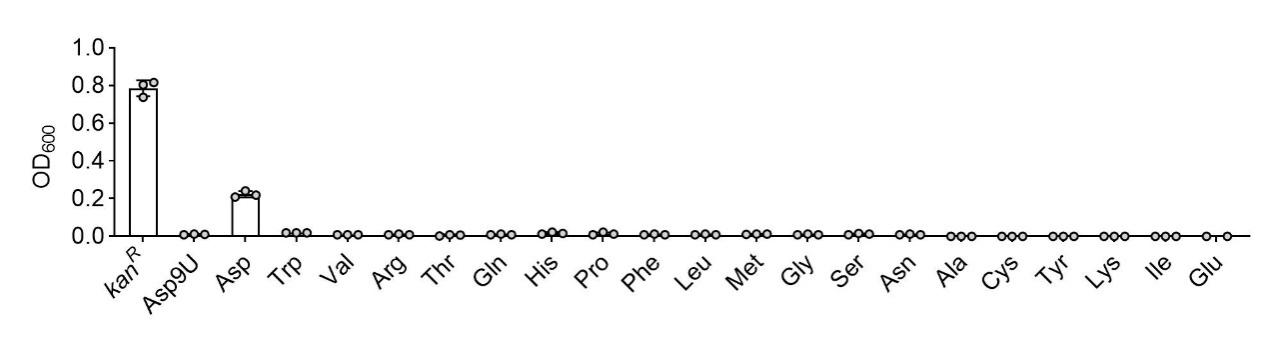

**Supplementary Figure 5**. Changes in cell growth of C321.ΔA strain harboring pKan-9UAG(D) after feeding 2 g l^–1^ 20 proteinogenic amino acids in LB medium. Values and error bars represent the mean and the s.d. (n = 3).

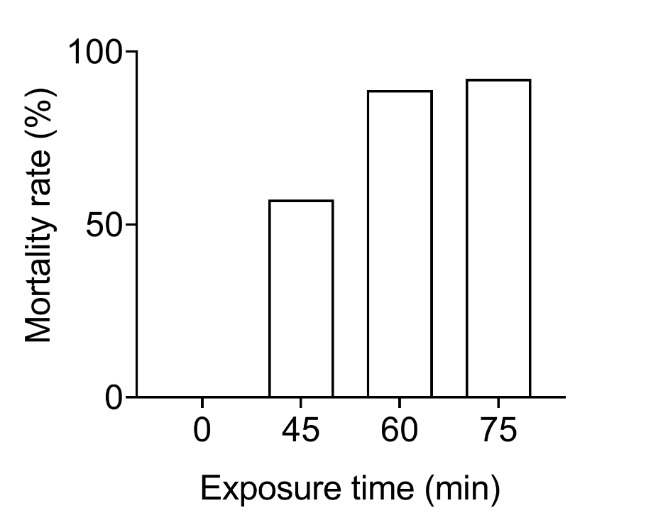

**Supplementary Figure 6.** Effects of different NTG treatment times on the mortality rate.

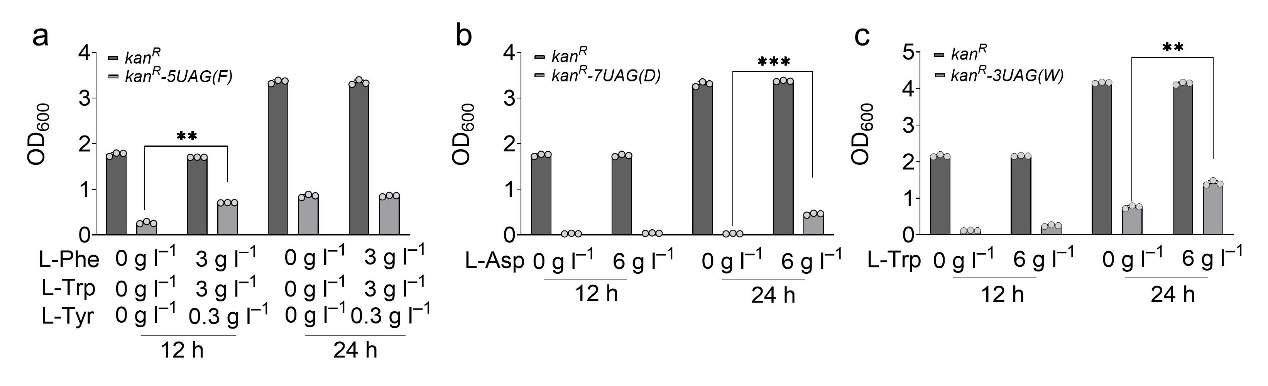

**Supplementary Figure 7**. Cell growth of *E*. *coli* JW128 (MG1655 derivative) harboring **a** pKan-5UAG(F), **b** pKan-7UAG(D) in LB medium and LB medium supplemented with the corresponding amino acid. *E*. *coli* JW128 harboring **c** pKan-3UAG(W) in TB medium and TB medium supplemented with 6 g l^–1^ L-tryptophan. Values and error bars represent the mean and the s.d. (n = 3, ***P* < 0.01, ****P* < 0.001 as determined by two-tailed t-test).

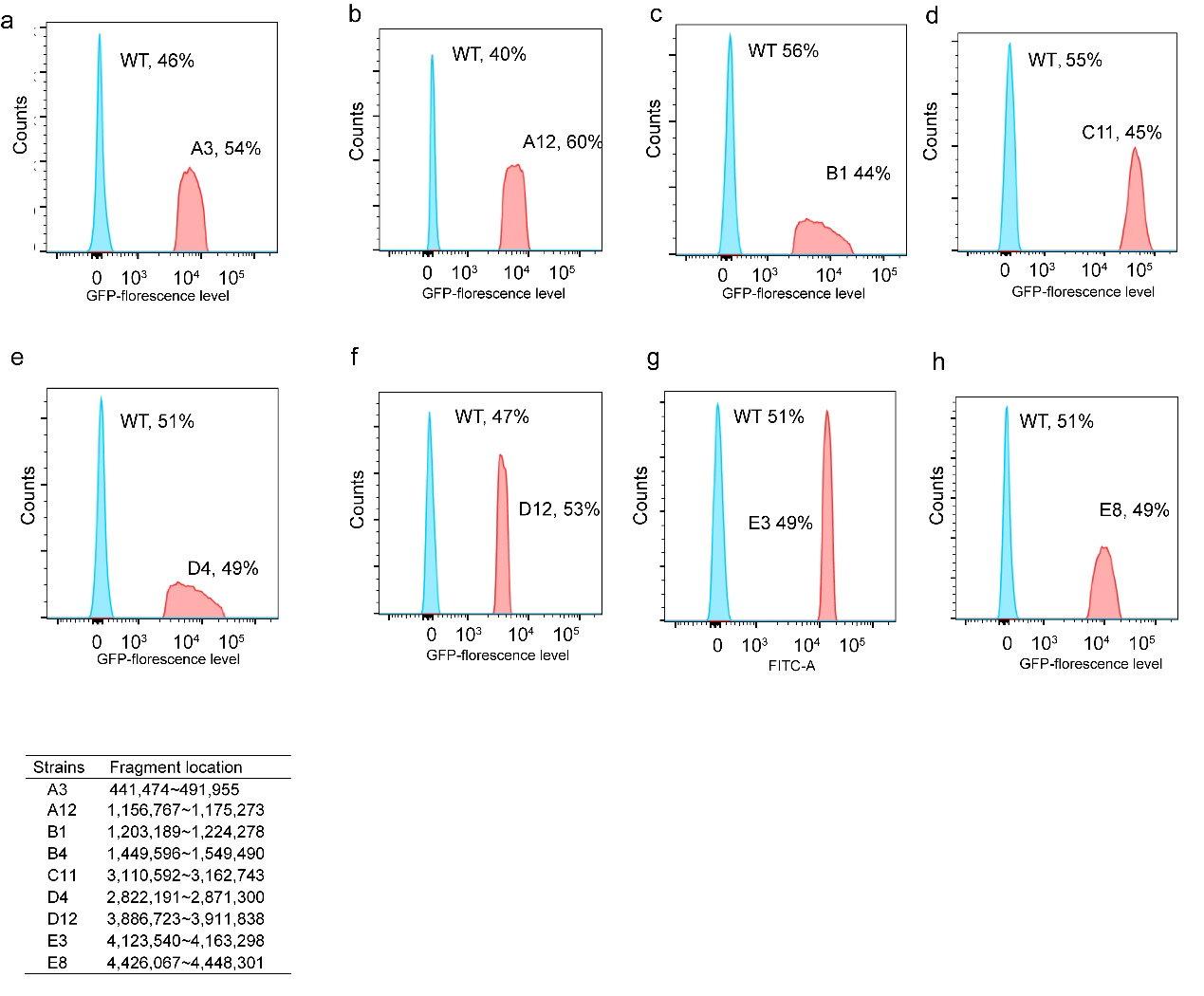

i

**Supplementary Figure 8.** Fluorescence-activated cell sorter sorting of the mixture containing wild-type strain and mutant strains (**a** A3, **b** A8, **c** B1, **d** C11, **e** D4, **f** D12, **g** E3, and, **h** E8) exhibiting high GFP/OD_600_ from 65 MG1655 derivative strains with different genomic large-fragment deletions (**i**). The red spots represent the mutant strains whereas the blue ones represent the wild-type strain.

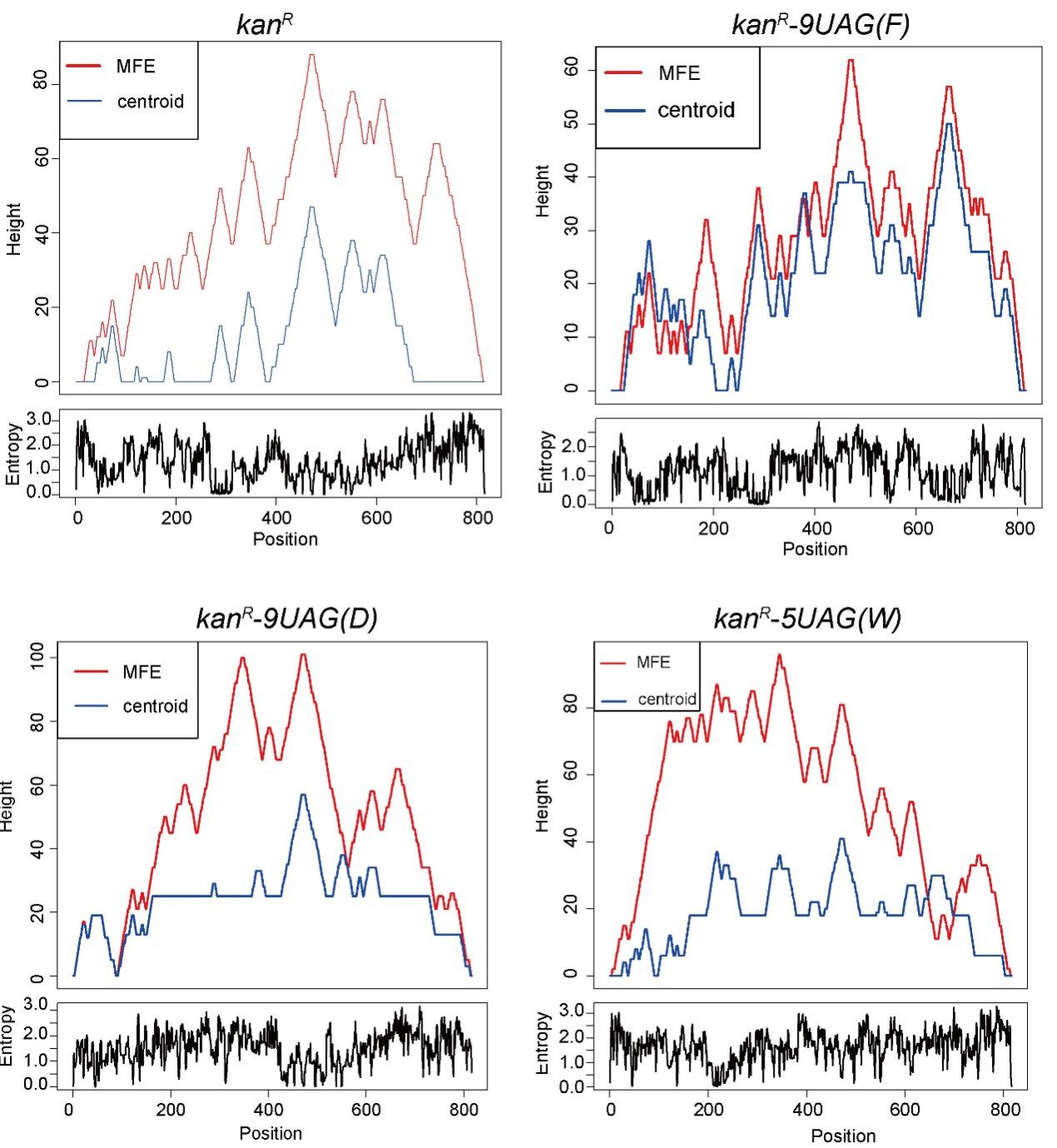

**Supplementary Figure 9.** Potential secondary structures of the mRNA transcribed from the wild-type *kan^R^*, *kan^R^-9UAG(F)*, *kan^R^-9UAG(D)*, and *kan^R^-5UAG(W)*. The red line represents the minimum free energy (MFE) structure. The blue line represents the centroid structure. The heights represent the number of potential base pairs in a defined position, where plateaus, peaks, and slopes represent loops, hairpin loops, and helices, respectively. The positional entropy for each position is presented in the lower boxes.

**Supplementary Table 2.** Related regulations and genetic mutations regarding up- and down-regulated genes.

| **Genes** | **Related regulations and genetic mutations** | **Refs** |
| --- | --- | --- |
| **Down-regulated genes** | | |
| *ackA* | A 12-fold down-regulation of its activator Nac, of which the promoter had a G to A mutation located in the center of the -12 element of Nac σ^54^-dependent promoter. | 27 |
| *pykF* | A 1.5-fold up-regulation of its repressor Cra. | 28 |
| *ptsG* | A 2.4-fold down-regulation of its activator Crp; A 4.9-fold up-regulation of its repressor SgrS, an RNA that also promotes degradation of the Crp mRNA. | 29,30 |
| *tnaA* | A 2.4-fold and a 1.4-fold down-regulation of its activator Crp and TorR, respectively; a 1.8-fold up-regulation of its repressor NarP, as well as a 2.5-fold increase of the regulator Fnr that facilitates *tnaA* repression. | 31-33 |
| *zwf* | A 1.5-fold down-regulation of its activator Rob; a 1.5-fold up-regulation of its repressor Cra. | 34,35 |
| **Up-regulated genes** | | |
| *ppsA* | A 22-fold and 1.5-fold up-regulations of its activator PdhR and Cra, respectively. | 36,37 |
| *aroF* | A 1.7-fold down-regulation of its repressor TyrR; a 2.2-fold up-regulation of its activator SoxR. | 38,39 |

**Supplementary Table 3.** The detailed information on mutation positions in the genome of the mutant strains TP2 and TP4.

| **Mutant strain** | **Position in the genome** | **Reference** | **Mutant** | **Location** |
| --- | --- | --- | --- | --- |
| TP2 | 10984 | G | A | exonic |
| TP2 | 23181 | G | A | exonic |
| TP2 | 43318 | G | A | exonic |
| TP2 | 44263 | G | A | exonic |
| TP2 | 46390 | G | A | exonic |
| TP2 | 50940 | G | A | exonic |
| TP2 | 62916 | G | A | exonic |
| TP2 | 108515 | G | A | exonic |
| TP2 | 115401 | G | A | exonic |
| TP2 | 120406 | G | A | exonic |
| TP2 | 158173 | G | A | exonic |
| TP2 | 166622 | G | A | exonic |
| TP2 | 203631 | G | A | exonic |
| TP2 | 234491 | G | A | exonic |
| TP2 | 239524 | G | A | exonic |
| TP2 | 246579 | G | A | exonic |
| TP2 | 264888 | T | C | exonic |
| TP2 | 283492 | G | A | exonic |
| TP2 | 519365 | A | G | exonic |
| TP2 | 602171 | C | T | exonic |
| TP2 | 612914 | C | T | exonic |
| TP2 | 916196 | G | A | exonic |
| TP2 | 1106424 | G | A | exonic |
| TP2 | 1111636 | G | A | exonic |
| TP2 | 1113956 | G | A | exonic |
| TP2 | 1124229 | G | A | exonic |
| TP2 | 1127623 | G | A | exonic |
| TP2 | 1128028 | G | A | exonic |
| TP2 | 1130116 | A | G | exonic |
| TP2 | 1132880 | G | A | exonic |
| TP2 | 1136897 | G | A | exonic |
| TP2 | 1139297 | G | A | exonic |
| TP2 | 1143892 | G | A | exonic |
| TP2 | 1144006 | G | A | exonic |
| TP2 | 1145103 | G | A | exonic |
| TP2 | 1150600 | G | A | exonic |
| TP2 | 1156358 | G | A | exonic |
| TP2 | 1156403 | G | A | exonic |
| TP2 | 1165415 | G | A | exonic |
| TP2 | 1196399 | C | T | exonic |
| TP2 | 1201451 | C | T | exonic |
| TP2 | 1237428 | C | T | exonic |
| TP2 | 1249859 | C | T | exonic |
| TP2 | 1399597 | C | T | exonic |
| TP2 | 1530307 | T | C | exonic |
| TP2 | 1701751 | T | C | exonic |
| TP2 | 1845069 | G | A | exonic |
| TP2 | 1883670 | G | A | exonic |
| TP2 | 1889777 | G | A | exonic |
| TP2 | 1916121 | G | A | exonic |
| TP2 | 1924486 | G | A | exonic |
| TP2 | 1927096 | G | A | exonic |
| TP2 | 1929097 | G | A | exonic |
| TP2 | 1957699 | G | A | exonic |
| TP2 | 1963936 | G | A | exonic |
| TP2 | 1966190 | G | A | exonic |
| TP2 | 1989469 | G | A | exonic |
| TP2 | 1996798 | G | A | exonic |
| TP2 | 2009656 | G | A | exonic |
| TP2 | 2014864 | G | A | exonic |
| TP2 | 2020248 | G | A | exonic |
| TP2 | 2022206 | G | A | exonic |
| TP2 | 2036868 | G | A | exonic |
| TP2 | 2158225 | G | A | exonic |
| TP2 | 2236743 | T | C | exonic |
| TP2 | 2272437 | C | T | exonic |
| TP2 | 2275266 | C | T | exonic |
| TP2 | 2280311 | C | T | exonic |
| TP2 | 2398651 | C | T | exonic |
| TP2 | 2408609 | C | T | exonic |
| TP2 | 2419434 | C | T | exonic |
| TP2 | 2421882 | C | T | exonic |
| TP2 | 2426130 | C | T | exonic |
| TP2 | 2427973 | C | T | exonic |
| TP2 | 2449668 | C | T | exonic |
| TP2 | 2454328 | C | T | exonic |
| TP2 | 2468470 | C | T | exonic |
| TP2 | 2479430 | T | C | exonic |
| TP2 | 2484955 | C | T | exonic |
| TP2 | 2485903 | C | T | exonic |
| TP2 | 2495176 | C | T | exonic |
| TP2 | 2699476 | C | T | exonic |
| TP2 | 2849609 | T | C | exonic |
| TP2 | 3027726 | G | A | exonic |
| TP2 | 3052058 | C | T | exonic |
| TP2 | 3070079 | C | T | exonic |
| TP2 | 3078922 | C | T | exonic |
| TP2 | 3122210 | C | T | exonic |
| TP2 | 3124917 | C | T | exonic |
| TP2 | 3125100 | T | C | exonic |
| TP2 | 3125823 | C | T | exonic |
| TP2 | 3129048 | C | T | exonic |
| TP2 | 3205493 | G | A | exonic |
| TP2 | 3486279 | C | T | exonic |
| TP2 | 3584376 | A | C | exonic |
| TP2 | 3631192 | G | A | exonic |
| TP2 | 3835523 | G | A | exonic |
| TP2 | 3857374 | C | T | exonic |
| TP2 | 3923176 | A | G | exonic |
| TP2 | 3939856 | G | A | exonic |
| TP2 | 4215613 | A | G | exonic |
| TP2 | 4414738 | C | T | exonic |
| TP2 | 4509716 | A | G | exonic |
| TP2 | 4535261 | A | G | exonic |
| TP4 | 215658 | G | A | exonic |
| TP4 | 264888 | T | C | exonic |
| TP4 | 452183 | C | T | exonic |
| TP4 | 551653 | C | T | exonic |
| TP4 | 584603 | A | G | exonic |
| TP4 | 916196 | G | A | exonic |
| TP4 | 1128870 | G | A | exonic |
| TP4 | 1134006 | G | A | exonic |
| TP4 | 1148750 | G | A | exonic |
| TP4 | 1158071 | G | A | exonic |
| TP4 | 1159862 | G | A | exonic |
| TP4 | 1174791 | G | A | exonic |
| TP4 | 1176946 | G | A | exonic |
| TP4 | 1193403 | G | A | exonic |
| TP4 | 1194331 | G | A | exonic |
| TP4 | 1989038 | C | T | exonic |
| TP4 | 2236743 | T | C | exonic |
| TP4 | 2287109 | C | T | exonic |
| TP4 | 2479430 | T | C | exonic |
| TP4 | 2870036 | G | A | exonic |
| TP4 | 3125100 | T | C | exonic |
| TP4 | 3486279 | C | T | exonic |
| TP4 | 3584376 | A | C | exonic |
| TP4 | 3828255 | G | A | exonic |
| TP4 | 3856260 | G | A | exonic |
| TP4 | 3923176 | A | G | exonic |
| TP4 | 3927489 | G | A | exonic |
| TP4 | 3937630 | G | A | exonic |
| TP4 | 4125813 | C | T | exonic |
| TP4 | 4164846 | C | T | exonic |
| TP4 | 4215613 | A | G | exonic |
| TP4 | 4277427 | C | T | exonic |
| TP4 | 4414738 | C | T | exonic |
| TP4 | 4509716 | A | G | exonic |

| **Supplementary Table 4.** Strains and plasmids used in this study | | |
| --- | --- | --- |
| **Name** | **Description** | **Reference** |
| C321.ΔA | *E. coli* MG1655 derivative, *Δ*(*ybhB-bioAB*)::[*lcI857 N(cro-ea59)::tet^R^-bla*] *ΔprfA* Δ*mutS*::*zeo^R^*; all 321 UAG codons changed to UAA | Addgene |
| XL10-gold | *E. coli* Tet^R^Δ*(mcrA)183* Δ*(mcrCB-hsdSMR-mrr)173 endA1 supE44 thi-1 recA1 gyrA96 relA1 lac Hte* [F´ *proAB lacI^q^Z*Δ*M15* Tn*10 (*Tet^R^*)* Amy Cam^R^]. | Lab stock |
| JW128 | MG1655 derivative, *ΔaraBAD::tet^R^*, *ΔmcrCB*-*hsdSMR*-*mrr*, *mcrA*^–^, *endA*^–^, *recA*^–^ | Lab stock |
| ATCC13032 | *C. glutamicum* wild type | ATCC |
| A3 | MG1655 derivative, deletion of fragment location (441,474~491,955) on the MG1655 genome (accession No. U00096) | This study |
| A12 | MG1655 derivative, deletion of fragment location (1,156,767~1,175,273) on the MG1655 genome (accession No. U00096) | This study |
| B1 | MG1655 derivative, deletion of fragment location (1,203,189~1,224,278) on the MG1655 genome (accession No. U00096) | This study |
| B4 | MG1655 derivative, deletion of fragment location (1,449,596~1,549,490) on the MG1655 genome (accession No. U00096) | This study |
| C11 | MG1655 derivative, deletion of fragment location (2,822,191~2,871,300) on the MG1655 genome (accession No. U00096) | This study |
| D4 | MG1655 derivative, deletion of fragment location (3,110,592~3,162,743) on the MG1655 genome (accession No. U00096) | This study |
| D12 | MG1655 derivative, deletion of fragment location (3,886,723~3,911,838) on the MG1655 genome (accession No. U00096) | This study |
| E3 | MG1655 derivative, deletion of fragment location (4,123,540~4,163,298) on the MG1655 genome (accession No. U00096) | This study |
| E8 | MG1655 derivative, deletion of fragment location (4,426,067~4,448,301) on the MG1655 genome | This study |
| TP1 | C321.ΔA L-tryptophan overproducer | This study |
| TP2 | C321.ΔA L-tryptophan overproducer | This study |
| TP3 | C321.ΔA L-tryptophan overproducer | This study |
| TP4 | C321.ΔA L-tryptophan overproducer | This study |
| TP5 | C321.ΔA L-tryptophan overproducer | This study |
| TP6 | C321.ΔA L-tryptophan overproducer | This study |
| TP7 | C321.ΔA L-tryptophan overproducer | This study |
| TP10 | JW128 derivative, L-tryptophan overproducer  *trpR*^–^, *tnaAB*^–^, *mtr*^–^, *pta*^–^, *P_T5_*: *aroG^fbr^*, *trpE^fbr^*, *trpEDCBA*, *serA^fbr^* with pTrp | This study |
| PP1 | C321.ΔA L-phenylalanine overproducer | This study |
| PP2 | C321.ΔA L-phenylalanine overproducer | This study |
| PP3 | C321.ΔA L-phenylalanine overproducer | This study |
| PP4 | C321.ΔA L-phenylalanine overproducer | This study |
| PP5 | C321.ΔA L-phenylalanine overproducer | This study |
| PP6 | C321.ΔA L-phenylalanine overproducer | This study |
| PP7 | C321.ΔA L-phenylalanine overproducer | This study |
| PP8 | C321.ΔA L-phenylalanine overproducer | This study |
| AP1 | C321.ΔA L-aspartic acid overproducer | This study |
| AP2 | C321.ΔA L-aspartic acid overproducer | This study |
| AP3 | C321.ΔA L-aspartic acid overproducer | This study |
| AP4 | C321.ΔA L-aspartic acid overproducer | This study |
| AP5 | C321.ΔA L-aspartic acid overproducer | This study |
| AP6 | C321.ΔA L-aspartic acid overproducer | This study |
| AP7 | C321.ΔA L-aspartic acid overproducer | This study |
| AP8 | C321.ΔA L-aspartic acid overproducer | This study |
| LP1 | ATCC13032 L-leucine overproducer | This study |
| LP2 | ATCC13032 L-leucine overproducer | This study |
| LP3 | ATCC13032 L-leucine overproducer | This study |
| LP4 | ATCC13032 L-leucine overproducer | This study |
| LP5 | ATCC13032 L-leucine overproducer | This study |
| LP6 | ATCC13032 L-leucine overproducer | This study |
| LP7 | ATCC13032 L-leucine overproducer | This study |
| LP8 | ATCC13032 L-leucine overproducer | This study |
| GP1 | ATCC13032 L-glycine overproducer | This study |
| GP2 | ATCC13032 L- glycine overproducer | This study |
| GP3 | ATCC13032 L-glycine overproducer | This study |
| GP4 | ATCC13032 L-glycine overproducer | This study |
| GP5 | ATCC13032 L-glycine overproducer | This study |
| GP6 | ATCC13032 L-glycine overproducer | This study |
| GP7 | ATCC13032 L-glycine overproducer | This study |
| GP8 | ATCC13032 L-glycine overproducer | This study |
| *Plasmids* | | |
| pET-28a | *kan^R^;* pBR322 origin | Lab stock |
| pACP | iGEM Part: BBa_K592009 | Lab stock |
| pTSpurple | iGEM Part: BBa_K1033906 | Lab stock |
| pKan | *kan^R^*; s*pec^R^*; p15A origin | Lab stock |
| pSpec | *spec^R^*; p15A origin |  |
| pRFP | *rfp*; *spec^R^*; p15A origin | Lab stock |
| pNG2 | *spec^R^*; NG2 origin | Lab stock |
| pGFP | *gfp*; *kan^R^*; p15A origin | Lab stock |
| pAmp | *amp^R^*; *kan^R^*; ColE1 origin | Lab stock |
| pTrp | *cm^R^*, *P_tac_:trpEDCBA, aroG, serA,* ColE1 origin | This study |
| pCm | *kan^R^*, *cm^R^*, adding the partial NusA tag and a flexible linker (GGGGS) on the N-terminal of *cm^R^* | This study |
| pGFP-2UAG(W) | pGFP derivative, replacing 2 trp codons of *gfp* with UAG, *P_lpp_*-*tRNATrp CUA*-*T_rrnB T1_* | This study |
| pCm-1UAG | pKan-Cm-Tag derivative, replacing 1 codon of the partial NusA tag with UAG | This study |
| pCm-2UAG | pKan-Cm-Tag derivative, replacing 2 codons of the partial NusA tag with UAG | This study |
| pCm-2UAG(D) | pCm-2UAG derivative, *P_lpp_*-*tRNAAsp CUA* | This study |
| pCm-2UAG(C) | pCm-2UAG derivative, *P_lpp_*-*tRNACys CUA* | This study |
| pCm-2UAG(L) | pCm-2UAG derivative, *P_lpp_*-*tRNALeu CUA* | This study |
| pCm-2UAG(E) | pCm-2UAG derivative, *P_lpp_*-*tRNAGlu CUA* | This study |
| pCm-2UAG(K) | pCm-2UAG derivative, *P_lpp_*-*tRNALys CUA* | This study |
| pCm-2UAG(I) | pCm-2UAG derivative, *P_lpp_*-*tRNAIle CUA* | This study |
| pCm-1UAG(L) | pCm-1UAG derivative, *P_gapA_*-*tRNALeu CUA* | This study |
| pCm-1UAG(G) | pCm-1UAG derivative, *P_gapA_*-*tRNAGly CUA* | This study |
| pKan-5HTP | pAmp derivative, *P_tac_*-WRS1-*T_Luxl_, P_LeuV_* -*ScWtRNA40A*-*T_rrnB T1_* | This study |
| pKan-1UAG(5-HTP) | pAmp-5HTP derivative, replacing 1 Trp codon of *kan^R^* with UAG, *P_tac_*-WRS1-*T_Luxl_, P_LeuV_* -*ScWtRNA40A*-*T_rrnB T1_* | This study |
| pKan-2UAG(5-HTP) | pAmp-5HTP derivative, replacing 2 Trp codons of *kan^R^* with UAG, *P_tac_*-WRS1-*T_Luxl_, P_LeuV_* -*ScWtRNA40A*-*T_rrnB T1_* | This study |
| pKan-3UAG(5-HTP) | pAmp-5HTP derivative, replacing 3 Trp codons of *kan^R^* with UAG, *P_tac_*-WRS1-*T_Luxl_, P_LeuV_* -*ScWtRNA40A*-*T_rrnB T1_* | This study |
| pKan-3MH | pAmp derivative, *P_lpp_*-*pylRS*-*T_Luxl_, P_LeuV_* -*pyltRNA*-*T_rrnB T1_* | This study |
| pKan-1UAG(3-MH) | pKan-3MH derivative, replacing 1 His codon of *kan^R^* with UAG, *P_lpp_*-*pylRS*-*T_Luxl_, P_LeuV_* -*pyltRNA*-*T_rrnB T1_* | This study |
| pKan-2UAG(3-MH) | pKan-3MH derivative, replacing 2 His codons of *kan^R^* with UAG, *P_lpp_*-*pylRS*-*T_Luxl_, P_LeuV_* -*pyltRNA*-*T_rrnB T1_* | This study |
| pKan-2UAG(3-MH) | pKan-3MH derivative, replacing 2 His codons of *kan^R^* with UAG, *P_lpp_*-*pylRS*-*T_Luxl_, P_LeuV_* -*pyltRNA(G73T)*-*T_rrnB T1_* | This study |
| pKan-5UAG(3-MH) | pKan-3MH derivative, replacing 3 His codons of *kan^R^* with UAG, *P_lpp_*-*pylRS*-*T_Luxl_, P_LeuV_* -*pyltRNA*-*T_rrnB T1_* | This study |
| pKan-3UAG(W) | pKan derivative, replacing 3 Trp codons of *kan^R^* with UAG, *P_lpp_*-*tRNA*Trp CUA-*T_rrnB T1_* | This study |
| pKan-4UAG(W) | pKan derivative, replacing 4 Trp codons of *kan^R^* with UAG, *P_lpp_*-*tRNA*Trp CUA-*T_rrnB T1_* | This study |
| pKan-5UAG(W) | pKan derivative, replacing 5 Trp codons of *kan^R^* with UAG, *P_lpp_*-*tRNATrp CUA*-*T_rrnB T1_* | This study |
| pKan-6UAG(W) | pKan derivative, replacing 6 Trp codons of *kan^R^* with UAG, *P_lpp_*-*tRNATrp CUA*-*T_rrnB T1_* | This study |
| pKan-10UAG(S) | pKan derivative, replacing 10 Ser codons of *kan^R^* with UAG, *P_lpp_*-*tRNASer CUA*-*T_rrnB T1_* | This study |
| pKan-5UAG(H) | pKan derivative, replacing 5 His codons of *kan^R^* with UAG, *P_lpp_*-*tRNAHis CUA*-*T_rrnB T1_* | This study |
| pKan-8UAG(I) | pKan derivative, replacing 8 Ile codons of *kan^R^* with UAG, *P_lpp_*-*tRNAIle CUA*-*T_rrnB T1_* | This study |
| pKan-7UAG(Q) | pKan derivative, replacing 7 Gln codons of *kan^R^* with UAG, *P_lpp_*-*tRNAGln CUA*-*T_rrnB T1_* | This study |
| pKan-10UAG(R) | pKan derivative, replacing 10 Arg codons of *kan^R^* with UAG, *P_lpp_*-*tRNAArg CUA*-*T_rrnB T1_* | This study |
| pKan-8UAG(E) | pKan derivative, replacing 8 Glu codons of *kan^R^* with UAG, *P_lpp_*-*tRNAGlu CUA*-*T_rrnB T1_* | This study |
| pKan-7UAG(T) | pKan derivative, replacing 7 Thr codons of *kan^R^* with UAG, *P_lpp_*-*tRNAThr CUA*-*T_rrnB T1_* | This study |
| pKan-4UAG(C) | pKan derivative, replacing 4 Cys codons of *kan^R^* with UAG, *P_lpp_*-*tRNACys CUA*-*T_rrnB T1_* | This study |
| pKan-10UAG(P) | pKan derivative, replacing 10 Pro codons of *kan^R^* with UAG, *P_lpp_*-*tRNAPro CUA*-*T_rrnB T1_* | This study |
| pKan-20UAG(L) | pKan derivative, replacing 20 Leu codons of *kan^R^* with UAG, *P_lpp_*-*tRNALeu CUA*-*T_rrnB T1_* | This study |
| pKan-10UAG(N) | pKan derivative, replacing 10 Asn codons of *kan^R^* with UAG, *P_lpp_*-*tRNAAsn CUA*-*T_rrnB T1_* | This study |
| pKan-4UAG(M) | pKan derivative, replacing 4 Met codons of *kan^R^* with UAG, *P_lpp_*-*tRNAMet CUA*-*T_rrnB T1_* | This study |
| pKan-5UAG(Y) | pKan derivative, replacing 5 Tyr codons of *kan^R^* with UAG, *P_lpp_*-*tRNATyr CUA*-*T_rrnB T1_* | This study |
| pKan-10UAG(A) | pKan derivative, replacing 10 Ala codons of *kan^R^* with UAG, *P_lpp_*-*tRNAAla CUA*-*T_rrnB T1_* | This study |
| pKan-10UAG(G) | pKan derivative, replacing 10 Gly codons of *kan^R^* with UAG, *P_lpp_*-*tRNAGly CUA*-*T_rrnB T1_* | This study |
| pKan-10UAG(V) | pKan derivative, replacing 10 Val codons of *kan^R^* with UAG, *P_lpp_*-*tRNAVal CUA*-*T_rrnB T1_* | This study |
| pKan-8UAG(K) | pKan derivative, replacing 8 Lys codons of *kan^R^* with UAG, *P_lpp_*-*tRNALys CUA*-*T_rrnB T1_* | This study |
| pKan-10UAG(F) | pKan derivative, replacing 10 Phe codons of *kan^R^* with UAG, *P_lpp_*-*tRNAPhe CUA*-*T_rrnB T1_* | This study |
| pKan-16UAG(D) | pKan derivative, replacing 16 Asp codons of *kan^R^* with UAG, *P_lpp_*-*tRNAAsp CUA*-*T_rrnB T1_* | This study |
| pKan-9UAG(F) | pKan derivative, replacing 9 Phe codons of *kan^R^* with UAG, *P_lpp_*-*tRNAPhe CUA*-*T_rrnB T1_* | This study |
| pKan-8UAG(F) | pKan derivative, replacing 8 Phe codons of *kan^R^* with UAG, *P_lpp_*-*tRNAPhe CUA*-*T_rrnB T1_* | This study |
| pKan-7UAG(F) | pKan derivative, replacing 7 Phe codons of *kan^R^* with UAG, *P_lpp_*-*tRNAPhe CUA*-*T_rrnB T1_* | This study |
| pKan-6UAG(F) | pKan derivative, replacing 6 Phe codons of *kan^R^* with UAG, *P_lpp_*-*tRNAPhe CUA*-*T_rrnB T1_* | This study |
| pKan-5UAG(F) | pKan derivative, replacing 5 Phe codons of *kan^R^* with UAG, *P_lpp_*-*tRNAPhe CUA*-*T_rrnB T1_* | This study |
| pKan-15UAG(D) | pKan derivative, replacing 15 Asp codons of *kan^R^* with UAG, *P_lpp_*-*tRNAAsp CUA*-*T_rrnB T1_* | This study |
| pKan-13UAG(D) | pKan derivative, replacing 13 Asp codons of *kan^R^* with UAG, *P_lpp_*-*tRNAAsp CUA*-*T_rrnB T1_* | This study |
| pKan-9UAG(D) | pKan derivative, replacing 9 Asp codons of *kan^R^* with UAG, *P_lpp_*-*tRNAAsp CUA*-*T_rrnB T1_* | This study |
| pKan-7UAG(D) | pKan derivative, replacing 7 Asp codons of *kan^R^* with UAG, *P_lpp_*-*tRNAAsp CUA*-*T_rrnB T1_* | This study |
| pKan-4UAG(W)-*delta* | pKan derivative, replacing 4 Trp codons of *kan^R^* with UAG | This study |
| pKan-13UAG(R) | pKan derivative, replaced 13 Arg codons of *kan^R^* by UAG, *P_lpp_*-*tRNAArg CUA*-*T_rrnB T1_* | This study |
| pKan-5UAG(A) | pKan derivative, replaced 5 Ala codons of *kan^R^* by UAG, *P_lpp_*-*tRNAAla CUA*-*T_rrnB T1_* | This study |
| pKan-3UAG(H) | pKan derivative, replaced 3 His codons of *kan^R^* by UAG, *P_lpp_*-*tRNAHis CUA*-*T_rrnB T1_* | This study |
| pKan-2UAG(I) | pKan derivative, replaced 2 Ile codons of *kan^R^* by UAG, *P_lpp_*-*tRNAIle CUA*-*T_rrnB T1_* | This study |
| pKan-2UAG(M) | pKan derivative, replaced 2 Met codons of *kan^R^* by UAG, *P_lpp_*-*tRNAMet CUA*-*T_rrnB T1_* | This study |
| pKan-6UAG(N) | pKan derivative, replaced 6 Asn codons of *kan^R^* by UAG, *P_lpp_*-*tRNAAsn CUA*-*T_rrnB T1_* | This study |
| pKan-9UAG(Q) | pKan derivative, replaced 9 Gln codons of *kan^R^* by UAG, *P_lpp_*-*tRNAGln CUA*-*T_rrnB T1_* | This study |
| pKan-7UAG(Y) | pKan derivative, replaced 7 Tyr codons of *kan^R^* by UAG, *P_lpp_*-*tRNATyr CUA*-*T_rrnB T1_* | This study |
| pKan-5UAG(V) | pKan derivative, replaced 5 Val codons of *kan^R^* by UAG, *P_lpp_*-*tRNAVal CUA*-*T_rrnB T1_* | This study |
| pKan-4UAG(T) | pKan derivative, replacing 4 Thr codons of *kan^R^* with UAG, *P_lpp_*-*tRNAThr CUA*-*T_rrnB T1_* | This study |
| ptRNATrp CUA | *spec^R^*; p15A origin, *P_lpp_-tRNATrp CUA-T_rrnB T1_* | This study |
| pRFP-3UAG(W) | pRFP derivative, *rfp,* replacing 3 Trp codons of rfp with UAG, *P_lpp_*-*tRNATrp CUA*-*T_rrnB T1_* | This study |
| pKan-5UAG (W)*-P_J23106_-tRNA_CUA_* | pKan derivative, replacing 5 Trp codons of *kan^R^* with UAG, *P_J23106_*-*tRNA*Trp CUA-*T_rrnB T1_* | This study |
| pGFP-Tag | pGFP derivative, the addition of the partial NusA tag at the N-terminus of GFP | This study |
| pGFP-2UAG(W) | pGFP-Tag derivative, replacing 2 Trp codons of the partial NusA tag with UAG | This study |

| **Supplementary Table 5.** Primers of qRT-PCR used in this study | |
| --- | --- |
| **Primers** | **Sequence 5’-3’** |
| gyrA-fw | GGCGAAGACGAAGTAATGCT |
| gyrA-rv | TTTATCGCCTTCACCTAAGCG |
| aroF-fw | CACGCAGCAAGGCATCGGTCAT |
| aroF-rv | GATCGAAAGTAATATCCACGAGGG |
| sthA-fw | CCGCCACCTTTCTGTTCCATAA |
| sthA-rv | TCGTCGGCATGAACGTGGGCA |
| pykA-fw | CGTTAGGCCCAGCAACAGATC |
| pykA-rv | TTATCCGCGCGCATTTTGTGATC |
| ackA-fw | GCGGTAGTTCTTCACTGAAATTTG |
| ackA-rv | GCCTGCACCTAAAGCCGCTTC |
| hisB-fw | CCCTGATTAGCGAACCGCCGA |
| hisB-rv | GTTCCAAGACCATCCTGATTAGTG |
| tnaA-fw | GAGCCAGTAAAACGTACCACTC |
| tnaA-rv | AGCCTGCATGCTCTGCGTCA |

**Supplementary Table 6.** The detailed information of *kan^R^*-*UAG*s in the tRNA_CUA_-based selection system.

| ***kan^R^*-*UAG*** | **Amino acid** | **The number of UAG codons** | **The total number of original codons** | **The proportion of UAG codons in total sense codons** |
| --- | --- | --- | --- | --- |
| *kan^R^*-*6UAG(W)* | L-tryptophan | 6 | 6 | 100.0% |
| *kan^R^*-*5UAG(W)* | L-tryptophan | 5 | 6 | 83.3% |
| *kan^R^*-*4UAG(W)* | L-tryptophan | 4 | 6 | 66.7% |
| *kan^R^*-*3UAG(W)* | L-tryptophan | 3 | 6 | 50.0% |
| *kan^R^*-*10UAG(S)* | L-serine | 10 | 16 | 62.5% |
| *kan^R^*-*5UAG(H)* | L-histidine | 5 | 7 | 71.4% |
| *kan^R^*-*3UAG(H)* | L-histidine | 3 | 7 | 42.8% |
| *kan^R^*-*8UAG(I)* | L-isoleucine | 8 | 13 | 61.5% |
| *kan^R^*-*2UAG(I)* | L-isoleucine | 2 | 13 | 15.4% |
| *kan^R^*-*7UAG(Q)* | L-glutamine | 7 | 10 | 70.0% |
| *kan^R^*-*9UAG(Q)* | L-glutamine | 9 | 10 | 90.0% |
| *kan^R^*-*13UAG(R)* | L-arginine | 13 | 16 | 81.25% |
| *kan^R^*-*10UAG(R)* | L-arginine | 10 | 16 | 62.5% |
| *kan^R^*-*8UAG(E)* | L-glutamic acid | 8 | 13 | 61.5% |
| *kan^R^*-*7UAG(T)* | L-threonine | 7 | 10 | 70.0% |
| *kan^R^*-*4UAG(T)* | L-threonine | 4 | 10 | 40.0% |
| *kan^R^*-*4UAG(C)* | L-cysteine | 4 | 5 | 80.0% |
| *kan^R^*-*10UAG(P)* | L-proline | 10 | 15 | 66.7% |
| *kan^R^*-*20UAG(L)* | L-leucine | 20 | 29 | 68.9% |
| *kan^R^*-*10UAG(N)* | L-asparagine | 10 | 15 | 66.7% |
| *kan^R^*-*6UAG(N)* | L-asparagine | 6 | 15 | 40.0% |
| *kan^R^*-*4UAG(M)* | L-methionine | 4 | 8 | 50.0% |
| *kan^R^*-*2UAG(M)* | L-methionine | 2 | 8 | 25.0% |
| *kan^R^*-*7UAG(Y)* | L-tyrosine | 7 | 7 | 100% |
| *kan^R^*-*5UAG(Y)* | L-tyrosine | 5 | 7 | 71.4% |
| *kan^R^*-*10UAG(A)* | L-alanine | 10 | 15 | 66.7% |
| *kan^R^*-*5UAG(A)* | L-alanine | 5 | 15 | 33.3% |
| *kan^R^*-*10UAG(G)* | L-glycine | 10 | 17 | 58.8% |
| *kan^R^*-*10UAG(V)* | L-valine | 10 | 16 | 62.5% |
| *kan^R^*-*5UAG(V)* | L-valine | 5 | 16 | 31.2% |
| *kan^R^*-*8UAG(K)* | L-lysine | 8 | 12 | 66.7% |
| *kan^R^*-*10UAG(F)* | L-phenylalanine | 10 | 16 | 66.7% |
| *kan^R^*-*9UAG(F)* | L-phenylalanine | 9 | 16 | 56.3% |
| *kan^R^*-*8UAG(F)* | L-phenylalanine | 8 | 16 | 50.0% |
| *kan^R^*-*7UAG(F)* | L-phenylalanine | 7 | 16 | 43.8% |
| *kan^R^*-*6UAG(F)* | L-phenylalanine | 6 | 16 | 37.5% |
| *kan^R^*-*5UAG(F)* | L-phenylalanine | 5 | 16 | 31.3% |
| *kan^R^*-*16UAG(D)* | L-aspartic acid | 16 | 25 | 64.0% |
| *kan^R^*-*15UAG(D)* | L-aspartic acid | 15 | 25 | 60.0% |
| *kan^R^*-*13UAG(D)* | L-aspartic acid | 13 | 25 | 52.0% |
| *kan^R^*-*9UAG(D)* | L-aspartic acid | 9 | 25 | 36.0% |
| *kan^R^*-*7UAG(D)* | L-aspartic acid | 7 | 25 | 28.0% |

**Supplementary Table 7.** The detailed information of pKan-UAGs in the tRNA_CUA_-based selection system.

| **Plasmids (pKan-UAGs)** | ***kan^R^*-*UAG*s** | **Mutated tRNAs and/or aaRS** | **Original tRNAs** |
| --- | --- | --- | --- |
| pKan-6UAG(W) | *kan^R^*-*6UAG(W)* | tRNATrp CUA | tRNATrp CCA |
| pKan-5UAG(W) | *kan^R^*-*5UAG(W)* | tRNATrp CUA | tRNATrp CCA |
| pKan-4UAG(W) | *kan^R^*-*4UAG(W)* | tRNATrp CUA | tRNATrp CCA |
| pKan-3UAG(W) | *kan^R^*-*3UAG(W)* | tRNATrp CUA | tRNATrp CCA |
| pKan-10UAG(S) | *kan^R^*-*10UAG(S)* | tRNASer CUA | tRNASer GCU |
| pKan-5UAG(H) | *kan^R^*-*5UAG(H)* | tRNAHis CUA | tRNAHis GUG |
| pKan-3UAG(H) | *kan^R^*-*3UAG(H)* | tRNAHis CUA | tRNAHis GUG |
| pKan-8UAG(I) | *kan^R^*-*8UAG(I)* | tRNAIle CUA | tRNAIle GAU |
| pKan-2UAG(I) | *kan^R^*-*2UAG(I)* | tRNAIle CUA | tRNAIle GAU |
| pKan-9UAG(Q) | *kan^R^*-*9UAG(Q)* | tRNAGln CUA | tRNAGln CUG |
| pKan-7UAG(Q) | *kan^R^*-*7UAG(Q)* | tRNAGln CUA | tRNAGln CUG |
| pKan-13UAG(R) | *kan^R^*-*13UAG(R)* | tRNAArg CUA | tRNAArg ACG |
| pKan-10UAG(R) | *kan^R^*-*10UAG(R)* | tRNAArg CUA | tRNAArg ACG |
| pKan-8UAG(E) | *kan^R^*-*8UAG(E)* | tRNAGlu CUA | tRNAGlu UUC |
| pKan-7UAG(T) | *kan^R^*-*7UAG(T)* | tRNAThr CUA | tRNAThr GGU |
| pKan-4UAG(T) | *kan^R^*-*4UAG(T)* | tRNAThr CUA | tRNAThr GGU |
| pKan-4UAG(C) | *kan^R^*-*4UAG(C)* | tRNACys CUA | tRNACys GCA |
| pKan-10UAG(P) | *kan^R^*-*10UAG(P)* | tRNAPro CUA | tRNAPro CGG |
| pKan-20UAG(L) | *kan^R^*-*20UAG(L)* | tRNALeu CUA | tRNALeu CAG |
| pKan-10UAG(N) | *kan^R^*-*10UAG(N)* | tRNAAsn CUA | tRNAAsn GUU |
| pKan-6UAG(N) | *kan^R^*-*6UAG(N)* | tRNAAsn CUA | tRNAAsn GUU |
| pKan-4UAG(M) | *kan^R^*-*4UAG(M)* | tRNAMet CUA | tRNAMet CAU |
| pKan-2UAG(M) | *kan^R^*-*2UAG(M)* | tRNAMet CUA | tRNAMet CAU |
| pKan-7UAG(Y) | *kan^R^*-*7UAG(Y)* | tRNATyr CUA | tRNATyr GUA |
| pKan-5UAG(Y) | *kan^R^*-*5UAG(Y)* | tRNATyr CUA | tRNATyr GUA |
| pKan-10UAG(A) | *kan^R^*-*10UAG(A)* | tRNAAla CUA | tRNAAla GGC |
| pKan-5UAG(A) | *kan^R^*-*5UAG(A)* | tRNAAla CUA | tRNAAla GGC |
| pKan-10UAG(G) | *kan^R^*-*10UAG(G)* | tRNAGly CUA | tRNAGly GCC |
| pKan-10UAG(V) | *kan^R^*-*10UAG(V)* | tRNAVal CUA | tRNAVal GAC |
| pKan-2UAG(V) | *kan^R^*-*2UAG(V)* | tRNAVal CUA | tRNAVal GAC |
| pKan-8UAG(K) | *kan^R^*-*8UAG(K)* | tRNALys CUA | tRNALys UUU |
| pKan-10UAG(F) | *kan^R^*-*10UAG(F)* | tRNAPhe CUA | tRNAPhe GAA |
| pKan-9UAG(F) | *kan^R^*-*9UAG(F)* | tRNAPhe CUA | tRNAPhe GAA |
| pKan-8UAG(F) | *kan^R^*-*8UAG(F)* | tRNAPhe CUA | tRNAPhe GAA |
| pKan-7UAG(F) | *kan^R^*-*7UAG(F)* | tRNAPhe CUA | tRNAPhe GAA |
| pKan-6UAG(F) | *kan^R^*-*6UAG(F)* | tRNAPhe CUA | tRNAPhe GAA |
| pKan-5UAG(F) | *kan^R^*-*5UAG(F)* | tRNAPhe CUA | tRNAPhe GAA |
| pKan-16UAG(D) | *kan^R^*-*16UAG(D)* | tRNAAsp CUA | tRNAAsp GUC |
| pKan-15UAG(D) | *kan^R^*-*15UAG(D)* | tRNAAsp CUA | tRNAAsp GUC |
| pKan-13UAG(D) | *kan^R^*-*13UAG(D)* | tRNAAsp CUA | tRNAAsp GUC |
| pKan-9UAG(D) | *kan^R^*-*9UAG(D)* | tRNAAsp CUA | tRNAAsp GUC |
| pKan-7UAG(D) | *kan^R^*-*7UAG(D)* | tRNAAsp CUA | tRNAAsp GUC |
| pKan-1UAG(3-MH) | *kan-1UAG(3-MH)* | pylRS and tRNAPyl CUA | none |
| pKan-2UAG(3-MH) | *kan-2UAG(3-MH)* | pylRS and tRNAPyl CUA | none |
| pKan-2UAG(3-MH) | *kan-2UAG(3-MH)* | pylRS and tRNAPyl CUA(G73T) | none |
| pKan-5UAG(3-MH) | *kan-5UAG(3-MH)* | pylRS and tRNApylPyl CUA | none |
| pKan-2UAG(5-HTP) | *kan-2UAG(3-HTP)* | WRS1 and ScWtRNA40A | none |
| pKan-3UAG(5-HTP) | *kan-3UAG(3-HTP)* | WRS1 and ScWtRNA40A | none |

| **Supplementary Table 8.** The sequences of *kan^R^*, *rfp*, *gfp*, and tagged *cm^R^* genes harboring UAG codons | |
| --- | --- |
| **Name** | **Sequence** |
| *kan^R^-10UAG(S)* | ATGTAGCATATTCAACGGGAAACGTAGTGCTAGAGGCCGCGATTAAATTAGAACATGGATGCTGATTTATATGGGTATAAATGGGCTCGCGATAATGTCGGGCAATAGGGTGCGACAATCTATCGATTGTATGGGAAGCCCGATGCGCCAGAGTTGTTTCTGAAACATGGCAAAGGTTAGGTTGCCAATGATGTTACAGATGAGATGGTCAGACTAAACTGGCTGACGGAATTTATGCCTCTTCCGACCATCAAGCATTTTATCCGTACTCCTGATGATGCATGGTTACTCACCACTGCGATCCCCGGGAAAACAGCATTCCAGGTATTAGAAGAATATCCTGATTAGGGTGAAAATATTGTTGATGCGCTGGCAGTGTTCCTGCGCCGGTTGCATTAGATTCCTGTTTGTAATTGTCCTTTTAACTAGGATCGCGTATTTCGTCTCGCTCAGGCGCAATAGCGAATGAATAACGGTTTGGTTGATGCGAGTGATTTTGATGACGAGCGTAATGGCTGGCCTGTTGAACAAGTCTGGAAAGAAATGCATAAACTTTTGCCATTCTCACCGGATTCAGTCGTCACTCATGGTGATTTCTCACTTGATAACCTTATTTTTGACGAGGGGAAATTAATAGGTTGTATTGATGTTGGACGAGTCGGAATCGCAGACCGATACCAGGATCTTGCCATCCTATGGAACTGCCTCGGTGAGTTTTCTCCTTCATTACAGAAACGGCTTTTTCAAAAATATGGTATTGATAATCCTGATATGAATAAATTGCAGTTTCATTTGATGCTCGATGAGTTTTTCTAA |
| *kan^R^-8UAG(I)* | ATGAGCCATTAGCAACGGGAAACGTCTTGCTCTAGGCCGCGATTAAATTCCAACATGGATGCTGATTTATATGGGTATAAATGGGCTCGCGATAATGTCGGGCAATCAGGTGCGACATAGTATCGATTGTATGGGAAGCCCGATGCGCCAGAGTTGTTTCTGAAACATGGCAAAGGTAGCGTTGCCAATGATGTTACAGATGAGATGGTCAGACTAAACTGGCTGACGGAATTTATGCCTCTTCCGACCTAGAAGCATTTTTAGCGTACTCCTGATGATGCATGGTTACTCACCACTGCGTAGCCCGGGAAAACAGCATTCCAGGTATTAGAAGAATATCCTGATTCAGGTGAAAATTAGGTTGATGCGCTGGCAGTGTTCCTGCGCCGGTTGCATTCGTAGCCTGTTTGTAATTGTCCTTTTAACAGCGATCGCGTATTTCGTCTCGCTCAGGCGCAATCACGAATGAATAACGGTTTGGTTGATGCGAGTGATTTTGATGACGAGCGTAATGGCTGGCCTGTTGAACAAGTCTGGAAAGAAATGCATAAACTTTTGCCATTCTCACCGGATTCAGTCGTCACTCATGGTGATTTCTCACTTGATAACCTTTAGTTTGACGAGGGGAAATTAATAGGTTGTATTGATGTTGGACGAGTCGGAATCGCAGACCGATACCAGGATCTTGCCATCCTATGGAACTGCCTCGGTGAGTTTTCTCCTTCATTACAGAAACGGCTTTTTCAAAAATATGGTATTGATAATCCTGATATGAATAAATTGCAGTTTCATTTGATGCTCGATGAGTTTTTCTAA |
| *kan^R^-5UAG(H)* | ATGAGCTAGATTCAACGGGAAACGTCTTGCTCTAGGCCGCGATTAAATTCCAACATGGATGCTGATTTATATGGGTATAAATGGGCTCGCGATAATGTCGGGCAATCAGGTGCGACAATCTATCGATTGTATGGGAAGCCCGATGCGCCAGAGTTGTTTCTGAAATAGGGCAAAGGTAGCGTTGCCAATGATGTTACAGATGAGATGGTCAGACTAAACTGGCTGACGGAATTTATGCCTCTTCCGACCATCAAGTAGTTTATCCGTACTCCTGATGATGCATGGTTACTCACCACTGCGATCCCCGGGAAAACAGCATTCCAGGTATTAGAAGAATATCCTGATTCAGGTGAAAATATTGTTGATGCGCTGGCAGTGTTCCTGCGCCGGTTGTAGTCGATTCCTGTTTGTAATTGTCCTTTTAACAGCGATCGCGTATTTCGTCTCGCTCAGGCGCAATCACGAATGAATAACGGTTTGGTTGATGCGAGTGATTTTGATGACGAGCGTAATGGCTGGCCTGTTGAACAAGTCTGGAAAGAAATGTAGAAACTTTTGCCATTCTCACCGGATTCAGTCGTCACTCATGGTGATTTCTCACTTGATAACCTTATTTTTGACGAGGGGAAATTAATAGGTTGTATTGATGTTGGACGAGTCGGAATCGCAGACCGATACCAGGATCTTGCCATCCTATGGAACTGCCTCGGTGAGTTTTCTCCTTCATTACAGAAACGGCTTTTTCAAAAATATGGTATTGATAATCCTGATATGAATAAATTGCAGTTTCATTTGATGCTCGATGAGTTTTTCTAA |
| *kan^R^-7UAG(Q)* | ATGAGCCATATTTAGCGGGAAACGTCTTGCTCTAGGCCGCGATTAAATTCCAACATGGATGCTGATTTATATGGGTATAAATGGGCTCGCGATAATGTCGGGTAGTCAGGTGCGACAATCTATCGATTGTATGGGAAGCCCGATGCGCCAGAGTTGTTTCTGAAACATGGCAAAGGTAGCGTTGCCAATGATGTTACAGATGAGATGGTCAGACTAAACTGGCTGACGGAATTTATGCCTCTTCCGACCATCAAGCATTTTATCCGTACTCCTGATGATGCATGGTTACTCACCACTGCGATCCCCGGGAAAACAGCATTCTAGGTATTAGAAGAATATCCTGATTCAGGTGAAAATATTGTTGATGCGCTGGCAGTGTTCCTGCGCCGGTTGCATTCGATTCCTGTTTGTAATTGTCCTTTTAACAGCGATCGCGTATTTCGTCTCGCTTAGGCGTAGTCACGAATGAATAACGGTTTGGTTGATGCGAGTGATTTTGATGACGAGCGTAATGGCTGGCCTGTTGAATAGGTCTGGAAAGAAATGCATAAACTTTTGCCATTCTCACCGGATTCAGTCGTCACTCATGGTGATTTCTCACTTGATAACCTTATTTTTGACGAGGGGAAATTAATAGGTTGTATTGATGTTGGACGAGTCGGAATCGCAGACCGATACTAGGATCTTGCCATCCTATGGAACTGCCTCGGTGAGTTTTCTCCTTCATTACAGAAACGGCTTTTTCAAAAATATGGTATTGATAATCCTGATATGAATAAATTGCAGTTTCATTTGATGCTCGATGAGTTTTTCTAA |
| *kan^R^-10UAG(R)* | ATGAGCCATATTCAATAGGAAACGTCTTGCTCTTAGCCGTAGTTAAATTCCAACATGGATGCTGATTTATATGGGTATAAATGGGCTTAGGATAATGTCGGGCAATCAGGTGCGACAATCTATTAGTTGTATGGGAAGCCCGATGCGCCAGAGTTGTTTCTGAAACATGGCAAAGGTAGCGTTGCCAATGATGTTACAGATGAGATGGTCTAGCTAAACTGGCTGACGGAATTTATGCCTCTTCCGACCATCAAGCATTTTATCTAGACTCCTGATGATGCATGGTTACTCACCACTGCGATCCCCGGGAAAACAGCATTCCAGGTATTAGAAGAATATCCTGATTCAGGTGAAAATATTGTTGATGCGCTGGCAGTGTTCCTGTAGCGGTTGCATTCGATTCCTGTTTGTAATTGTCCTTTTAACAGCGATTAGGTATTTCGTCTCGCTCAGGCGCAATCATAGATGAATAACGGTTTGGTTGATGCGAGTGATTTTGATGACGAGCGTAATGGCTGGCCTGTTGAACAAGTCTGGAAAGAAATGCATAAACTTTTGCCATTCTCACCGGATTCAGTCGTCACTCATGGTGATTTCTCACTTGATAACCTTATTTTTGACGAGGGGAAATTAATAGGTTGTATTGATGTTGGACGAGTCGGAATCGCAGACCGATACCAGGATCTTGCCATCCTATGGAACTGCCTCGGTGAGTTTTCTCCTTCATTACAGAAACGGCTTTTTCAAAAATATGGTATTGATAATCCTGATATGAATAAATTGCAGTTTCATTTGATGCTCGATGAGTTTTTCTAA |
| *kan^R^-8UAG(E)* | ATGAGCCATATTCAACGGTAGACGTCTTGCTCTAGGCCGCGATTAAATTCCAACATGGATGCTGATTTATATGGGTATAAATGGGCTCGCGATAATGTCGGGCAATCAGGTGCGACAATCTATCGATTGTATGGGAAGCCCGATGCGCCATAGTTGTTTCTGAAACATGGCAAAGGTAGCGTTGCCAATGATGTTACAGATTAGATGGTCAGACTAAACTGGCTGACGTAGTTTATGCCTCTTCCGACCATCAAGCATTTTATCCGTACTCCTGATGATGCATGGTTACTCACCACTGCGATCCCCGGGAAAACAGCATTCCAGGTATTATAGGAATATCCTGATTCAGGTTAGAATATTGTTGATGCGCTGGCAGTGTTCCTGCGCCGGTTGCATTCGATTCCTGTTTGTAATTGTCCTTTTAACAGCGATCGCGTATTTCGTCTCGCTCAGGCGCAATCACGAATGAATAACGGTTTGGTTGATGCGAGTGATTTTGATGACTAGCGTAATGGCTGGCCTGTTTAGCAAGTCTGGAAAGAAATGCATAAACTTTTGCCATTCTCACCGGATTCAGTCGTCACTCATGGTGATTTCTCACTTGATAACCTTATTTTTGACGAGGGGAAATTAATAGGTTGTATTGATGTTGGACGAGTCGGAATCGCAGACCGATACCAGGATCTTGCCATCCTATGGAACTGCCTCGGTGAGTTTTCTCCTTCATTACAGAAACGGCTTTTTCAAAAATATGGTATTGATAATCCTGATATGAATAAATTGCAGTTTCATTTGATGCTCGATGAGTTTTTCTAA |
| *kan^R^-7UAG(T)* | ATGAGCCATATTCAACGGGAATAGTCTTGCTCTAGGCCGCGATTAAATTCCAACATGGATGCTGATTTATATGGGTATAAATGGGCTCGCGATAATGTCGGGCAATCAGGTGCGTAGATCTATCGATTGTATGGGAAGCCCGATGCGCCAGAGTTGTTTCTGAAACATGGCAAAGGTAGCGTTGCCAATGATGTTTAGGATGAGATGGTCAGACTAAACTGGCTGTAGGAATTTATGCCTCTTCCGTAGATCAAGCATTTTATCCGTTAGCCTGATGATGCATGGTTACTCTAGACTGCGATCCCCGGGAAAACAGCATTCCAGGTATTAGAAGAATATCCTGATTCAGGTGAAAATATTGTTGATGCGCTGGCAGTGTTCCTGCGCCGGTTGCATTCGATTCCTGTTTGTAATTGTCCTTTTAACAGCGATCGCGTATTTCGTCTCGCTCAGGCGCAATCACGAATGAATAACGGTTTGGTTGATGCGAGTGATTTTGATGACGAGCGTAATGGCTGGCCTGTTGAACAAGTCTGGAAAGAAATGCATAAACTTTTGCCATTCTCACCGGATTCAGTCGTCACTCATGGTGATTTCTCACTTGATAACCTTATTTTTGACGAGGGGAAATTAATAGGTTGTATTGATGTTGGACGAGTCGGAATCGCAGACCGATACCAGGATCTTGCCATCCTATGGAACTGCCTCGGTGAGTTTTCTCCTTCATTACAGAAACGGCTTTTTCAAAAATATGGTATTGATAATCCTGATATGAATAAATTGCAGTTTCATTTGATGCTCGATGAGTTTTTCTAA |
| *kan^R^-4UAG(C)* | ATGAGCCATATTCAACGGGAAACGTCTTAGTCTAGGCCGCGATTAAATTCCAACATGGATGCTGATTTATATGGGTATAAATGGGCTCGCGATAATGTCGGGCAATCAGGTGCGACAATCTATCGATTGTATGGGAAGCCCGATGCGCCAGAGTTGTTTCTGAAACATGGCAAAGGTAGCGTTGCCAATGATGTTACAGATGAGATGGTCAGACTAAACTGGCTGACGGAATTTATGCCTCTTCCGACCATCAAGCATTTTATCCGTACTCCTGATGATGCATGGTTACTCACCACTGCGATCCCCGGGAAAACAGCATTCCAGGTATTAGAAGAATATCCTGATTCAGGTGAAAATATTGTTGATGCGCTGGCAGTGTTCCTGCGCCGGTTGCATTCGATTCCTGTTTAGAATTAGCCTTTTAACAGCGATCGCGTATTTCGTCTCGCTCAGGCGCAATCACGAATGAATAACGGTTTGGTTGATGCGAGTGATTTTGATGACGAGCGTAATGGCTGGCCTGTTGAACAAGTCTGGAAAGAAATGCATAAACTTTTGCCATTCTCACCGGATTCAGTCGTCACTCATGGTGATTTCTCACTTGATAACCTTATTTTTGACGAGGGGAAATTAATAGGTTAGATTGATGTTGGACGAGTCGGAATCGCAGACCGATACCAGGATCTTGCCATCCTATGGAACTGCCTCGGTGAGTTTTCTCCTTCATTACAGAAACGGCTTTTTCAAAAATATGGTATTGATAATCCTGATATGAATAAATTGCAGTTTCATTTGATGCTCGATGAGTTTTTCTAA |
| *kan^R^-10UAG(P)* | ATGAGCCATATTCAACGGGAAACGTCTTGCTCTAGGTAGCGATTAAATTCCAACATGGATGCTGATTTATATGGGTATAAATGGGCTCGCGATAATGTCGGGCAATCAGGTGCGACAATCTATCGATTGTATGGGAAGTAGGATGCGTAGGAGTTGTTTCTGAAACATGGCAAAGGTAGCGTTGCCAATGATGTTACAGATGAGATGGTCAGACTAAACTGGCTGACGGAATTTATGTAGCTTTAGACCATCAAGCATTTTATCCGTACTTAGGATGATGCATGGTTACTCACCACTGCGATCTAGGGGAAAACAGCATTCCAGGTATTAGAAGAATATTAGGATTCAGGTGAAAATATTGTTGATGCGCTGGCAGTGTTCCTGCGCCGGTTGCATTCGATTTAGGTTTGTAATTGTTAGTTTAACAGCGATCGCGTATTTCGTCTCGCTCAGGCGCAATCACGAATGAATAACGGTTTGGTTGATGCGAGTGATTTTGATGACGAGCGTAATGGCTGGCCTGTTGAACAAGTCTGGAAAGAAATGCATAAACTTTTGCCATTCTCACCGGATTCAGTCGTCACTCATGGTGATTTCTCACTTGATAACCTTATTTTTGACGAGGGGAAATTAATAGGTTGTATTGATGTTGGACGAGTCGGAATCGCAGACCGATACCAGGATCTTGCCATCCTATGGAACTGCCTCGGTGAGTTTTCTCCTTCATTACAGAAACGGCTTTTTCAAAAATATGGTATTGATAATCCTGATATGAATAAATTGCAGTTTCATTTGATGCTCGATGAGTTTTTCTAA |
| *kan^R^-20UAG(L)* | ATGAGCCATATTCAACGGGAAACGTCTTGCTCTAGGCCGCGATAGAATTCCAACATGGATGCTGATTAGTATGGGTATAAATGGGCTCGCGATAATGTCGGGCAATCAGGTGCGACAATCTATCGATAGTATGGGAAGCCCGATGCGCCAGAGTAGTTTTAGAAACATGGCAAAGGTAGCGTTGCCAATGATGTTACAGATGAGATGGTCAGATAGAACTGGTAGACGGAATTTATGCCTTAGCCGACCATCAAGCATTTTATCCGTACTCCTGATGATGCATGGTAGCTCACCACTGCGATCCCCGGGAAAACAGCATTCCAGGTATAGGAAGAATATCCTGATTCAGGTGAAAATATTGTTGATGCGTAGGCAGTGTTCTAGCGCCGGTAGCATTCGATTCCTGTTTGTAATTGTCCTTTTAACAGCGATCGCGTATTTCGTTAGGCTCAGGCGCAATCACGAATGAATAACGGTTAGGTTGATGCGAGTGATTTTGATGACGAGCGTAATGGCTGGCCTGTTGAACAAGTCTGGAAAGAAATGCATAAATAGTTGCCATTCTCACCGGATTCAGTCGTCACTCATGGTGATTTCTCATAGGATAACTAGATTTTTGACGAGGGGAAATAGATAGGTTGTATTGATGTTGGACGAGTCGGAATCGCAGACCGATACCAGGATTAGGCCATCCTATGGAACTGCCTCGGTGAGTTTTCTCCTTCATTACAGAAACGGCTTTTTCAAAAATATGGTATTGATAATCCTGATATGAATAAATTGCAGTTTCATTTGATGCTCGATGAGTTTTTCTAA |
| *kan^R^-10UAG(N)* | ATGAGCCATATTCAACGGGAAACGTCTTGCTCTAGGCCGCGATTATAGTCCTAGATGGATGCTGATTTATATGGGTATAAATGGGCTCGCGATTAGGTCGGGCAATCAGGTGCGACAATCTATCGATTGTATGGGAAGCCCGATGCGCCAGAGTTGTTTCTGAAACATGGCAAAGGTAGCGTTGCCTAGGATGTTACAGATGAGATGGTCAGACTATAGTGGCTGACGGAATTTATGCCTCTTCCGACCATCAAGCATTTTATCCGTACTCCTGATGATGCATGGTTACTCACCACTGCGATCCCCGGGAAAACAGCATTCCAGGTATTAGAAGAATATCCTGATTCAGGTGAATAGATTGTTGATGCGCTGGCAGTGTTCCTGCGCCGGTTGCATTCGATTCCTGTTTGTTAGTGTCCTTTTTAGAGCGATCGCGTATTTCGTCTCGCTCAGGCGCAATCACGAATGTAGAACGGTTTGGTTGATGCGAGTGATTTTGATGACGAGCGTTAGGGCTGGCCTGTTGAACAAGTCTGGAAAGAAATGCATAAACTTTTGCCATTCTCACCGGATTCAGTCGTCACTCATGGTGATTTCTCACTTGATAACCTTATTTTTGACGAGGGGAAATTAATAGGTTGTATTGATGTTGGACGAGTCGGAATCGCAGACCGATACCAGGATCTTGCCATCCTATGGAACTGCCTCGGTGAGTTTTCTCCTTCATTACAGAAACGGCTTTTTCAAAAATATGGTATTGATAATCCTGATATGAATAAATTGCAGTTTCATTTGATGCTCGATGAGTTTTTCTAA |
| *kan^R^-4UAG(M)* | ATGAGCCATATTCAACGGGAAACGTCTTGCTCTAGGCCGCGATTAAATTCCAACTAGGATGCTGATTTATATGGGTATAAATGGGCTCGCGATAATGTCGGGCAATCAGGTGCGACAATCTATCGATTGTATGGGAAGCCCGATGCGCCAGAGTTGTTTCTGAAACATGGCAAAGGTAGCGTTGCCAATGATGTTACAGATGAGTAGGTCAGACTAAACTGGCTGACGGAATTTTAGCCTCTTCCGACCATCAAGCATTTTATCCGTACTCCTGATGATGCATGGTTACTCACCACTGCGATCCCCGGGAAAACAGCATTCCAGGTATTAGAAGAATATCCTGATTCAGGTGAAAATATTGTTGATGCGCTGGCAGTGTTCCTGCGCCGGTTGCATTCGATTCCTGTTTGTAATTGTCCTTTTAACAGCGATCGCGTATTTCGTCTCGCTCAGGCGCAATCACGATAGAATAACGGTTTGGTTGATGCGAGTGATTTTGATGACGAGCGTAATGGCTGGCCTGTTGAACAAGTCTGGAAAGAAATGCATAAACTTTTGCCATTCTCACCGGATTCAGTCGTCACTCATGGTGATTTCTCACTTGATAACCTTATTTTTGACGAGGGGAAATTAATAGGTTGTATTGATGTTGGACGAGTCGGAATCGCAGACCGATACCAGGATCTTGCCATCCTATGGAACTGCCTCGGTGAGTTTTCTCCTTCATTACAGAAACGGCTTTTTCAAAAATATGGTATTGATAATCCTGATATGAATAAATTGCAGTTTCATTTGATGCTCGATGAGTTTTTCTAA |
| *kan^R^-5UAG(Y)* | ATGAGCCATATTCAACGGGAAACGTCTTGCTCTAGGCCGCGATTAAATTCCAACATGGATGCTGATTTATAGGGGTATAAATGGGCTCGCGATAATGTCGGGCAATCAGGTGCGACAATCTAGCGATTGTAGGGGAAGCCCGATGCGCCAGAGTTGTTTCTGAAACATGGCAAAGGTAGCGTTGCCAATGATGTTACAGATGAGATGGTCAGACTAAACTGGCTGACGGAATTTATGCCTCTTCCGACCATCAAGCATTTTATCCGTACTCCTGATGATGCATGGTTACTCACCACTGCGATCCCCGGGAAAACAGCATTCCAGGTATTAGAAGAATAGCCTGATTCAGGTGAAAATATTGTTGATGCGCTGGCAGTGTTCCTGCGCCGGTTGCATTCGATTCCTGTTTGTAATTGTCCTTTTAACAGCGATCGCGTATTTCGTCTCGCTCAGGCGCAATCACGAATGAATAACGGTTTGGTTGATGCGAGTGATTTTGATGACGAGCGTAATGGCTGGCCTGTTGAACAAGTCTGGAAAGAAATGCATAAACTTTTGCCATTCTCACCGGATTCAGTCGTCACTCATGGTGATTTCTCACTTGATAACCTTATTTTTGACGAGGGGAAATTAATAGGTTGTATTGATGTTGGACGAGTCGGAATCGCAGACCGATAGCAGGATCTTGCCATCCTATGGAACTGCCTCGGTGAGTTTTCTCCTTCATTACAGAAACGGCTTTTTCAAAAATATGGTATTGATAATCCTGATATGAATAAATTGCAGTTTCATTTGATGCTCGATGAGTTTTTCTAA |
| *kan^R^-10UAG(A)* | ATGAGCCATATTCAACGGGAAACGTCTTGCTCTAGGCCGCGATTAAATTCCAACATGGATTAGGATTTATATGGGTATAAATGGTAGCGCGATAATGTCGGGCAATCAGGTTAGACAATCTATCGATTGTATGGGAAGCCCGATTAGCCAGAGTTGTTTCTGAAACATGGCAAAGGTAGCGTTTAGAATGATGTTACAGATGAGATGGTCAGACTAAACTGGCTGACGGAATTTATGCCTCTTCCGACCATCAAGCATTTTATCCGTACTCCTGATGATTAGTGGTTACTCACCACTTAGATCCCCGGGAAAACATAGTTCCAGGTATTAGAAGAATATCCTGATTCAGGTGAAAATATTGTTGATTAGCTGTAGGTGTTCCTGCGCCGGTTGCATTCGATTCCTGTTTGTAATTGTCCTTTTAACAGCGATCGCGTATTTCGTCTCGCTCAGGCGCAATCACGAATGAATAACGGTTTGGTTGATGCGAGTGATTTTGATGACGAGCGTAATGGCTGGCCTGTTGAACAAGTCTGGAAAGAAATGCATAAACTTTTGCCATTCTCACCGGATTCAGTCGTCACTCATGGTGATTTCTCACTTGATAACCTTATTTTTGACGAGGGGAAATTAATAGGTTGTATTGATGTTGGACGAGTCGGAATCGCAGACCGATACCAGGATCTTGCCATCCTATGGAACTGCCTCGGTGAGTTTTCTCCTTCATTACAGAAACGGCTTTTTCAAAAATATGGTATTGATAATCCTGATATGAATAAATTGCAGTTTCATTTGATGCTCGATGAGTTTTTCTAA |
| *kan^R^-10UAG(G)* | ATGAGCCATATTCAACGGGAAACGTCTTGCTCTAGGCCGCGATTAAATTCCAACATGGATGCTGATTTATATTAGTATAAATGGGCTCGCGATAATGTCTAGCAATCATAGGCGACAATCTATCGATTGTATTAGAAGCCCGATGCGCCAGAGTTGTTTCTGAAACATTAGAAATAGAGCGTTGCCAATGATGTTACAGATGAGATGGTCAGACTAAACTGGCTGACGGAATTTATGCCTCTTCCGACCATCAAGCATTTTATCCGTACTCCTGATGATGCATGGTTACTCACCACTGCGATCCCCTAGAAAACAGCATTCCAGGTATTAGAAGAATATCCTGATTCATAGGAAAATATTGTTGATGCGCTGGCAGTGTTCCTGCGCCGGTTGCATTCGATTCCTGTTTGTAATTGTCCTTTTAACAGCGATCGCGTATTTCGTCTCGCTCAGGCGCAATCACGAATGAATAACTAGTTGGTTGATGCGAGTGATTTTGATGACGAGCGTAATTAGTGGCCTGTTGAACAAGTCTGGAAAGAAATGCATAAACTTTTGCCATTCTCACCGGATTCAGTCGTCACTCATGGTGATTTCTCACTTGATAACCTTATTTTTGACGAGGGGAAATTAATAGGTTGTATTGATGTTGGACGAGTCGGAATCGCAGACCGATACCAGGATCTTGCCATCCTATGGAACTGCCTCGGTGAGTTTTCTCCTTCATTACAGAAACGGCTTTTTCAAAAATATGGTATTGATAATCCTGATATGAATAAATTGCAGTTTCATTTGATGCTCGATGAGTTTTTCTAA |
| *kan^R^-10UAG(V)* | ATGAGCCATATTCAACGGGAAACGTCTTGCTCTAGGCCGCGATTAAATTCCAACATGGATGCTGATTTATATGGGTATAAATGGGCTCGCGATAATTAGGGGCAATCAGGTGCGACAATCTATCGATTGTATGGGAAGCCCGATGCGCCAGAGTTGTTTCTGAAACATGGCAAAGGTAGCTAGGCCAATGATTAGACAGATGAGATGTAGAGACTAAACTGGCTGACGGAATTTATGCCTCTTCCGACCATCAAGCATTTTATCCGTACTCCTGATGATGCATGGTTACTCACCACTGCGATCCCCGGGAAAACAGCATTCCAGTAGTTAGAAGAATATCCTGATTCAGGTGAAAATATTTAGGATGCGCTGGCATAGTTCCTGCGCCGGTTGCATTCGATTCCTTAGTGTAATTGTCCTTTTAACAGCGATCGCTAGTTTCGTCTCGCTCAGGCGCAATCACGAATGAATAACGGTTTGTAGGATGCGAGTGATTTTGATGACGAGCGTAATGGCTGGCCTGTTGAACAAGTCTGGAAAGAAATGCATAAACTTTTGCCATTCTCACCGGATTCAGTCGTCACTCATGGTGATTTCTCACTTGATAACCTTATTTTTGACGAGGGGAAATTAATAGGTTGTATTGATGTTGGACGAGTCGGAATCGCAGACCGATACCAGGATCTTGCCATCCTATGGAACTGCCTCGGTGAGTTTTCTCCTTCATTACAGAAACGGCTTTTTCAAAAATATGGTATTGATAATCCTGATATGAATAAATTGCAGTTTCATTTGATGCTCGATGAGTTTTTCTAA |
| *kan^R^-8UAG(K)* | ATGAGCCATATTCAACGGGAAACGTCTTGCTCTAGGCCGCGATTAAATTCCAACATGGATGCTGATTTATATGGGTATTAGTGGGCTCGCGATAATGTCGGGCAATCAGGTGCGACAATCTATCGATTGTATGGGTAGCCCGATGCGCCAGAGTTGTTTCTGTAGCATGGCTAGGGTAGCGTTGCCAATGATGTTACAGATGAGATGGTCAGACTAAACTGGCTGACGGAATTTATGCCTCTTCCGACCATCTAGCATTTTATCCGTACTCCTGATGATGCATGGTTACTCACCACTGCGATCCCCGGGTAGACAGCATTCCAGGTATTAGAAGAATATCCTGATTCAGGTGAAAATATTGTTGATGCGCTGGCAGTGTTCCTGCGCCGGTTGCATTCGATTCCTGTTTGTAATTGTCCTTTTAACAGCGATCGCGTATTTCGTCTCGCTCAGGCGCAATCACGAATGAATAACGGTTTGGTTGATGCGAGTGATTTTGATGACGAGCGTAATGGCTGGCCTGTTGAACAAGTCTGGTAGGAAATGCATTAGCTTTTGCCATTCTCACCGGATTCAGTCGTCACTCATGGTGATTTCTCACTTGATAACCTTATTTTTGACGAGGGGAAATTAATAGGTTGTATTGATGTTGGACGAGTCGGAATCGCAGACCGATACCAGGATCTTGCCATCCTATGGAACTGCCTCGGTGAGTTTTCTCCTTCATTACAGAAACGGCTTTTTCAAAAATATGGTATTGATAATCCTGATATGAATAAATTGCAGTTTCATTTGATGCTCGATGAGTTTTTCTAA |
| *kan^R^-10UAG(F)* | ATGAGCCATATTCAACGGGAAACGTCTTGCTCTAGGCCGCGATTAAATTCCAACATGGATGCTGATTTATATGGGTATAAATGGGCTCGCGATAATGTCGGGCAATCAGGTGCGACAATCTATCGATTGTATGGGAAGCCCGATGCGCCAGAGTTGTAGCTGAAACATGGCAAAGGTAGCGTTGCCAATGATGTTACAGATGAGATGGTCAGACTAAACTGGCTGACGGAATAGATGCCTCTTCCGACCATCAAGCATTAGATCCGTACTCCTGATGATGCATGGTTACTCACCACTGCGATCCCCGGGAAAACAGCATAGCAGGTATTAGAAGAATATCCTGATTCAGGTGAAAATATTGTTGATGCGCTGGCAGTGTAGCTGCGCCGGTTGCATTCGATTCCTGTTTGTAATTGTCCTTAGAACAGCGATCGCGTATAGCGTCTCGCTCAGGCGCAATCACGAATGAATAACGGTTTGGTTGATGCGAGTGATTAGGATGACGAGCGTAATGGCTGGCCTGTTGAACAAGTCTGGAAAGAAATGCATAAACTTTTGCCATAGTCACCGGATTCAGTCGTCACTCATGGTGATTAGTCACTTGATAACCTTATTTTTGACGAGGGGAAATTAATAGGTTGTATTGATGTTGGACGAGTCGGAATCGCAGACCGATACCAGGATCTTGCCATCCTATGGAACTGCCTCGGTGAGTTTTCTCCTTCATTACAGAAACGGCTTTTTCAAAAATATGGTATTGATAATCCTGATATGAATAAATTGCAGTTTCATTTGATGCTCGATGAGTTTTTCTAA |
| *kan^R^-16UAG(D)* | ATGAGCCATATTCAACGGGAAACGTCTTGCTCTAGGCCGCGATTAAATTCCAACATGTAGGCTTAGTTATATGGGTATAAATGGGCTCGCTAGAATGTCGGGCAATCAGGTGCGACAATCTATCGATTGTATGGGAAGCCCTAGGCGCCAGAGTTGTTTCTGAAACATGGCAAAGGTAGCGTTGCCAATTAGGTTACATAGGAGATGGTCAGACTAAACTGGCTGACGGAATTTATGCCTCTTCCGACCATCAAGCATTTTATCCGTACTCCTTAGTAGGCATGGTTACTCACCACTGCGATCCCCGGGAAAACAGCATTCCAGGTATTAGAAGAATATCCTTAGTCAGGTGAAAATATTGTTTAGGCGCTGGCAGTGTTCCTGCGCCGGTTGCATTCGATTCCTGTTTGTAATTGTCCTTTTAACAGCTAGCGCGTATTTCGTCTCGCTCAGGCGCAATCACGAATGAATAACGGTTTGGTTTAGGCGAGTTAGTTTTAGTAGGAGCGTAATGGCTGGCCTGTTGAACAAGTCTGGAAAGAAATGCATAAACTTTTGCCATTCTCACCGTAGTCAGTCGTCACTCATGGTGATTTCTCACTTGATAACCTTATTTTTGACGAGGGGAAATTAATAGGTTGTATTGATGTTGGACGAGTCGGAATCGCAGACCGATACCAGGATCTTGCCATCCTATGGAACTGCCTCGGTGAGTTTTCTCCTTCATTACAGAAACGGCTTTTTCAAAAATATGGTATTGATAATCCTGATATGAATAAATTGCAGTTTCATTTGATGCTCGATGAGTTTTTCTAA |
| *kan^R^-9UAG(F)* | ATGAGCCATATTCAACGGGAAACGTCTTGCTCTAGGCCGCGATTAAATTCCAACATGGATGCTGATTTATATGGGTATAAATGGGCTCGCGATAATGTCGGGCAATCAGGTGCGACAATCTATCGATTGTATGGGAAGCCCGATGCGCCAGAGTTGTAGCTGAAACATGGCAAAGGTAGCGTTGCCAATGATGTTACAGATGAGATGGTCAGACTAAACTGGCTGACGGAATAGATGCCTCTTCCGACCATCAAGCATTAGATCCGTACTCCTGATGATGCATGGTTACTCACCACTGCGATCCCCGGGAAAACAGCATAGCAGGTATTAGAAGAATATCCTGATTCAGGTGAAAATATTGTTGATGCGCTGGCAGTGTAGCTGCGCCGGTTGCATTCGATTCCTGTTTGTAATTGTCCTTAGAACAGCGATCGCGTATAGCGTCTCGCTCAGGCGCAATCACGAATGAATAACGGTTTGGTTGATGCGAGTGATTAGGATGACGAGCGTAATGGCTGGCCTGTTGAACAAGTCTGGAAAGAAATGCATAAACTTTTGCCATAGTCACCGGATTCAGTCGTCACTCATGGTGATTTCTCACTTGATAACCTTATTTTTGACGAGGGGAAATTAATAGGTTGTATTGATGTTGGACGAGTCGGAATCGCAGACCGATACCAGGATCTTGCCATCCTATGGAACTGCCTCGGTGAGTTTTCTCCTTCATTACAGAAACGGCTTTTTCAAAAATATGGTATTGATAATCCTGATATGAATAAATTGCAGTTTCATTTGATGCTCGATGAGTTTTTCTAA |
| *kan^R^-8UAG(F)* | ATGAGCCATATTCAACGGGAAACGTCTTGCTCTAGGCCGCGATTAAATTCCAACATGGATGCTGATTTATATGGGTATAAATGGGCTCGCGATAATGTCGGGCAATCAGGTGCGACAATCTATCGATTGTATGGGAAGCCCGATGCGCCAGAGTTGTAGCTGAAACATGGCAAAGGTAGCGTTGCCAATGATGTTACAGATGAGATGGTCAGACTAAACTGGCTGACGGAATAGATGCCTCTTCCGACCATCAAGCATTAGATCCGTACTCCTGATGATGCATGGTTACTCACCACTGCGATCCCCGGGAAAACAGCATAGCAGGTATTAGAAGAATATCCTGATTCAGGTGAAAATATTGTTGATGCGCTGGCAGTGTAGCTGCGCCGGTTGCATTCGATTCCTGTTTGTAATTGTCCTTAGAACAGCGATCGCGTATAGCGTCTCGCTCAGGCGCAATCACGAATGAATAACGGTTTGGTTGATGCGAGTGATTAGGATGACGAGCGTAATGGCTGGCCTGTTGAACAAGTCTGGAAAGAAATGCATAAACTTTTGCCATTCTCACCGGATTCAGTCGTCACTCATGGTGATTTCTCACTTGATAACCTTATTTTTGACGAGGGGAAATTAATAGGTTGTATTGATGTTGGACGAGTCGGAATCGCAGACCGATACCAGGATCTTGCCATCCTATGGAACTGCCTCGGTGAGTTTTCTCCTTCATTACAGAAACGGCTTTTTCAAAAATATGGTATTGATAATCCTGATATGAATAAATTGCAGTTTCATTTGATGCTCGATGAGTTTTTCTAA |
| *kan^R^-7UAG(F)* | ATGAGCCATATTCAACGGGAAACGTCTTGCTCTAGGCCGCGATTAAATTCCAACATGGATGCTGATTTATATGGGTATAAATGGGCTCGCGATAATGTCGGGCAATCAGGTGCGACAATCTATCGATTGTATGGGAAGCCCGATGCGCCAGAGTTGTAGCTGAAACATGGCAAAGGTAGCGTTGCCAATGATGTTACAGATGAGATGGTCAGACTAAACTGGCTGACGGAATAGATGCCTCTTCCGACCATCAAGCATTAGATCCGTACTCCTGATGATGCATGGTTACTCACCACTGCGATCCCCGGGAAAACAGCATAGCAGGTATTAGAAGAATATCCTGATTCAGGTGAAAATATTGTTGATGCGCTGGCAGTGTAGCTGCGCCGGTTGCATTCGATTCCTGTTTGTAATTGTCCTTAGAACAGCGATCGCGTATAGCGTCTCGCTCAGGCGCAATCACGAATGAATAACGGTTTGGTTGATGCGAGTGATTTTGATGACGAGCGTAATGGCTGGCCTGTTGAACAAGTCTGGAAAGAAATGCATAAACTTTTGCCATTCTCACCGGATTCAGTCGTCACTCATGGTGATTTCTCACTTGATAACCTTATTTTTGACGAGGGGAAATTAATAGGTTGTATTGATGTTGGACGAGTCGGAATCGCAGACCGATACCAGGATCTTGCCATCCTATGGAACTGCCTCGGTGAGTTTTCTCCTTCATTACAGAAACGGCTTTTTCAAAAATATGGTATTGATAATCCTGATATGAATAAATTGCAGTTTCATTTGATGCTCGATGAGTTTTTCTAA |
| *kan^R^-6UAG(F)* | ATGAGCCATATTCAACGGGAAACGTCTTGCTCTAGGCCGCGATTAAATTCCAACATGGATGCTGATTTATATGGGTATAAATGGGCTCGCGATAATGTCGGGCAATCAGGTGCGACAATCTATCGATTGTATGGGAAGCCCGATGCGCCAGAGTTGTAGCTGAAACATGGCAAAGGTAGCGTTGCCAATGATGTTACAGATGAGATGGTCAGACTAAACTGGCTGACGGAATAGATGCCTCTTCCGACCATCAAGCATTAGATCCGTACTCCTGATGATGCATGGTTACTCACCACTGCGATCCCCGGGAAAACAGCATAGCAGGTATTAGAAGAATATCCTGATTCAGGTGAAAATATTGTTGATGCGCTGGCAGTGTAGCTGCGCCGGTTGCATTCGATTCCTGTTTGTAATTGTCCTTAGAACAGCGATCGCGTATTTCGTCTCGCTCAGGCGCAATCACGAATGAATAACGGTTTGGTTGATGCGAGTGATTTTGATGACGAGCGTAATGGCTGGCCTGTTGAACAAGTCTGGAAAGAAATGCATAAACTTTTGCCATTCTCACCGGATTCAGTCGTCACTCATGGTGATTTCTCACTTGATAACCTTATTTTTGACGAGGGGAAATTAATAGGTTGTATTGATGTTGGACGAGTCGGAATCGCAGACCGATACCAGGATCTTGCCATCCTATGGAACTGCCTCGGTGAGTTTTCTCCTTCATTACAGAAACGGCTTTTTCAAAAATATGGTATTGATAATCCTGATATGAATAAATTGCAGTTTCATTTGATGCTCGATGAGTTTTTCTAA |
| *kan^R^-5UAG(F)* | ATGAGCCATATTCAACGGGAAACGTCTTGCTCTAGGCCGCGATTAAATTCCAACATGGATGCTGATTTATATGGGTATAAATGGGCTCGCGATAATGTCGGGCAATCAGGTGCGACAATCTATCGATTGTATGGGAAGCCCGATGCGCCAGAGTTGTAGCTGAAACATGGCAAAGGTAGCGTTGCCAATGATGTTACAGATGAGATGGTCAGACTAAACTGGCTGACGGAATAGATGCCTCTTCCGACCATCAAGCATTAGATCCGTACTCCTGATGATGCATGGTTACTCACCACTGCGATCCCCGGGAAAACAGCATAGCAGGTATTAGAAGAATATCCTGATTCAGGTGAAAATATTGTTGATGCGCTGGCAGTGTAGCTGCGCCGGTTGCATTCGATTCCTGTTTGTAATTGTCCTTTTAACAGCGATCGCGTATTTCGTCTCGCTCAGGCGCAATCACGAATGAATAACGGTTTGGTTGATGCGAGTGATTTTGATGACGAGCGTAATGGCTGGCCTGTTGAACAAGTCTGGAAAGAAATGCATAAACTTTTGCCATTCTCACCGGATTCAGTCGTCACTCATGGTGATTTCTCACTTGATAACCTTATTTTTGACGAGGGGAAATTAATAGGTTGTATTGATGTTGGACGAGTCGGAATCGCAGACCGATACCAGGATCTTGCCATCCTATGGAACTGCCTCGGTGAGTTTTCTCCTTCATTACAGAAACGGCTTTTTCAAAAATATGGTATTGATAATCCTGATATGAATAAATTGCAGTTTCATTTGATGCTCGATGAGTTTTTCTAA |
| *kan^R^-15UAG(D)* | ATGAGCCATATTCAACGGGAAACGTCTTGCTCTAGGCCGCGATTAAATTCCAACATGTAGGCTTAGTTATATGGGTATAAATGGGCTCGCTAGAATGTCGGGCAATCAGGTGCGACAATCTATCGATTGTATGGGAAGCCCTAGGCGCCAGAGTTGTTTCTGAAACATGGCAAAGGTAGCGTTGCCAATTAGGTTACATAGGAGATGGTCAGACTAAACTGGCTGACGGAATTTATGCCTCTTCCGACCATCAAGCATTTTATCCGTACTCCTTAGTAGGCATGGTTACTCACCACTGCGATCCCCGGGAAAACAGCATTCCAGGTATTAGAAGAATATCCTTAGTCAGGTGAAAATATTGTTTAGGCGCTGGCAGTGTTCCTGCGCCGGTTGCATTCGATTCCTGTTTGTAATTGTCCTTTTAACAGCTAGCGCGTATTTCGTCTCGCTCAGGCGCAATCACGAATGAATAACGGTTTGGTTTAGGCGAGTTAGTTTTAGTAGGAGCGTAATGGCTGGCCTGTTGAACAAGTCTGGAAAGAAATGCATAAACTTTTGCCATTCTCACCGGATTCAGTCGTCACTCATGGTGATTTCTCACTTGATAACCTTATTTTTGACGAGGGGAAATTAATAGGTTGTATTGATGTTGGACGAGTCGGAATCGCAGACCGATACCAGGATCTTGCCATCCTATGGAACTGCCTCGGTGAGTTTTCTCCTTCATTACAGAAACGGCTTTTTCAAAAATATGGTATTGATAATCCTGATATGAATAAATTGCAGTTTCATTTGATGCTCGATGAGTTTTTCTAA |
| *kan^R^-13UAG(D)* | ATGAGCCATATTCAACGGGAAACGTCTTGCTCTAGGCCGCGATTAAATTCCAACATGTAGGCTTAGTTATATGGGTATAAATGGGCTCGCTAGAATGTCGGGCAATCAGGTGCGACAATCTATCGATTGTATGGGAAGCCCTAGGCGCCAGAGTTGTTTCTGAAACATGGCAAAGGTAGCGTTGCCAATTAGGTTACATAGGAGATGGTCAGACTAAACTGGCTGACGGAATTTATGCCTCTTCCGACCATCAAGCATTTTATCCGTACTCCTTAGTAGGCATGGTTACTCACCACTGCGATCCCCGGGAAAACAGCATTCCAGGTATTAGAAGAATATCCTTAGTCAGGTGAAAATATTGTTTAGGCGCTGGCAGTGTTCCTGCGCCGGTTGCATTCGATTCCTGTTTGTAATTGTCCTTTTAACAGCTAGCGCGTATTTCGTCTCGCTCAGGCGCAATCACGAATGAATAACGGTTTGGTTTAGGCGAGTTAGTTTGATGACGAGCGTAATGGCTGGCCTGTTGAACAAGTCTGGAAAGAAATGCATAAACTTTTGCCATTCTCACCGGATTCAGTCGTCACTCATGGTGATTTCTCACTTGATAACCTTATTTTTGACGAGGGGAAATTAATAGGTTGTATTGATGTTGGACGAGTCGGAATCGCAGACCGATACCAGGATCTTGCCATCCTATGGAACTGCCTCGGTGAGTTTTCTCCTTCATTACAGAAACGGCTTTTTCAAAAATATGGTATTGATAATCCTGATATGAATAAATTGCAGTTTCATTTGATGCTCGATGAGTTTTTCTAA |
| *kan^R^-9UAG(D)* | ATGAGCCATATTCAACGGGAAACGTCTTGCTCTAGGCCGCGATTAAATTCCAACATGTAGGCTTAGTTATATGGGTATAAATGGGCTCGCTAGAATGTCGGGCAATCAGGTGCGACAATCTATCGATTGTATGGGAAGCCCTAGGCGCCAGAGTTGTTTCTGAAACATGGCAAAGGTAGCGTTGCCAATTAGGTTACATAGGAGATGGTCAGACTAAACTGGCTGACGGAATTTATGCCTCTTCCGACCATCAAGCATTTTATCCGTACTCCTTAGTAGGCATGGTTACTCACCACTGCGATCCCCGGGAAAACAGCATTCCAGGTATTAGAAGAATATCCTTAGTCAGGTGAAAATATTGTTGATGCGCTGGCAGTGTTCCTGCGCCGGTTGCATTCGATTCCTGTTTGTAATTGTCCTTTTAACAGCGATCGCGTATTTCGTCTCGCTCAGGCGCAATCACGAATGAATAACGGTTTGGTTGATGCGAGTGATTTTGATGACGAGCGTAATGGCTGGCCTGTTGAACAAGTCTGGAAAGAAATGCATAAACTTTTGCCATTCTCACCGGATTCAGTCGTCACTCATGGTGATTTCTCACTTGATAACCTTATTTTTGACGAGGGGAAATTAATAGGTTGTATTGATGTTGGACGAGTCGGAATCGCAGACCGATACCAGGATCTTGCCATCCTATGGAACTGCCTCGGTGAGTTTTCTCCTTCATTACAGAAACGGCTTTTTCAAAAATATGGTATTGATAATCCTGATATGAATAAATTGCAGTTTCATTTGATGCTCGATGAGTTTTTCTAA |
| *kan^R^-7UAG(D)* | ATGAGCCATATTCAACGGGAAACGTCTTGCTCTAGGCCGCGATTAAATTCCAACATGTAGGCTTAGTTATATGGGTATAAATGGGCTCGCTAGAATGTCGGGCAATCAGGTGCGACAATCTATCGATTGTATGGGAAGCCCTAGGCGCCAGAGTTGTTTCTGAAACATGGCAAAGGTAGCGTTGCCAATTAGGTTACATAGGAGATGGTCAGACTAAACTGGCTGACGGAATTTATGCCTCTTCCGACCATCAAGCATTTTATCCGTACTCCTTAGGATGCATGGTTACTCACCACTGCGATCCCCGGGAAAACAGCATTCCAGGTATTAGAAGAATATCCTGATTCAGGTGAAAATATTGTTGATGCGCTGGCAGTGTTCCTGCGCCGGTTGCATTCGATTCCTGTTTGTAATTGTCCTTTTAACAGCGATCGCGTATTTCGTCTCGCTCAGGCGCAATCACGAATGAATAACGGTTTGGTTGATGCGAGTGATTTTGATGACGAGCGTAATGGCTGGCCTGTTGAACAAGTCTGGAAAGAAATGCATAAACTTTTGCCATTCTCACCGGATTCAGTCGTCACTCATGGTGATTTCTCACTTGATAACCTTATTTTTGACGAGGGGAAATTAATAGGTTGTATTGATGTTGGACGAGTCGGAATCGCAGACCGATACCAGGATCTTGCCATCCTATGGAACTGCCTCGGTGAGTTTTCTCCTTCATTACAGAAACGGCTTTTTCAAAAATATGGTATTGATAATCCTGATATGAATAAATTGCAGTTTCATTTGATGCTCGATGAGTTTTTCTAA |
| *kan^R^*-*13UAG(R)* | ATGAGCCATATTCAATAGGAAACGTCTTGCTCTTAGCCGTAGTTAAATTCCAACATGGATGCTGATTTATATGGGTATAAATGGGCTTAGGATAATGTCGGGCAATCAGGTGCGACAATCTATTAGTTGTATGGGAAGCCCGATGCGCCAGAGTTGTTTCTGAAACATGGCAAAGGTAGCGTTGCCAATGATGTTACAGATGAGATGGTCTAGCTAAACTGGCTGACGGAATTTATGCCTCTTCCGACCATCAAGCATTTTATCTAGACTCCTGATGATGCATGGTTACTCACCACTGCGATCCCCGGGAAAACAGCATTCCAGGTATTAGAAGAATATCCTGATTCAGGTGAAAATATTGTTGATGCGCTGGCAGTGTTCCTGTAGCGGTTGCATTCGATTCCTGTTTGTAATTGTCCTTTTAACAGCGATTAGGTATTTTAGCTCGCTCAGGCGCAATCATAGATGAATAACGGTTTGGTTGATGCGAGTGATTTTGATGACGAGTAGAATGGCTGGCCTGTTGAACAAGTCTGGAAAGAAATGCATAAACTTTTGCCATTCTCACCGGATTCAGTCGTCACTCATGGTGATTTCTCACTTGATAACCTTATTTTTGACGAGGGGAAATTAATAGGTTGTATTGATGTTGGATAGGTCGGAATCGCAGACCGATACCAGGATCTTGCCATCCTATGGAACTGCCTCGGTGAGTTTTCTCCTTCATTACAGAAACGGCTTTTTCAAAAATATGGTATTGATAATCCTGATATGAATAAATTGCAGTTTCATTTGATGCTCGATGAGTTTTTCTAA |
| *kan^R^*-*5UAG(A)* | ATGAGCCATATTCAACGGGAAACGTCTTGCTCTAGGCCGCGATTAAATTCCAACATGGATTAGGATTTATATGGGTATAAATGGTAGCGCGATAATGTCGGGCAATCAGGTTAGACAATCTATCGATTGTATGGGAAGCCCGATTAGCCAGAGTTGTTTCTGAAACATGGCAAAGGTAGCGTTTAGAATGATGTTACAGATGAGATGGTCAGACTAAACTGGCTGACGGAATTTATGCCTCTTCCGACCATCAAGCATTTTATCCGTACTCCTGATGATGCATGGTTACTCACCACTGCGATCCCCGGGAAAACAGCATTCCAGGTATTAGAAGAATATCCTGATTCAGGTGAAAATATTGTTGATGCGCTGGCAGTGTTCCTGCGCCGGTTGCATTCGATTCCTGTTTGTAATTGTCCTTTTAACAGCGATCGCGTATTTCGTCTCGCTCAGGCGCAATCACGAATGAATAACGGTTTGGTTGATGCGAGTGATTTTGATGACGAGCGTAATGGCTGGCCTGTTGAACAAGTCTGGAAAGAAATGCATAAACTTTTGCCATTCTCACCGGATTCAGTCGTCACTCATGGTGATTTCTCACTTGATAACCTTATTTTTGACGAGGGGAAATTAATAGGTTGTATTGATGTTGGACGAGTCGGAATCGCAGACCGATACCAGGATCTTGCCATCCTATGGAACTGCCTCGGTGAGTTTTCTCCTTCATTACAGAAACGGCTTTTTCAAAAATATGGTATTGATAATCCTGATATGAATAAATTGCAGTTTCATTTGATGCTCGATGAGTTTTTCTAA |
| *kan^R^*-*3UAG(H)* | ATGAGCTAGATTCAACGGGAAACGTCTTGCTCTAGGCCGCGATTAAATTCCAACATGGATGCTGATTTATATGGGTATAAATGGGCTCGCGATAATGTCGGGCAATCAGGTGCGACAATCTATCGATTGTATGGGAAGCCCGATGCGCCAGAGTTGTTTCTGAAATAGGGCAAAGGTAGCGTTGCCAATGATGTTACAGATGAGATGGTCAGACTAAACTGGCTGACGGAATTTATGCCTCTTCCGACCATCAAGTAGTTTATCCGTACTCCTGATGATGCATGGTTACTCACCACTGCGATCCCCGGGAAAACAGCATTCCAGGTATTAGAAGAATATCCTGATTCAGGTGAAAATATTGTTGATGCGCTGGCAGTGTTCCTGCGCCGGTTGCATTCGATTCCTGTTTGTAATTGTCCTTTTAACAGCGATCGCGTATTTCGTCTCGCTCAGGCGCAATCACGAATGAATAACGGTTTGGTTGATGCGAGTGATTTTGATGACGAGCGTAATGGCTGGCCTGTTGAACAAGTCTGGAAAGAAATGCATAAACTTTTGCCATTCTCACCGGATTCAGTCGTCACTCATGGTGATTTCTCACTTGATAACCTTATTTTTGACGAGGGGAAATTAATAGGTTGTATTGATGTTGGACGAGTCGGAATCGCAGACCGATACCAGGATCTTGCCATCCTATGGAACTGCCTCGGTGAGTTTTCTCCTTCATTACAGAAACGGCTTTTTCAAAAATATGGTATTGATAATCCTGATATGAATAAATTGCAGTTTCATTTGATGCTCGATGAGTTTTTCTAA |
| *kan^R^*-*2UAG(I)* | ATGTAGGAACAAGATGGATTGCACGCAGGTTCTCCGGCCGCTTGGGTGGAGAGGCTATTCGGCTATGACTGGGCACAACAGACAATCGGCTGCTCTGATGCCGCCGTGTTCCGGCTGTCAGCGCAGGGGCGCCCGGTTCTTTTTGTCAAGACCGACCTGTCCGGTGCCCTGAATGAACTGCAGGACGAGGCAGCGCGGCTATCGTGGCTGGCCACGACGGGCGTTCCTTGCGCAGCTGTGCTCGACGTTGTCACTGAAGCGGGAAGGGACTGGCTGCTATTGGGCGAAGTGCCGGGGCAGGATCTCCTGTCATCTCACCTTGCTCCTGCCGAGAAAGTATCCATCATGGCTGATGCAATGCGGCGGCTGCATACGCTTGATCCGGCTACCTGCCCATTCGACCACCAAGCGAAACATCGCATCGAGCGAGCACGTACTCGGATGGAAGCCGGTCTTGTCGATCAGGATGATCTGGACGAAGAGCATCAGGGGCTCGCGCCAGCCGAACTGTTCGCCAGGCTCAAGGCGCGCATGCCCGACGGCGAGGATCTCGTCGTGACCCATGGCGATGCCTGCTTGCCGAATATCATGGTGGAAAATGGCCGCTTTTCTGGATTCATCGACTGTGGCCGGCTGGGTGTGGCGGACCGCTATCAGGACATAGCGTTGGCTACCCGTGATATTGCTGAAGAGCTTGGCGGCGAATGGGCTGACCGCTTCCTCGTGCTTTACGGTATCGCCGCTCCCGATTCGCAGCGCTAGGCCTTCTATCGCCTTCTTGACGAGTTCTTCTGA |
| *kan^R^*-*2UAG(M)* | ATGAGCCATATTCAACGGGAAACGTCTTGCTCTAGGCCGCGATTAAATTCCAACATGGATGCTGATTTATATGGGTATAAATGGGCTCGCGATAATGTCGGGCAATCAGGTGCGACAATCTATCGATTGTATGGGAAGCCCGATGCGCCAGAGTTGTTTCTGAAACATGGCAAAGGTAGCGTTGCCAATGATGTTACAGATGAGTAGGTCAGACTAAACTGGCTGACGGAATTTTAGCCTCTTCCGACCATCAAGCATTTTATCCGTACTCCTGATGATGCATGGTTACTCACCACTGCGATCCCCGGGAAAACAGCATTCCAGGTATTAGAAGAATATCCTGATTCAGGTGAAAATATTGTTGATGCGCTGGCAGTGTTCCTGCGCCGGTTGCATTCGATTCCTGTTTGTAATTGTCCTTTTAACAGCGATCGCGTATTTCGTCTCGCTCAGGCGCAATCACGAATGAATAACGGTTTGGTTGATGCGAGTGATTTTGATGACGAGCGTAATGGCTGGCCTGTTGAACAAGTCTGGAAAGAAATGCATAAACTTTTGCCATTCTCACCGGATTCAGTCGTCACTCATGGTGATTTCTCACTTGATAACCTTATTTTTGACGAGGGGAAATTAATAGGTTGTATTGATGTTGGACGAGTCGGAATCGCAGACCGATACCAGGATCTTGCCATCCTATGGAACTGCCTCGGTGAGTTTTCTCCTTCATTACAGAAACGGCTTTTTCAAAAATATGGTATTGATAATCCTGATATGAATAAATTGCAGTTTCATTTGATGCTCGATGAGTTTTTCTAA |
| *kan^R^*-*6UAG(N)* | ATGAGCCATATTCAACGGGAAACGTCTTGCTCTAGGCCGCGATTATAGTCCAACATGGATGCTGATTTATATGGGTATAAATGGGCTCGCGATTAGGTCGGGCAATCAGGTGCGACAATCTATCGATTGTATGGGAAGCCCGATGCGCCAGAGTTGTTTCTGAAACATGGCAAAGGTAGCGTTGCCTAGGATGTTACAGATGAGATGGTCAGACTATAGTGGCTGACGGAATTTATGCCTCTTCCGACCATCAAGCATTTTATCCGTACTCCTGATGATGCATGGTTACTCACCACTGCGATCCCCGGGAAAACAGCATTCCAGGTATTAGAAGAATATCCTGATTCAGGTGAATAGATTGTTGATGCGCTGGCAGTGTTCCTGCGCCGGTTGCATTCGATTCCTGTTTGTTAGTGTCCTTTTAACAGCGATCGCGTATTTCGTCTCGCTCAGGCGCAATCACGAATGAATAACGGTTTGGTTGATGCGAGTGATTTTGATGACGAGCGTAATGGCTGGCCTGTTGAACAAGTCTGGAAAGAAATGCATAAACTTTTGCCATTCTCACCGGATTCAGTCGTCACTCATGGTGATTTCTCACTTGATAACCTTATTTTTGACGAGGGGAAATTAATAGGTTGTATTGATGTTGGACGAGTCGGAATCGCAGACCGATACCAGGATCTTGCCATCCTATGGAACTGCCTCGGTGAGTTTTCTCCTTCATTACAGAAACGGCTTTTTCAAAAATATGGTATTGATAATCCTGATATGAATAAATTGCAGTTTCATTTGATGCTCGATGAGTTTTTCTAA |
| *kan^R^*-*9UAG(Q)* | ATGAGCCATATTCAACGGGAAACGTCTTGCTCTAGGCCGCGATTATAGTCCAACATGGATGCTGATTTATATGGGTATAAATGGGCTCGCGATTAGGTCGGGCAATCAGGTGCGACAATCTATCGATTGTATGGGAAGCCCGATGCGCCAGAGTTGTTTCTGAAACATGGCAAAGGTAGCGTTGCCTAGGATGTTACAGATGAGATGGTCAGACTATAGTGGCTGACGGAATTTATGCCTCTTCCGACCATCAAGCATTTTATCCGTACTCCTGATGATGCATGGTTACTCACCACTGCGATCCCCGGGAAAACAGCATTCCAGGTATTAGAAGAATATCCTGATTCAGGTGAATAGATTGTTGATGCGCTGGCAGTGTTCCTGCGCCGGTTGCATTCGATTCCTGTTTGTTAGTGTCCTTTTAACAGCGATCGCGTATTTCGTCTCGCTCAGGCGCAATCACGAATGAATAACGGTTTGGTTGATGCGAGTGATTTTGATGACGAGCGTAATGGCTGGCCTGTTGAACAAGTCTGGAAAGAAATGCATAAACTTTTGCCATTCTCACCGGATTCAGTCGTCACTCATGGTGATTTCTCACTTGATAACCTTATTTTTGACGAGGGGAAATTAATAGGTTGTATTGATGTTGGACGAGTCGGAATCGCAGACCGATACCAGGATCTTGCCATCCTATGGAACTGCCTCGGTGAGTTTTCTCCTTCATTACAGAAACGGCTTTTTCAAAAATATGGTATTGATAATCCTGATATGAATAAATTGCAGTTTCATTTGATGCTCGATGAGTTTTTCTAA |
| *kan^R^*-*7UAG(Y)* | ATGAGCCATATTCAACGGGAAACGTCTTGCTCTAGGCCGCGATTAAATTCCAACATGGATGCTGATTTATAGGGGTAGAAATGGGCTCGCGATAATGTCGGGCAATCAGGTGCGACAATCTAGCGATTGTAGGGGAAGCCCGATGCGCCAGAGTTGTTTCTGAAACATGGCAAAGGTAGCGTTGCCAATGATGTTACAGATGAGATGGTCAGACTAAACTGGCTGACGGAATTTATGCCTCTTCCGACCATCAAGCATTTTATCCGTACTCCTGATGATGCATGGTTACTCACCACTGCGATCCCCGGGAAAACAGCATTCCAGGTATTAGAAGAATAGCCTGATTCAGGTGAAAATATTGTTGATGCGCTGGCAGTGTTCCTGCGCCGGTTGCATTCGATTCCTGTTTGTAATTGTCCTTTTAACAGCGATCGCGTATTTCGTCTCGCTCAGGCGCAATCACGAATGAATAACGGTTTGGTTGATGCGAGTGATTTTGATGACGAGCGTAATGGCTGGCCTGTTGAACAAGTCTGGAAAGAAATGCATAAACTTTTGCCATTCTCACCGGATTCAGTCGTCACTCATGGTGATTTCTCACTTGATAACCTTATTTTTGACGAGGGGAAATTAATAGGTTGTATTGATGTTGGACGAGTCGGAATCGCAGACCGATAGCAGGATCTTGCCATCCTATGGAACTGCCTCGGTGAGTTTTCTCCTTCATTACAGAAACGGCTTTTTCAAAAATAGGGTATTGATAATCCTGATATGAATAAATTGCAGTTTCATTTGATGCTCGATGAGTTTTTCTAA |
| *kan^R^*-*2UAG(V)* | ATGATTGAACAAGATGGATTGCACGCAGGTTCTCCGGCCGCTTGGTAGGAGAGGCTATTCGGCTATGACTGGGCACAACAGACAATCGGCTGCTCTGATGCCGCCGTGTTCCGGCTGTCAGCGCAGGGGCGCCCGGTTCTTTTTGTCAAGACCGACCTGTCCGGTGCCCTGAATGAACTGCAGGACGAGGCAGCGCGGCTATCGTGGCTGGCCACGACGGGCGTTCCTTGCGCAGCTGTGCTCGACGTTGTCACTGAAGCGGGAAGGGACTGGCTGCTATTGGGCGAAGTGCCGGGGCAGGATCTCCTGTCATCTCACCTTGCTCCTGCCGAGAAATAGTCCATCATGGCTGATGCAATGCGGCGGCTGCATACGCTTGATCCGGCTACCTGCCCATTCGACCACCAAGCGAAACATCGCATCGAGCGAGCACGTACTCGGATGGAAGCCGGTCTTGTCGATCAGGATGATCTGGACGAAGAGCATCAGGGGCTCGCGCCAGCCGAACTGTTCGCCAGGCTCAAGGCGCGCATGCCCGACGGCGAGGATCTCGTCGTGACCCATGGCGATGCCTGCTTGCCGAATATCATGGTGGAAAATGGCCGCTTTTCTGGATTCATCGACTGTGGCCGGCTGGGTGTGGCGGACCGCTATCAGGACATAGCGTTGGCTACCCGTGATATTGCTGAAGAGCTTGGCGGCGAATGGGCTGACCGCTTCCTCTAGCTTTACGGTATCGCCGCTCCCGATTCGCAGCGCATCGCCTTCTATCGCCTTCTTGACGAGTTCTTCTGA |
| *cm^R^*-*1UAG* | ATGAACAAAGAAATTTAGGCTGTAGTTGAAGCCGTATCCAATGAAAAGGCGCTACCTCGCGAGAAGATTTTCGAAGCATTGGAAAGCGCGCTGGCGACAGCAGGTGGCGGAGGGTCAGGCGGTGGAGGGTCTGGAGGTGGCGGGTCAGAGAAAAAAATCACTGGATATACCACCGTTGATATATCCCAATGGCATCGTAAAGAACATTTTGAGGCATTTCAGTCAGTTGCTCAATGTACCTATAACCAGACCGTTCAGCTGGATATTACGGCCTTTTTAAAGACCGTAAAGAAAAATAAGCACAAGTTTTATCCGGCCTTTATTCACATTCTTGCCCGCCTGATGAATGCTCATCCGGAATTCCGTATGGCAATGAAAGACGGTGAGCTGGTGATATGGGATAGTGTTCACCCTTGTTACACCGTTTTCCATGAGCAAACTGAAACGTTTTCATCGCTTTGGAGTGAATACCACGACGATTTCCGGCAGTTTCTACACATATATTCGCAAGATGTGGCGTGTTACGGTGAAAACCTGGCCTATTTCCCTAAAGGGTTTATTGAGAATATGTTTTTCGTCTCAGCCAATCCCTGGGTGAGTTTCACCAGTTTTGATTTAAACGTGGCCAATATGGACAACTTCTTCGCCCCCGTTTTCACCATGGGCAAATATTATACGCAAGGCGACAAGGTGCTGATGCCGCTGGCGATTCAGGTTCATCATGCCGTCTGTGATGGCTTCCATGTCGGCAGAATGCTTAATGAATTACAACAGTACTGCGATGAGTGGCAGGGCGGGGCGTAA |
| *cm^R^*-*2UAG* | ATGAACAAAGAAATTTAGGCTGTAGTTGAATAGGTATCCAATGAAAAGGCGCTACCTCGCGAGAAGATTTTCGAAGCATTGGAAAGCGCGCTGGCGACAGCAGGTGGCGGAGGGTCAGGCGGTGGAGGGTCTGGAGGTGGCGGGTCAGAGAAAAAAATCACTGGATATACCACCGTTGATATATCCCAATGGCATCGTAAAGAACATTTTGAGGCATTTCAGTCAGTTGCTCAATGTACCTATAACCAGACCGTTCAGCTGGATATTACGGCCTTTTTAAAGACCGTAAAGAAAAATAAGCACAAGTTTTATCCGGCCTTTATTCACATTCTTGCCCGCCTGATGAATGCTCATCCGGAATTCCGTATGGCAATGAAAGACGGTGAGCTGGTGATATGGGATAGTGTTCACCCTTGTTACACCGTTTTCCATGAGCAAACTGAAACGTTTTCATCGCTTTGGAGTGAATACCACGACGATTTCCGGCAGTTTCTACACATATATTCGCAAGATGTGGCGTGTTACGGTGAAAACCTGGCCTATTTCCCTAAAGGGTTTATTGAGAATATGTTTTTCGTCTCAGCCAATCCCTGGGTGAGTTTCACCAGTTTTGATTTAAACGTGGCCAATATGGACAACTTCTTCGCCCCCGTTTTCACCATGGGCAAATATTATACGCAAGGCGACAAGGTGCTGATGCCGCTGGCGATTCAGGTTCATCATGCCGTCTGTGATGGCTTCCATGTCGGCAGAATGCTTAATGAATTACAACAGTACTGCGATGAGTGGCAGGGCGGGGCGTAA |
| *gfp-2UAG(W)* | ATGGTTAGCAAAGGTGAAGAACTGTTTACCGGCGTTGTGCCGATTCTGGTGGAACTGGATGGTGATGTGAATGGCCATAAATTTAGCGTTCGTGGCGAAGGCGAAGGTGATGCGACCAACGGTAAACTGACCCTGAAATTTATTTGCACCACCGGTAAACTGCCGGTTCCGTAGCCGACCCTGGTGACCACCCTGACCTATGGCGTTCAGTGCTTTAGCCGCTATCCGGATCATATGAAACGCCATGATTTCTTTAAAAGCGCGATGCCGGAAGGCTATGTGCAGGAACGTACCATTAGCTTCAAAGATGATGGCACCTATAAAACCCGTGCGGAAGTTAAATTTGAAGGCGATACCCTGGTGAACCGCATTGAACTGAAAGGTATTGATTTTAAAGAAGATGGCAACATTCTGGGTCATAAACTGGAATATAATTTCAACAGCCATTAGGTGTATATTACCGCCGATAAACAGAAAAATGGCATCAAAGCGAACTTTAAAATCCGTCACAACGTGGAAGATGGTAGCGTGCAGCTGGCGGATCATTATCAGCAGAATACCCCGATTGGTGATGGCCCGGTGCTGCTGCCGGATAATCATTATCTGAGCACCCAGAGCGTTCTGAGCAAAGATCCGAATGAAAAACGTGATCATATGGTGCTGCTGGAATTTGTTACCGCCGCGGGCATTACCCACGGTATGGATGAACTGTATAAAGGCAGCCACCATCATCATCACCATTAA |
| *rfp-3UAG(W)* | ATGTCAGTGATTAAGCAGGTAATGAAGACCAAGTTGCACCTTGAGGGCACTGTCAATGGCCATGATTTTACGATCGAGGGTAAAGGTGAAGGCAAGCCGTACGAAGGGTTACAGCACATGAAAATGACAGTCACCAAAGGCGCGCCTCTGCCGTTTTCCGTTCATATTCTTACACCTAGCCACATGTATGGAAGCAAACCGTTTAATAAGTATCCAGCGGATATCCCAGACTACCACAAACAGTCTTTTCCCGAAGGTATGTCTTAGGAGCGGTCGATGATTTTTGAAGATGGTGGCGTATGCACCGCCAGTAATCATTCCAGCATAAACTTGCAAGAGAACTGTTTCATCTATGATGTTAAATTTCATGGTGTGAACCTGCCTCCGGATGGGCCCGTAATGCAAAAAACCATTGCTGGATAGGAGCCGAGCGTGGAAACACTGTACGTGCGTGACGGGATGTTAAAAAGTGACACTGCAATGGTTTTTAAACTGAAAGGAGGCGGTCATCATCGTGTTGATTTCAAAACGACGTATAAAGCCAAAAAACCTGTCAAGCTGCCAGAATTTCATTTCGTTGAACATCGCCTGGAACTGACCAAACACGATAAAGATTTCACAACTTAGGACCAGCAGGAGGCAGCCGAAGGCCATTTCTCACCGCTGCCGAAGGCTCTCCCATAA |

| **Supplementary Table 9.** The gene sequences of the tRNAs that decode the UAG codon | |
| --- | --- |
| **Name** | **Sequence (5’ – 3’)** |
| *E. coli* tRNATrp CUA | AGGGGCGTAGTTCAATTGGTAGAGCACCGGTCTCTAAAACCGGGTGTTGGGAGTTCGAGTCTCTCCGCCCCTGCCA |
| *E. coli* tRNASer CUA | GGTGAGGTGGCCGAGAGGCTGAAGGCGCTCCCCTCTAAAGGGAGTATGCGGTCAAAAGCTGCATCCGGGGTTCGAATCCCCGCCTCACCGCCA |
| *E. coli* tRNAHis CUA | GTGGCTATAGCTCAGTTGGTAGAGCCCTGGATTCTAATTCCAGTTGTCGTGGGTTCGAATCCCATTAGCCACCCCA |
| *E. coli* tRNAIle CUA | AGGCTTGTAGCTCAGGTGGTTAGAGCGCACCCCTCTAAAGGGTGAGGTCGGTGGTTCAAGTCCACTCAGGCCTACCA |
| *E. coli* tRNAGln CUA | TGGGGTATCGCCAAGCGGTAAGGCACCGGATTCTAATTCCGGCATTCCGAGGTTCGAATCCTCGTACCCCAGCCA |
| *E. coli* tRNAArg CUA | GCATCCGTAGCTCAGCTGGATAGAGTACTCGGCTCTAAACCGAGCGGTCGGAGGTTCGAATCCTCCCGGATGCACCA |
| *E. coli* tRNAGlu CUA | GTCCCCTTCGTCTAGAGGCCCAGGACACCGCCCTCTAACGGCGGTAACAGGGGTTCGAATCCCCTAGGGGACGCCA |
| *E. coli* tRNAThr CUA | GCTGATATAGCTCAGTTGGTAGAGCGCACCCTTCTAAAGGGTGAGGtCGGCAGTTCGAATCTGCCTATCAGCACCA |
| *E. coli* tRNACys CUA | GGCGCGTTAACAAAGCGGTTATGTAGCGGATTCTAAATCCGTCTAGTCCGGTTCGACTCCGGAACGCGCCTCCA |
| *E. coli* tRNAPro CUA | CGGTGATTGGCGCAGCCtGGTAGCGCACTTCGTTCTAGACGAAGGGGtCGGAGGTTCGAATCCTCTATCACCGACCA |
| *E. coli* tRNALeu CUA | GCGAAGGTGGCGGAATTGGTAGACGCGCTAGCTTCTAGTGTTAGTGTCCTTACGGACGTGGGGGTTCAAGTCCCCCCCCTCGCACCA |
| *E. coli* tRNAAsn CUA | TCCTCTGTAGTTCAGTCGGTAGAACGGCGGACTCTAAATCCGTATGTCACTGGTTCGAGTCCAGTCAGAGGAGCCA |
| *E. coli* tRNAMet CUA | GGCTACGTAGCTCAGTTGGTtAGAGCACATCACTCTAAATGATGGGGTCACAGGTTCGAATCCCGTCGTAGCCACCA |
| *E. coli* tRNATyr CUA | GGTGGGGTTCCCGAGCGGCCAAAGGGAGCAGACTCTAAATCTGCCGTCATCGACTTCGAAGGTTCGAATCCTTCCCCCACCACCA |
| *E. coli* tRNAAla CUA | GGGGCTATAGCTCAGCTGGGAGAGCGCTTGCATCTAATGCAAGAGGtCAGCGGTTCGATCCCGCTTAGCTCCACCA |
| *E. coli* tRNAGly CUA | GGAATAGCTCAGTTGGTAGAGCACGACCTTCTAAAGGTCGGGGTCGCGAGTTCGAGTCTCGTTTCCCGC |
| *E. coli* tRNAVal CUA | GCGTCCGTAGCTCAGTTGGTtAGAGCACCACCTTCTAATGGTGGGGGtCGGTGGTTCGAGTCCACTCGGACGCACCA |
| *E. coli* tRNALys CUA | GGGTCGTTAGCTCAGTTGGTAGAGCAGTTGACTCTAAATCAATTGGtCGCAGGTTCGAATCCTGCACGACCCACCA |
| *E. coli* tRNAPhe CUA | GCCCGGATAGCTCAGTCGGTAGAGCAGGGGATTCTAAATCCCCGTGTCCTTGGTTCGATTCCGAGTCCGGGCACCA |
| *E. coli* tRNAAsp CUA | GGAGCGGTAGTTCAGTCGGTTAGAATACCTGCCTCTAACGCAGGGGGTCGCGGGTTCGAGTCCCGTCCGTTCCGCCA |
| *C. glutamicum* tRNALeu CUA | GCCCTTGTAGCCCAATTGGCAGAGGCAACGGATTCTAAACCCGTCCAGTGTGAGTTCGAGTCTCACCAGGGGCACCA |
| *C. glutamicum*  tRNAGly CUA | GCGGATGTAGCGCAGTTGGTAGCGCATCACCTTCTAAAGGTGAGGGTCGCGAGTTCGAGTCTCGTCATCCGCT |
